# Sequencing of the *Prnp–Prnd* deletion junction in *Prn*^-/-^ mouse tissues after formaldehyde fixation and >22 years at room temperature

**DOI:** 10.64898/2026.11.10.765550

**Authors:** Michele Biase, Adriano Aguzzi

## Abstract

In 2004, Nicolas Genoud and colleagues reported that deletion of *Prnd*, which encodes Doppel, prevented neurodegeneration in mice with extensive *Prnp* deletions. The work tested the proposal that abnormal Doppel expression in the brain caused cerebellar degeneration in mice lacking the cellular prion protein, PrP. Advances in automated image analysis have expanded scrutiny of published figures, and post-publication reviews have raised concerns about apparent figure manipulation in this work, calling its authenticity and reliability into question. Here we assemble the surviving contemporaneous records of the original work and report new experiments on archived tissue. To reassess the reported genetic rearrangement, DNA from archived paraffin-embedded brain tissue was amplified across the putative deletion junction and analyzed by bidirectional Sanger sequencing and comparison with mouse reference sequences. The two reads spanned 607 positions and placed *Prnp*-side sequence, a 34-base loxP site and *Prnd*-side sequence in the expected order, with 93 intervening bases that matched bacteriophage P1 loxP-flanking and plasmid cloning-site sequences rather than the mouse reference and therefore probably derive from the targeting construct. Independent de novo reassembly from the 2025 Sanger chromatogram files reproduced the junction sequence; the assemblies differed only at three terminal bases with low base-call quality. These findings support the expected local junction arrangement and provide a publicly inspectable sequence description that extends the molecular information presented in the original paper. The amplicon size and the position of the loxP site match the values stated in the original paper, as does the approximately 18-kb distance between the two flanking sequences in the wild-type genome, which the deletion brings together. Recovery of junction DNA after more than two decades of room-temperature storage as paraformaldehyde-fixed, paraffin-embedded tissue demonstrates the value of retained material for reassessing historical experimental claims.

## Introduction

The 2004 study by Genoud and colleagues from the laboratory of one of us (A.A.) addressed why some mice lacking the cellular prion protein, PrP, developed cerebellar degeneration whereas other PrP-deficient lines remained healthy.^1^ The *Prnp* gene encodes PrP; its neighbouring gene, *Prnd*, encodes the related protein Doppel, also called *Dpl*. Extensive *Prnp* deletions caused abnormal Doppel expression in the brain, and earlier work linked increased Doppel levels to loss of cerebellar Purkinje neurons and impaired movement.^3,5^ Deleting *Prnd* together with *Prnp* tested whether this degeneration arose from the absence of PrP itself or from abnormal Doppel expression. The reported absence of cerebellar degeneration in the double-deficient mice supported a causal role for Doppel in this model.^1^ Reassessing the genetic evidence for the deletion is therefore relevant to a central mechanistic conclusion of the original study.

Advances in automated image-comparison tools have made systematic screening of scientific figures increasingly feasible.^18^ Post-publication review sites raised concerns about Figure 1B of the original paper.^17^ Two lanes labelled “Som” and “H₂O” appeared similar, and a sharp change in the image background was visible between them. “Som” denoted DNA from somatic tissue, specifically liver, whereas “H₂O” denoted a water-only PCR control containing no tissue DNA. The concern was that lanes presented as separate controls might share image content or have been joined without disclosure. A.A. reported the matter to PNAS on 25 March 2024, with the research-integrity officer of the University of Zurich in copy. The same altered panel appears in Genoud’s doctoral thesis,^22^ establishing that the alteration predates submission of the manuscript. These concerns raised substantial questions about the authenticity and reliability of the reported evidence and warranted an extended investigation.

**Figure 1.**
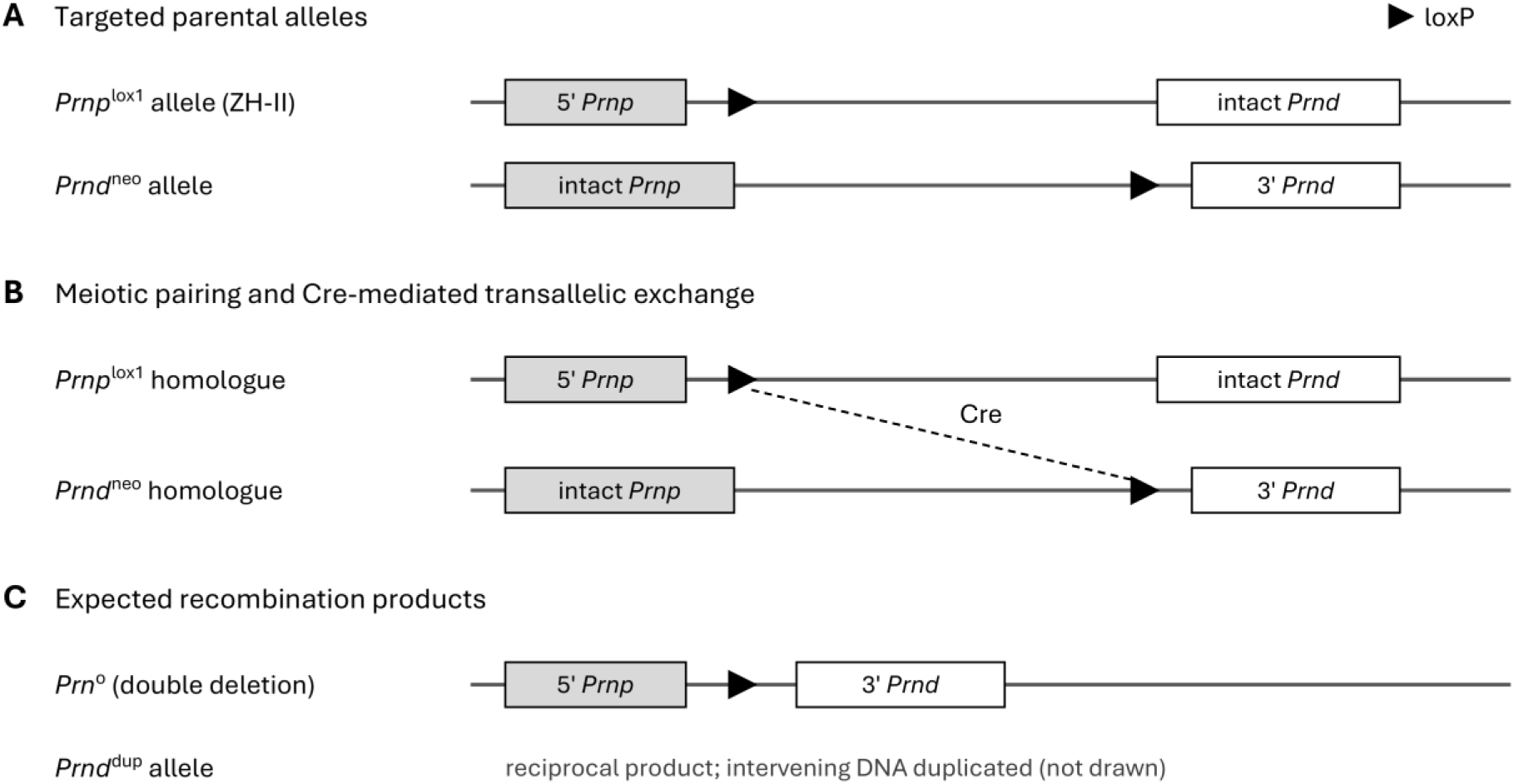
Genetic strategy for producing a chromosome lacking *Prnp* and *Prnd* by transallelic meiotic recombination (TAMERE). (A) One copy of chromosome 2 carries the *Prnp ^lox1^* allele, including a retained 5′ *Prnp* region and a loxP site, together with intact *Prnd*. The other carries intact *Prnp* and the *Prnd ^neo^* allele, including the second exchange site and retained 3′ *Prnd* sequence. (B) Cre-mediated exchange between loxP sites on these opposite chromosome copies during sperm formation is expected to generate (C) the *Prn °* allele lacking both genes and the reciprocal *Prnd ^dup^* product with duplicated intervening DNA. The drawing omits the selection cassette and its additional loxP site for clarity and is not to scale. The archival assay examined the inherited junction in brain tissue from later descendants; its local sequence arrangement is shown in Figure 3. Adapted from Fig. 1A of Genoud et al.¹.

The genetic strategy underlying the original Figure 1B used Cre, an enzyme that recombines DNA at short recognition sequences called loxP sites. The breeding males carried loxP sites near *Prnp* and *Prnd* on opposite members of chromosome pair 2; these animals were termed “transloxing” males. Cre expression driven by the Sycp1 promoter (the promoter of the gene encoding synaptonemal complex protein 1, which is active in meiotic germ cells) was used to induce exchange between these sites during meiosis, the cell division that produces sperm. The intended product was a chromosome lacking both genes, which could then be inherited by offspring. This deleted allele is designated *Prn°* (Figure 1); mice carrying it on both chromosomes are written *Prn*^o/o^ or, equivalently, *Prn*^−/−^, and heterozygous carriers *Prn*^+/o^. Mice lacking only *Prnp* are written *Prnp*^−/−^ or, as in the laboratory records, Prp^−/−^ or PrP^−/−^. This approach was called transallelic meiotic recombination, or TAMERE.^19^ The original Figure 1B used PCR to test for the resulting junction in testis DNA from males with or without the Cre transgene, with liver DNA as a control for somatic cells, meaning cells outside the reproductive lineage. The original paper reported that primers D and E, whose annealing sites lie approximately 18 kb apart in the wild-type locus, amplify a 600-bp product only from the recombined allele; that 1 of 482 pups sired by transloxing males carried the deletion; and that the breakpoint had been sequenced with the same primers, without showing the sequence.^1^

The retained paraffin archive allowed investigation of the reported rearrangement in surviving brain tissue from descendants of the original colony. DNA extraction, PCR and sequencing were used to examine the local order of *Prnp* sequence, loxP and *Prnd* sequence.

## Results

### Archival material, chronology and recovery of junction DNA

One of us (A.A.) has retained a complete archive of the paraffin-embedded material produced since his arrival in Zurich in 1986, including the blocks from all mice in this study; none of these materials has been destroyed. The blocks are catalogued in the “histobook” of the Institute of Neuropathology, an internal record shared by all its staff in which the provenance and experimental details of every block of fixed, paraffin-embedded tissue are entered. Originally kept as a bound physical book, and later as a Microsoft Access database, the histobook has recorded more than 75,000 paraffin blocks and identifies each block by a number with the prefix AG-. The blocks studied here were located through the histobook.

The histobook entries also constrain the approximate chronology of the specimens. A printout of histobook entries, scanned into the investigation report of 26 November 2025 (Supplementary Fig. S1), records AG-28112 as a *Prn*^−/−^ brain from a 61-week-old animal and AG-27374 as a *Prn*^−/−^ brain, both with N. Genoud as responsible scientist; the consecutively numbered entries on the same page (AG-27375 to AG-27381), made by another scientist, give 6 May 2003 as the date of death. AG-27371 to AG-27374 are listed among the histobook entries made by N. Genoud as brain specimens recorded as wild type, Prp^−/−^, Lox1^+/−^ (presumably heterozygous for the *Prnp^lox^*^1^ allele; Figure 1A) and *Prn*^−/−^, respectively, all entered on 28 May 2003; AG-28112 is absent from the export of his entries (Genoud_HistobookMain.xls), although the printout lists it under his name (Supplementary Fig. S1); its number falls between export entries made on 28 May and 12 November 2003 (Supplementary Table S1). An export of N. Genoud’s entries (Genoud_HistobookMain.xls) comprises 333 records created between 20 July 2001 and 15 November 2005 for which he is recorded as the responsible scientist, alone or with colleagues. AG-27374 is the only record in it entered as *Prn*^−/−^; the other Doppel-related entries concern *Dpl*-deficient males and their littermates sampled for testis and epididymis in 2001, skeletal muscle from mice expressing a Doppel transgene in muscle (HSA-*Dpl*) in 2003–2004, and *Prnd*^−/−^ males examined for fertility in 2005 (Supplementary Table S2). The archival material examined here had therefore been kept at room temperature for more than 22 years. Protein, cell and tissue homogenates were removed from deep-freezer storage after several years and are no longer available.

Breeding was difficult because of the male infertility reported for the double-null mice in the original paper.^1^ Breeding was eventually discontinued for economic reasons and because the laboratory had stopped working on Doppel (*Dpl*). Consequently, no living mice from this colony are now available for experimentation. The line had, however, been deposited with the European Mouse Mutant Archive (EMMA), where it is listed as strain EM:02432, B6.129P2-*Prnp/Prnd*^tm1Aag^/Cnrm (MGI allele MGI:3047634), backcrossed for seven generations to C57BL/6J, with male infertility as its recorded phenotype, and available to order;^20,21^. The Whitehead Institute (Cambridge, MA, USA) is recorded as an additional owner of the strain. Any laboratory can therefore obtain the line and verify the deletion independently.

Contemporaneous correspondence places work on generating the deletion line earlier than publication. In an email to Charles Weissmann dated 10 June 2002, A.A. reported a first live candidate carrying the intended *Prnp*–*Prnd* deletion, detected by PCR, with Southern confirmation still pending. The same message states that obtaining this mouse had taken a year and that recombination had previously been detected in sperm, indicating that the effort began around 2001. A progress report of 12 July 2001 already laid out the TAMERE breeding scheme, and on 17 June 2002 A.A. gave the yield as one positive among 614 screened mice, with the loxP sites about 20 kb apart; the manuscript drafts, the thesis and the published paper give 1 of 482 pups sired by transloxing males. The generation and initial screening of the line therefore date back more than 24 years.

On 8 October 2003, Nicolas Genoud reported that the Northern analyses, sequencing and spleen Doppel Western blotting were complete and that the manuscript was in PNAS format; he also reported that Southern blotting had again failed, adding that ‘we have Northern, Western and sequencing which all show that the locus has been depleted’. The original Figure 1B experiment itself, PCR on testis DNA of transloxing males, corresponds to the germ-cell recombination that the June 2002 message describes as long established; the panel is described in a manuscript draft dated 10 June 2003, with figure source files dated June 2003, and is shown in the September 2003 progress report. Hence, it was acquired before June 2003. The original paper appeared in the 23 March 2004 issue of PNAS and records receipt for review on 25 October 2003.^1^ The underlying experiments therefore predated publication, but the exact date of the original Figure 1B experiment and the collection dates of the sequenced specimens remain unknown.

Genoud’s physical laboratory notebooks could not be located in the institute archives in 2024, and the original 2004 gel and sequence traces have not been found. The records that were retrieved comprise his dated progress-report presentations from 2001 to 2004, the manuscript drafts of June and December 2003, his doctoral thesis,^22^ the laboratory’s correspondence with him (623 messages from 1999 onwards), the colony database and animal register, and the paraffin blocks. Apart from the number of screened offspring (614 in the June 2002 email versus 482 pups in the drafts, thesis and paper), these records agree with the published description of the experiment, the primers, the product size and the screening outcome. The records do not explain the difference in the number of screened offspring; one possibility is that the 614 mice screened by June 2002 included offspring of matings other than those of transloxing males, which the published denominator excludes. The September 2003 progress report gives the frequency of the deleted allele as 0.2%, which corresponds to 1 of 482 (0.21%) rather than 1 of 614 (0.16%); either figure indicates that the deletion arose rarely.

Genomic DNA was extracted from paraffin blocks of four animals recorded in the laboratory database as wild type (AG-27371), *Prp*^−/−^ (AG-27372) and *Prn*^−/−^ (AG-27374 and AG-28112). The corresponding DNA concentrations were 123.2, 149.1, 270.3 and 93.1 ng/µL, respectively. Initial extraction attempts in July 2025 had yielded DNA of insufficient quality for PCR, so additional sections were cut and the extraction was repeated with more material. On a 0.5% agarose gel, all four extracts appeared as continuous smears of fragmented DNA extending below the smallest band of the 1-kb ladder, without a discrete high-molecular-weight band (Supplementary Fig. S2).

PCR of DNA recovered from archived paraffin-embedded tissue yielded a band migrating at approximately 600 bp in sample AG-28112 (lane 112), the size predicted for the recombined allele in the original paper. A weaker band was visible in sample AG-27374 (lane 74) at higher exposure, whereas no corresponding band was detected in the wild-type control AG-27371 (lane 71) or in the Prp^−/−^ sample AG-27372 (lane 72). Additional bands migrating near 100 bp were present in all samples, including the wild-type control, and yielded no interpretable sequence. The wild-type and Prp^−/−^ samples processed in parallel are therefore the only negative controls for this reaction. Figure 2 reproduces two exposures that the 2025 laboratory report identifies as the same gel; band sizes are the report’s estimates from migration relative to the molecular-weight markers.

**Figure 2.**
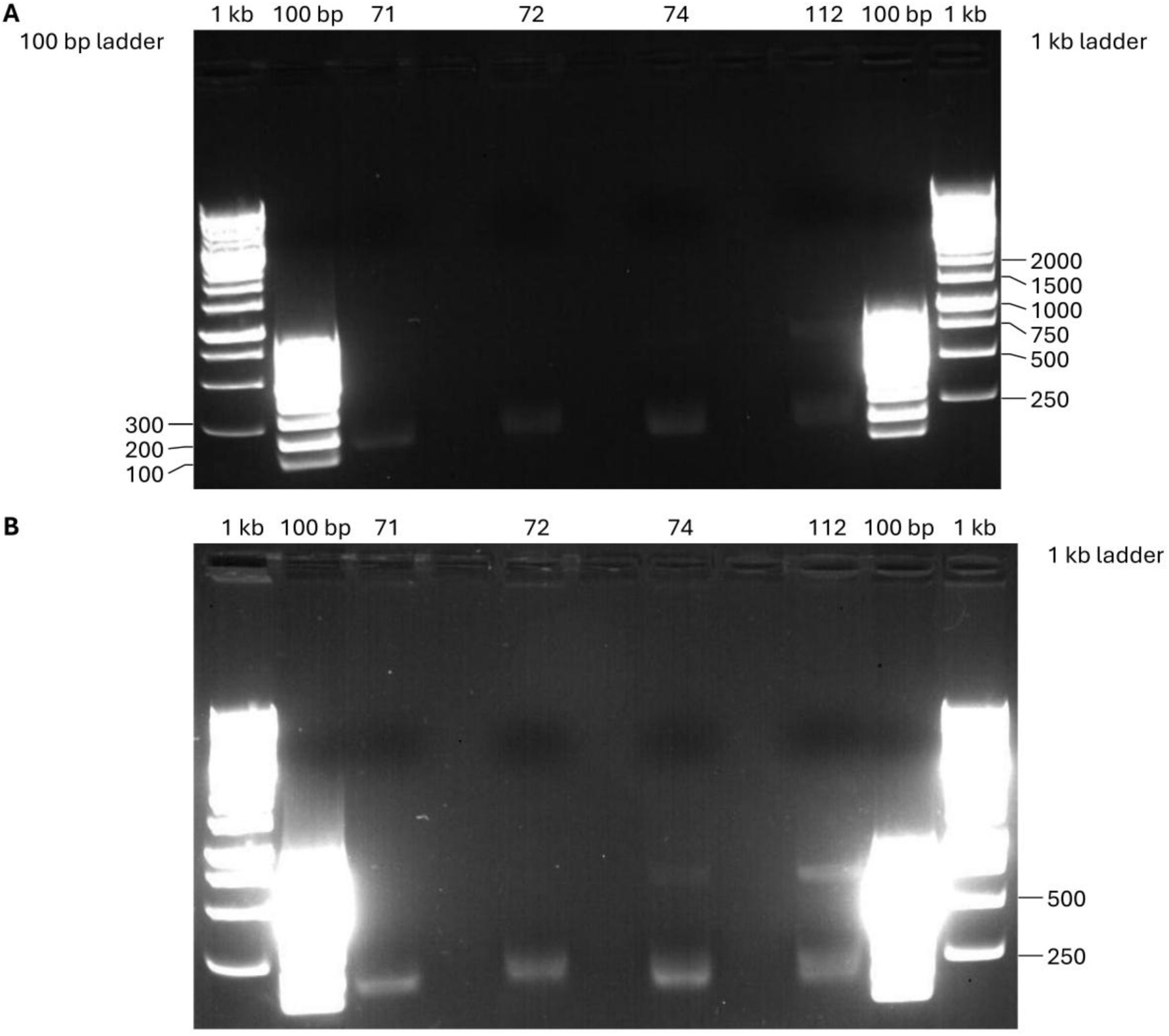
Gel panels reproduced from the 2025 laboratory report. The two panels are identified in that report as the same agarose gel at different exposures. The report attributes the approximately 600-bp band at standard exposure to lane 112 (AG-28112) and describes a band in lane 74 (AG-27374) as visible at higher exposure. Lane identities follow the report; lane labels and molecular-size annotations were redrawn uniformly outside the gel images. Ladder bands are labelled with the manufacturer’s nominal sizes for the two ladders listed in the report (GeneRuler 1 kb, SM0311; GeneRuler 100 bp, SM0241), only where a discrete band could be assigned (left margin: 100 bp ladder, second lane; right margin: 1 kb ladder, last lane); the approximately 600-bp description is the report’s estimate, not a calibrated measurement made here. The identity of the 112-B band is not inferred. These panels show observations reported for the 2025 archival assay, not the unavailable 2004 gel and not independent replicates. No image pixels were enhanced, erased, or rearranged for this figure.

Bidirectional Sanger sequencing of the upper band from sample 112, designated 112A, yielded reads of 543 bases with primer D and 540 bases with primer E. Both reads originated from the same gel-purified PCR product. The trace files are named 112A_D and 112A_E, whereas the November 2025 laboratory report lists the sequences as AG-28112_D and AG-28112_E. Their base calls are identical, so the present reanalysis rests on the same November 2025 sequencing data; the attribution of these files to animal AG-28112 relies on the sample labelling in that report. Primers D and E were those published in the original paper,^1^ so the 2025 assay reproduced the original genotyping reaction on descendant tissue, using newly synthesized aliquots of the D/E primer pair. The identity of the separate 112B band, presumably the lower band of lane 112, remains unresolved; both of its sequencing reactions returned no base calls (Supplementary Fig. S3).

### Sequence overlap and identification of the deletion junction

In the reference orientation shown in Figure 3 (*Prnp* side first, corresponding to the plus strand of the chromosome 2 reference), the reverse-complemented D read and the E read overlapped by 476 bases, with 475 matches, one disagreement and no gaps (99.79% identity). Read D covers positions 1–543 of the combined sequence (the contig) and read E positions 68–607. The combined reads spanned 607 positions (543 + 540 − 476), a sequence-assembly length distinct from the approximately 600-bp estimate obtained by gel electrophoresis. Both values agree with the 600-bp product predicted for primers D and E in the original paper.^1^ At position 520, D calls G (Phred base-call quality Q56; Q30 corresponds to an estimated error probability of 0.1%) and E calls A (Q17), so the base is recorded as N in the 607-position sequence; SnapGene consensus calling assigns G, the higher-quality call, which also matches the C57BL/6J reference (Supplementary Tables S3 and S4). The canonical 34-base loxP sequence is supported by both read directions once both are written in the same orientation. The complete 607-position sequence is shown in Table 1, and its sequence features are summarized in Table 2.

**Figure 3.**
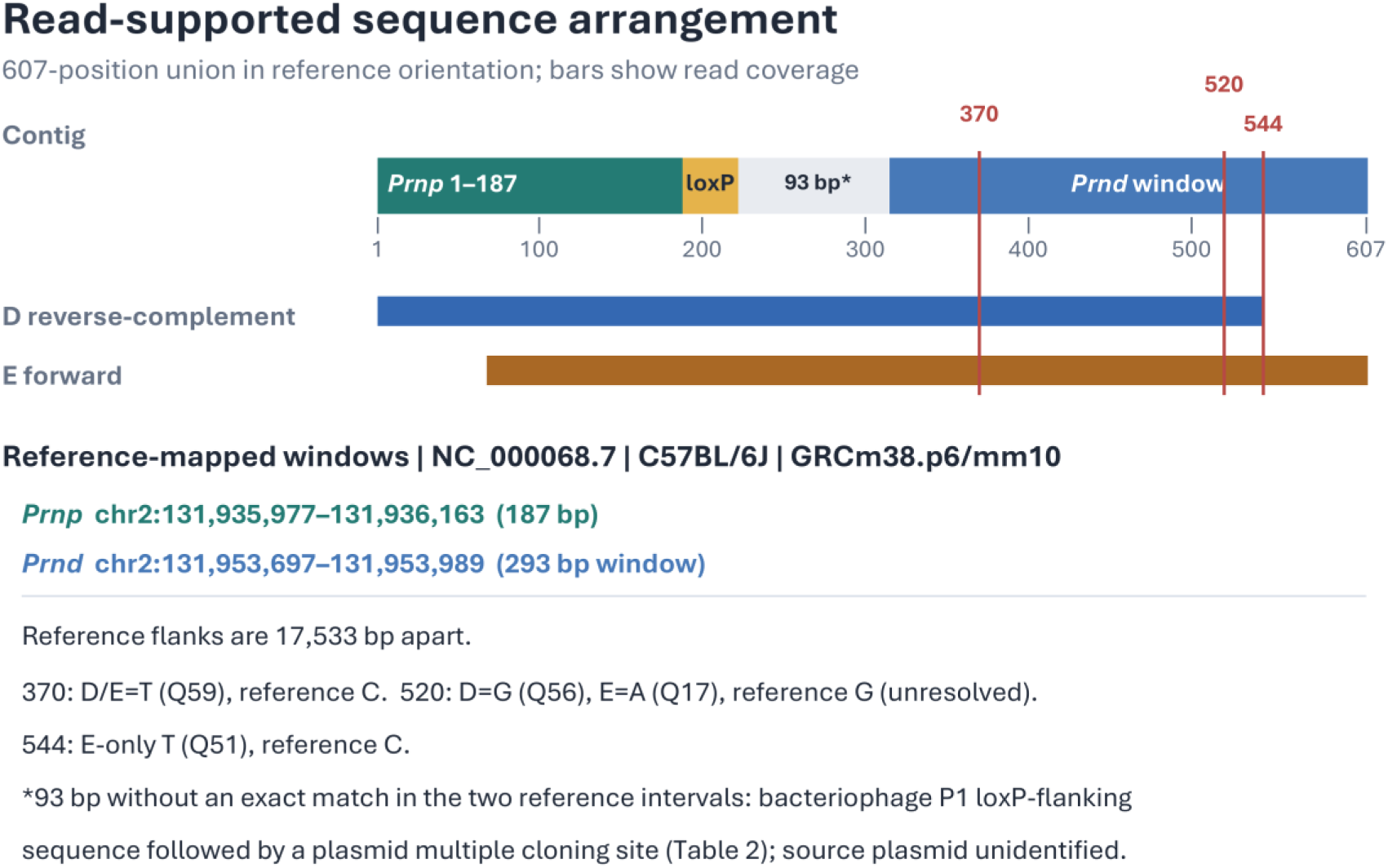
Read-supported sequence arrangement and reference comparison. In reference orientation, the 607-position combined read sequence contains an exact 187-bp *Prnp*-side anchor (NC_000068.7 chr2:131,935,977–131,936,163), reverse-complement-oriented canonical loxP at positions 188–221, 93 bases without an exact match in the two reference intervals searched (bacteriophage P1 loxP-flanking sequence followed by a plasmid multiple cloning site; Table 2), and a gap-free 293-bp *Prnd*-side reference window (chr2:131,953,697–131,953,989) anchored by exact seeds. D (reverse-complemented) and E read coverage is shown separately. The *Prnd* window has a shared T/reference-C difference at position 370, an unresolved G/A D/E disagreement at position 520, and an E-only T/reference-C difference at position 544. The C57BL/6J reference intervals are separated by 17,533 bp; the diagram does not depict the interval without a reference match as a continuous mapped sequence. The same intervals in GRCm39 are chr2:131,777,897–131,778,083 and chr2:131,795,617–131,795,909. Reference placement was limited to those two intervals. The complete sequence is given in Table 1 and the base-by-base alignments in Supplementary Table S5.

**Table 1.**
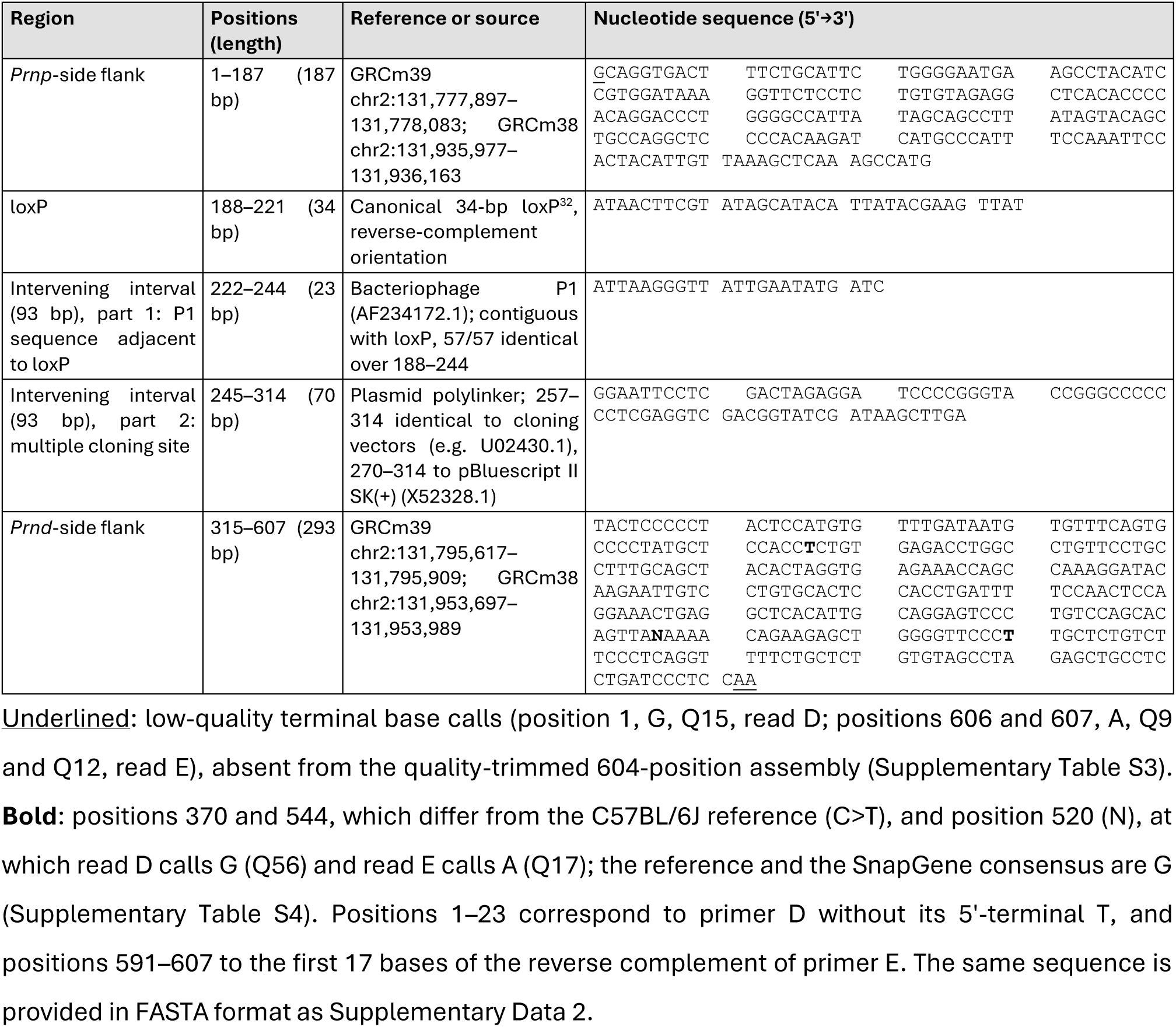
Complete nucleotide sequence of the AG-28112 *Prnp*–*Prnd* junction amplicon. The 607-position sequence assembled from reads 112A_D and 112A_E is shown 5′→3′ in the reference orientation of Figure 3, in consecutive 10-base blocks numbered from the first position of each region (positions are 1-based and inclusive). Sequence and region boundaries were provided by M.B. from the SnapGene assembly and checked against the reference and database entries listed (Table 2).

**Table 2.** Sequence features of the junction amplicon and their reference or database matches. Positions refer to the 607-position sequence in Table 1. Identities are exact matches over the stated positions, without gaps. Wild-type reference separation of the two flanks, 17,533 bp; of the primer D and E annealing sites, 18,020 bp (GRCm38).

| Feature | Positions | Best match (accession: coordinates) | Identity | Notes |
|---|---|---|---|---|
| Primer D site | 1–23 | Primer D, bases 2–24 | 23/23 | Position 1 is a low-quality terminal call (Q15) |
| *Prnp*-side flank | 1–187 | NC_000068.8 chr2:131,777,897–131,778,083 (GRCm38 chr2:131,935,977–131,936,163) | 187/187 | Within *Prnp* (Gene ID 19122); BLASTN of the 604-position assembly: 186/186 over 2–187 |
| loxP | 188–221 | Canonical loxP<sup>32</sup> | 34/34 | Abuts the *Prnp*-side flank directly |
| P1 loxP-flanking sequence | 222–244 | Bacteriophage P1, AF234172.1: 94,784–94,800 and 1–40 (188–244, with loxP) | 57/57 | Also identical to pRAY-1 (U63018.1) over 188–257 (70/70) |
| Multiple cloning site | 245–314 | Cloning vectors, e.g. pMAMneoBlue, U02430.1: 1,804–1,747 (257–314); pBluescript II SK(+), X52328.1: 652–696 (270–314) | 58/58; 45/45 | Sites (first base): EcoRI 246, BamHI 262, SmaI/XmaI 267, KpnI 271, ApaI 277, XhoI 286, SalI 292, ClaI 301, HindIII 307 |
| *Prnd*-side flank | 315–607 | NC_000068.8 chr2:131,795,617–131,795,909 (GRCm38 chr2:131,953,697–131,953,989) | 290/293 (291/293 with G at 520) | Within *Prnd* (Gene ID 26434); C>T at 370 and 544; N at 520 (reference G); BLASTN of the 604-position assembly: 289/291 over 315–605 |
| Primer E site | 591–607 | Reverse complement of primer E, first 17 bases | 17/17 | Positions 606–607 are low-quality terminal calls (Q9, Q12) |

Database matches for the intervening interval are listed in Supplementary Table S6, and base-by-base alignments in Supplementary Table S5.

Alignment to the C57BL/6J chromosome 2 reference intervals NC_000068.7 (GRCm38.p6/mm10) and NC_000068.8 (GRCm39) identified *Prnp*-side and *Prnd*-side sequence matches within the annotated *Prnp* and *Prnd* genes, respectively.^2^ An uninterrupted exact match of 187 bases (the *Prnp* anchor) places the *Prnp* flank at NC_000068.7 chr2:131,935,977–131,936,163. Short exact matches (seeds), extended without gaps and in reference order, place a 293-base aligned segment, the *Prnd*-side window, at chr2:131,953,697–131,953,989; three positions in that window differ from reference or disagree between reads. The corresponding GRCm39 intervals are chr2:131,777,897–131,778,083 and chr2:131,795,617–131,795,909. In reference orientation the combined read sequence comprises the *Prnp* anchor at positions 1–187, loxP at 188–221, 93 intervening positions without an exact match in the two reference intervals searched (222–314), which are characterized further below, and the *Prnd*-side window at 315–607. Contig positions 1–23 match primer D without its 5′-terminal T, and positions 591–607 match the first 17 bases of the reverse complement of primer E, so the reads span the amplicon almost end to end; including the unread primer bases, the predicted product is 614 bp. The wild-type reference intervals are separated by 17,533 bases, and the annealing sites of primers D and E by 18,020 bases. This agrees with the separation of approximately 18 kb between the annealing sites of primers D and E stated in the original paper.^1^ The ordered *Prnp* and *Prnd* anchors, separated by loxP and 93 intervening bases, correspond to the junction predicted for the recombined allele.

Within the *Prnd*-side window, both reads call T at a reference-C position (contig position 370; Q59 in both directions). At the unresolved position 520, the reference is G and D agrees with it, while E calls A at Q17. At position 544, covered only by E, the read calls T and the reference is C (Q51). The source of these differences remains unresolved. The archived strain is unknown, and the C57BL/6J comparison covered only the two reference intervals containing the *Prnp* and *Prnd* matches, not the whole genome. The EMMA strain is a B6.129P2 congenic line,^20^ in which the region flanking the targeted locus is expected to remain 129P2-derived, but no assembled 129P2 reference is available. In the 129S1/SvImJ strain assembly,^29^ both windows are identical to C57BL/6J, and no variant catalogued by Ensembl coincides with positions 370, 520 or 544. A C57BL/6J–129 difference is therefore unlikely to explain these positions, although 129P2-specific variants cannot be excluded. The differences at positions 370 and 544 are C>T transitions, the substitution type expected from cytosine deamination (see below). Figure 3 shows the read coverage, reference alignments and intervening interval; base-by-base alignments are given in Supplementary Table S5.

The original paper reported that the breakpoint had been sequenced and that sequencing confirmed the deletion allele but presented these results as “data not shown”.^1^ The present archival analysis provides the underlying reads and their reference comparison, including the local sequence order, the 93 intervening positions and the observed base differences. It therefore adds an inspectable nucleotide-level description to the molecular characterization presented in the original article. The complete sequence is printed in Table 1 and provided in FASTA format as Supplementary Data 2.

Of the ten available Sanger chromatograms, only 112A_D and 112A_E yielded called sequence. Each of the remaining eight files contained five N calls (N denoting an undetermined base) with zero base-call quality and was classified as a failed sequencing read (Supplementary Fig. S3). The ten files represent technical sequencing records rather than independent biological replicates.

### Independent reassembly from the Sanger chromatogram files

To test whether the 607-position sequence depended on how it had been reconstructed, M.B. reassembled the two reads de novo from the 2025 Sanger chromatogram files (112A_D.ab1 and 112A_E.ab1) with the CAP3 algorithm^30^ in SnapGene sequence-analysis software (Supplementary Table S3). With automatic quality trimming (removal of low-quality read ends), the assembly comprised 604 positions; without trimming, it comprised 605. The two assemblies were generated in opposite orientations; after reverse complementation of one of them, they agreed at all 604 shared positions and differed only by one additional terminal G in the untrimmed assembly. Assembly of the archived FASTA base calls (543 and 540 bases) in SnapGene reproduced the 607-position sequence. The length differences are therefore confined to three terminal bases with low base-call quality: the G at position 1 (read D, Q15), which is retained in the 605- and 607-position sequences, and the A residues at positions 606 and 607 (read E, Q9 and Q12), which are retained only in the 607-position sequence (Table 1; Supplementary Table S4). All three bases match the C57BL/6J reference and lie within the primer-annealing sites at the ends of the amplicon. The junction, the loxP site and the intervening interval are identical in all assemblies. Manual inspection of the chromatograms confirmed the single internal disagreement at position 520 (read D, G at Q56; read E, A at Q17). SnapGene assigns G at this position, consistent with the higher quality of read D, whereas the 607-position sequence records N to retain the disagreement. We retain the 607-position sequence for consistency with Figure 3 and with the coordinates used throughout and mark the three low-confidence terminal bases and position 520 in Table 1.

BLASTN^31^ comparison of the quality-trimmed 604-position assembly with the NCBI mouse genomic and transcript database aligned its positions 1–291 to the *Prnd* region of GRCm39 chromosome 2 (NC_000068.8; 289 of 291 bases identical, no gaps) and its positions 419–604 to the *Prnp* region (186 of 186 bases identical, no gaps), both in reverse orientation relative to the reference. The 127 positions between them (292–418) comprised the canonical 34-base loxP sequence^32^ and the 93 intervening bases. In the coordinates of the 607-position sequence, these intervals correspond to positions 315–605, 2–187 and 188–314, respectively, and the two mismatches in the *Prnd* alignment are the C>T differences at positions 370 and 544. The independent assembly and alignment therefore reproduce the regional assignments obtained by the comparison restricted to the two reference intervals (Table 2; Supplementary Table S5).

### Composition of the 93 intervening bases

The 93 intervening bases (positions 222–314) are covered by both reads with identical base calls, so their nucleotide sequence is established; what remained open was their origin. BLASTN searches of the NCBI core nucleotide database (Methods) returned no match to natural mouse sequence for this interval; all significant matches were cloning vectors and synthetic or transgenic constructs (Table 2; Supplementary Table S6). Positions 188–244, comprising the loxP site and the following 23 bases, are identical over all 57 bases to the loxP site of bacteriophage P1 and its native flanking sequence (GenBank AF234172.1),^33^ and positions 188–257 are identical to the loxP segment of Cre/loxP cloning vectors such as pRAY-1 (U63018.1; 70 of 70 bases). Positions 245–314 form a contiguous multiple cloning site (a synthetic stretch of closely spaced restriction-enzyme sites used to insert DNA into plasmids, also called a polylinker) containing recognition sequences for EcoRI, BamHI, SmaI/XmaI, KpnI, ApaI, XhoI, SalI, ClaI and HindIII. Positions 257–314 are identical to the polylinker of several cloning vectors (58 of 58 bases; for example pMAMneoBlue, U02430.1), and positions 270–314 to the polylinker of pBluescript II SK(+) (45 of 45 bases; X52328.1).^34^ No single database entry was identical over the whole interval. The intervening interval therefore consists of P1-derived loxP-flanking DNA followed by plasmid polylinker sequence, as expected for DNA carried over from a targeting construct at a recombined loxP site, and contains no detectable mouse genomic sequence. Because the sequence of the targeting vector used to construct the *Prnd* allele was not available, the source plasmid could not be identified directly.

## Discussion

The paired reads place a *Prnp*-side flank, loxP and a *Prnd*-side flank in the expected order in sample 112A. The present evidence adds a detailed description of the recovered junction from archival tissue, including the mapped flanks, loxP site and 93 intervening bases. Because the *Prnp*-side sequence abuts loxP directly, the 93 bases lie on the *Prnd* side of the recombined loxP site, where sequence carried over from the *Prnd* targeting construct, such as vector polylinker or residual cassette sequence, would be expected; a database search showed that these bases consist of bacteriophage P1 sequence adjacent to loxP followed by a plasmid multiple cloning site (Table 2; Supplementary Table S6). The targeting-vector sequence itself was not available, so the source plasmid could not be identified directly. The complete deletion structure, zygosity and exact source plasmid of the intervening bases remain unresolved; the reference comparison covered the two C57BL/6J intervals and the corresponding 129S1/SvImJ sequences.

Our reanalysis of the trace files reproduces the 475/476 overlap including its unresolved base disagreement. An independent de novo reassembly of the trace files in SnapGene reproduced the same internal sequence; the 604-, 605- and 607-position versions differ only at terminal bases with low base-call quality (Supplementary Table S3), and the single internal disagreement, at position 520, sets a high-quality G in read D against a low-quality A in read E. Lane and tissue assignments follow the report’s labels and, for AG-27371 to AG-27374, agree with Genoud’s histobook entries (Supplementary Table S1). No sequence corresponding to the faint AG-27374 band was available for comparison, leaving its identity uncertain: both sequencing reactions submitted for sample 74 returned no base calls (Supplementary Fig. S3), possibly because the faint band yielded too little purified product.

The rationale for this study is primarily forensic: to reassess experimental claims in the original publication after concerns about figure manipulation were raised on post-publication review sites. Retention of the complete paraffin archive made this investigation possible more than two decades after the original work. Its scope followed from the material available: fixed tissue could be examined for the inherited DNA rearrangement, whereas protein-based assays and experiments in living animals could no longer be repeated.

Archived brain tissue provides evidence about the inherited junction in descendants. Independent transgenic and deletion-model studies have shown that Doppel can cause cerebellar degeneration when PrP is absent or insufficient, and that PrP can suppress this toxicity in a model- and expression-dependent manner.^3,4,5^ These studies provide a mechanistic basis for the original interpretation. In our view, the central biological interpretation of the original study remains valid. The present results provide molecular evidence concerning the inherited junction in the archived descendants. However, the undisclosed manipulation of the published image was, and remains, inexcusable, irrespective of the biological interpretation.

Independently of its Figure 1B, the original paper contains evidence for the deletion that does not involve the spliced lane: absence of *Prnp* and *Prnd* transcripts on Northern blots of brain, spleen and testis with probes B and C, absence of PrP and *Dpl* on Western blots of brain, testis and spleen, male infertility of *Prn*^o/o^ mice, and absence of cerebellar degeneration in *Prn*^o/o^ mice derived from the ataxia-prone ZH-II allele (Zurich II, a *Prnp* knockout allele whose extended deletion leads to Doppel expression in the brain).^1^ Genoud’s progress report of September 2003 (see Data availability) further documents genotyping of *Prn*^+/+^, *Prn*^+/o^ and *Prn*^o/o^ offspring with the deletion-specific primers and with primer pairs flanking each loxP site, the identification of 17 *Prn*^−/−^ and 16 *Prn*^+/+^ animals among about 120 offspring by two independent PCRs, and confirmation of the absence of *Dpl* mRNA by RT-PCR with a separate primer set; it also records the repeated failure of the Southern blot that was attempted as an additional confirmation. The junction sequence recovered here, the deposit of the line in a public repository and its transfer to other laboratories add to this body of evidence, while the published image remains defective.

The methods available when the work was done also bear on what the original study could show and on what can be shown now. Dideoxy sequencing on capillary instruments was routine in 2002–2003, but the mouse genome existed only as the draft assembly published in December 2002.^23^ The *Prn* locus was known from cosmid sequence;^24^ a finished C57BL/6J assembly with stable coordinates appeared in 2009,^25^ and it is this reference that permits the base-by-base comparison reported here. Massively parallel sequencing was introduced in 2005–2008^26,27^ and could now be applied to the junction amplicon, or to a longer fragment spanning the deletion, to characterize the complete allele; in 2003 the expected confirmation of a genomic rearrangement was the Southern blot, which failed repeatedly in this case. Extraction chemistries for fixed tissue with heat-assisted reversal of cross-links and silica purification,^11,15^ together with high-fidelity polymerase master mixes, made it possible in 2025 to amplify a 600-bp target from blocks that had been prepared for histology more than two decades earlier. Finally, in 2004 the journal accepted ‘data not shown’ for confirmatory sequencing, whereas deposition of primary sequence data is now expected, and the automated image-comparison tools that brought the original Figure 1B splice to attention did not exist.^17,18^

### Fixation history and chemical damage

The archived tissue was fixed in 4% paraformaldehyde for periods ranging from several hours to several days before paraffin embedding. The blocks had been prepared for routine histopathology: no step was taken to limit formaldehyde exposure or to wash out fixative before dehydration, because nobody anticipated that the material would ever be used for sequencing, and the blocks were then stored at room temperature for more than two decades. Differences between paraformaldehyde and neutral-buffered formalin in formulation and tissue handling limit direct comparison with published formalin studies.

Formaldehyde can modify DNA bases and form covalent links between nucleic acids and nearby proteins. Defined-reactant experiments characterize hydroxymethyl (methylol) adducts and methylene-bridge cross-links^6,7^. These modifications can impede access of polymerases to template DNA. Cytosine deamination can produce uracil, while deamination of methylated cytosine can produce thymine; after amplification these lesions can appear as C>T or complementary G>A substitutions. Separately, hydrolysis and depurination can generate abasic sites and strand breaks. Rait et al. observed modified nucleotide and depurination products in a model of formalin fixation and tissue dehydration^6^. In rat liver tissue, Tokuda et al. detected DNA degradation before extraction and found that immediate buffered fixation at low temperature or EDTA reduced it, implicating tissue nucleases under their tested conditions^8^. Reversal of cross-links can improve template accessibility, while strand breaks and base modifications may persist.

Fixation effects vary with formulation, pH, exposure time, tissue, and target length. Karlsen et al. found that longer fixation impaired amplification of longer products in their samples: in some tissues, 8 h inhibited products above 421 bp, while after more than 215 h products above 200 bp required 50–80 PCR cycles^9^. In controlled neutral-buffered-formalin studies, Kim et al. reported increased C:G>T:A changes after more than 48 h, with additional effects of fixation pH and delayed fixation^10^.

### Reference methods for recovery

Recovery methods commonly address several different tasks: removing paraffin, making tissue material accessible, digesting protein-rich material, purifying DNA, and assessing the recovered DNA’s ability to serve as a template. Studies have compared enzymatic extraction and purification with heat-assisted approaches intended to reverse or reduce fixation-associated masking. Shi et al. compared conventional enzymatic extraction with heated extraction across pH conditions, assessing both measured DNA yield and PCR amplification^11^. Bonin et al. evaluated a pre-PCR restoration approach on postmortem paraffin material and showed that some longer targets became amplifiable in their specimens^12^. DNA yield, target amplifiability and sequence accuracy therefore require separate evaluation.

Silica-based extraction followed by PCR and gel purification yielded a product suitable for Sanger sequencing with primers D and E. PCR selectively enriches molecules that remain amplifiable after extraction, even when other molecules are damaged.

### Expected quality and observed junction recovery

Recoverable sequence from paraffin-embedded material is possible but variable. In postmortem archival tissue, Bonin et al. reported 287- and 291-base products after their restoration approach^12^. Alqahtani et al. also recovered PCR products of at least 400 bp from excessively formalin-fixed tissues, demonstrating that useful longer targets can sometimes survive extensive fixation^13^. Karlsen et al. showed the opposite practical constraint: longer products were more sensitive to fixation in their tissues^9^. Against these study-specific comparisons, recovery of the present target, with its approximately 600-bp gel migration estimate, is noteworthy. Recovery of useful sequence from extensively fixed tissue is nevertheless well documented. The expectation that the blocks might still yield amplifiable DNA rested on the experience that prolonged formaldehyde exposure damages DNA, whereas dehydration through graded alcohols and long-term storage in paraffin preserve it; this reasoning was recorded before the experiment, in correspondence of 12 August 2025. A readable PCR product can arise from a small surviving fraction of molecules that span the target. Enrichment of such templates offers a plausible explanation for this recovery, although the fraction of amplifiable DNA in these extracts was not measured.

An approximately 600-bp band was detected in AG-28112, and a weaker band in AG-27374 was visible at higher exposure; no corresponding band was detected in wild-type AG-27371. The upper band from sample 112 (112A) was selected for bidirectional Sanger sequencing. The 543-base D and 540-base E reads came from the same gel-purified PCR product: they are opposite sequencing directions, not independent PCRs, extractions, or biological samples. Our reanalysis found a 476-position overlap with 475 agreements and one unresolved disagreement. The two reads together cover 607 positions, or 604 positions after quality trimming of three terminal bases (Supplementary Table S3); the gel provides a separate, approximate estimate of product size.

Of the ten available trace files, only 112A_D/E have called sequence; the other eight have five N calls and zero base-call quality each. The cause of these failed reads is uncertain and may involve template quality, amplification or sequencing. In D, 449/543 called bases have Q30 or higher; in E, 452/540 do. These quality scores describe the sequencing reads; they do not detect errors already present in the amplified template. The recorded DNA concentration for AG-28112 was 93.1 ng/µL, but the proportion of DNA supporting the target amplification was not measured. The reads support a 34-bp loxP motif and *Prnp*/*Prnd*-side sequence placement in a comparison restricted to the two reference intervals. Reference differences could reflect strain variation not represented in the available references, fixation damage or errors during amplification or sequencing. These data do not distinguish among those explanations. Together, the reads support the expected local junction arrangement in sample 112A.

### Interpretation

Amplifiable junction DNA was recovered from the archived material, consistent with previous studies of sequence recovery from fixed tissue. The fraction of intact templates was not measured, and the fragment-size distribution was assessed only qualitatively, on an agarose gel that showed extensive fragmentation. Williams et al. found nonreproducible sequence changes in formalin-fixed archival samples, with artifact frequency increasing when fewer cells were used in PCR^14^. The D/E reads agree at 475 of 476 overlapping positions, leaving one base unresolved (position 520, at which the higher-quality call, in read D, matches the reference). Both were obtained from the same PCR product, so this agreement does not provide an independent check for errors introduced before sequencing. The recovered sequence nonetheless provides useful evidence about the junction in the archived material.

### Concluding remarks

The image manipulation that prompted this study is deplorable. Splicing a control lane into a gel image misrepresents an experiment whatever the motive, and this was as clear in 2003 as it is today: the standards of scientific record-keeping have not changed in the intervening years, only the tools for detecting breaches of them. Nothing in the present results lessens that judgement, and the correction of the original article records the splice irrespective of what the archival analysis found.

The purpose of this study was correspondingly limited. It was not, and could not be, a reproduction of the 2004 paper: the original gels and homogenates no longer exist, the colony has been discontinued, and observations in living animals cannot be repeated from fixed tissue. What could be tested was tested. Archived brain tissue of an animal recorded as *Prn*^−/−^ contains the *Prnp*–loxP–*Prnd* junction predicted by the published breeding scheme, obtained with the primers, at the product size and with the flanking sequences of the original description, and a second animal of the same recorded genotype gave a fainter band of the same size, which could not be sequenced; the mouse line is held in a public repository; and the contemporaneous presentations, manuscript drafts, correspondence and animal licences describe the same experiments, numbers and dates as the published paper. Every point that could be checked gave a result compatible with the essential truthfulness of the study. The existence of these mice and the predicted local structure of the engineered junction are, in our assessment, confirmed. How much weight to give the remainder of the 2004 paper, in the light of an acknowledged manipulation of one of its figures, is a judgement that we leave to the reader; our obligation was to make the surviving evidence available in full, and that is what this report attempts to do.

## Materials and methods

### DNA extraction from archival tissue

DNA was extracted from archival paraffin-embedded tissue using the QIAamp DNA FFPE Tissue Kit (QIAGEN), following the manufacturer’s instructions. The tissue had been fixed in 4% paraformaldehyde for periods ranging from several hours to several days before embedding; specimen-specific fixation times were not recorded. The sample records designated AG-27371 as wild type, AG-27372 as *Prp*^−/−^, and AG-27374 and AG-28112 as *Prn*^−/−^ (Supplementary Table S1). Sections were cut from the blocks in July 2025; two initial extractions yielded DNA of insufficient quality for PCR, and the extraction was repeated with additional sections. Extract integrity was checked by electrophoresis of 10 µL per sample on a 0.5% agarose gel alongside a 1-kb ladder on 18 August 2025 (Supplementary Fig. S2).

### Principle of the QIAamp FFPE method

Paraffin removal exposes the tissue to the lysis reagents. The method combines proteolytic lysis, heat-assisted reduction of fixation cross-links and silica-membrane purification. Proteinase K digests tissue proteins and releases DNA, while heating helps reverse a fraction of the formaldehyde-induced cross-links. Guanidine-containing binding buffer and ethanol promote DNA adsorption to the silica membrane; subsequent washing removes soluble contaminants, and low-salt elution releases the DNA into aqueous solution.^15,16^ This approach is suited to archived paraffin-embedded tissue because it addresses protein cross-linking and removes substances that can inhibit PCR while recovering fragmented DNA. It does not repair strand breaks or reconstruct DNA molecules lost during fixation and storage.^15^

### PCR amplification and Sanger sequencing

The junction was amplified with UltraII Q5 High-Fidelity 2× Master Mix and primers D (5′-TGCAGGTGACTTTCTGCATTCTGG-3′) and E (5′-TGAGTCTTGGAGGGATCAGGAGG-3′), identical to primers D and E of the original paper. PCR products were separated on a 1.5% agarose gel. The selected band was excised, purified with the NucleoSpin Gel and PCR Clean-up Kit (Macherey-Nagel), and submitted to Microsynth for Sanger sequencing with primers D and E. The two sequencing reactions used the same purified PCR product. In preliminary trials on the same extracts, Phusion and GoTaq polymerases gave no or only faint products, whereas the Q5 master mix amplified products from all samples; an initial Q5 reaction with 100 ng genomic DNA in 50 µL and 35 cycles gave the expected band, and the reaction was therefore repeated as three parallel 50-µL reactions per sample, each containing 25 µL of 2× Q5 master mix, 100 ng genomic DNA, 0.5 µL of the D/E primer mix (recorded as 10 µM) and water, to obtain enough product for gel extraction and sequencing. Cycling conditions were 98 °C for 30 s, followed by 36 cycles (35 in the initial reaction) of 98 °C for 10 s, 60 °C for 30 s and 72 °C for 20 s, and a final extension at 72 °C for 5 min.

The archival assays were performed in 2025, and the resulting chromatograms and base-call files were reanalysed in 2026. Figure 2 reproduces two gel panels identified in the 2025 laboratory report as exposures of the same gel. Lane identities follow the report; lane labels and molecular-size annotations were redrawn uniformly without altering the gel images.

### Sequence analysis and reference alignment

Ten Sanger chromatograms and their corresponding FASTA files were analysed. Base-call quality was extracted from the per-base quality values stored in the trace files (PCON1 field of the ABIF file format). For sample 112A, read E was reverse-complemented for the initial alignment to read D. The resulting assembly was subsequently represented in the reference orientation used in Figure 3, in which read D is reverse-complemented and read E retains its original orientation. Because primer D lies at the *Prnp* end of the amplicon in this orientation and a Sanger read does not contain its own primer, the file designated D, which spans the primer D site, must have been extended from primer E, and vice versa; the D/E designations of the two files therefore appear to be interchanged relative to the priming oligonucleotide, without affecting the assembly. Ungapped overlaps of at least 30 bases were scored +2 per match and −3 per mismatch and ranked by score, length and sequence identity. The selected overlap was checked by Smith–Waterman local alignment with match, mismatch and gap scores of +2, −3 and −5, respectively. Disagreement between the reads was represented by N in the combined sequence. The same base is assigned G by SnapGene consensus calling (see below).

Reference alignment was restricted to the downloaded *Prnp* and *Prnd* intervals of C57BL/6J chromosome 2, using accessions NC_000068.7 (GRCm38.p6/mm10) and NC_000068.8 (GRCm39). Exact 17-base matches were used as seeds and extended maximally within these intervals. Reference differences and unassigned sequence were recorded separately. Primer binding sites were located by exact matching of primer D and of the reverse complement of primer E to NC_000068.7, and gene boundaries were taken from NCBI Gene (*Prnp*, Gene ID 19122; *Prnd*, Gene ID 26434). Both windows were also compared with the 129S1/SvImJ strain assembly (129S1_SvImJ_v3, GCA_921998555.2; chr2:128,810,664–128,810,850 and chr2:128,828,382–128,828,674), and variants catalogued in the *Prnd* window (GRCm39 chr2:131,795,617–131,795,909) were retrieved; both queries used the Ensembl REST interface (release 116).^29^ Analyses were implemented with Python standard-library scripts supplied with the accompanying data package.

### Independent reassembly and BLAST alignment

M.B. reassembled reads 112A_D and 112A_E independently, starting from the 2025 Sanger chromatogram files (112A_D.ab1 and 112A_E.ab1) rather than from the previously reconstructed sequence. De novo assembly was performed with the CAP3 algorithm^30^ as implemented in SnapGene version 9.0.0 (Dotmatics), once with automatic quality trimming and once without preliminary trimming; the two assemblies, which were generated in opposite orientations, were compared after reverse complementation of one of them. CAP3 was run with its default settings, including a base-quality cutoff of 12 for clipping, a base-quality cutoff of 20 for differences, a minimum overlap length of 40 bp, a minimum overlap identity of 90%, and end clipping enabled (flag = 1). Preliminary quality trimming in SnapGene used the medium-stringency setting, for which no separate numerical Phred threshold was specified. The archived FASTA base calls were assembled in the same way. Chromatograms and Phred quality values at the terminal positions and at the discordant position were inspected manually in SnapGene. The quality-trimmed 604-position assembly was then compared by BLASTN^31^ with the NCBI mouse genomic and transcript database, which aligned it to GRCm39 chromosome 2 (NC_000068.8). Position p of the 604-position assembly corresponds to position 606 − p of the 607-position sequence.

### Database search of the intervening interval

The 93 intervening positions were compared with the NCBI core nucleotide database (core_nt) by BLASTN^31^ through the NCBI BLAST web service on 7 November 2026, with all three searches repeated manually by M.B. (see Use of AI tools) on 8 November 2026: positions 222–314 with blastn (word size 11), positions 222–256 with blastn using short-query settings (word size 7) and the complete 607-position sequence, with G at position 520, with megablast. Hits were annotated with their GenBank titles. Exact identity with the bacteriophage P1 genome (AF234172.1)^33^ and with pBluescript II SK(+) (X52328.1)^34^ was initially assessed by longest-common-substring comparison on both strands, with the P1 genome treated as circular. These matches were subsequently checked by M.B. with pairwise BLASTN. Additional pairwise comparisons were performed against cloning vectors pRAY-1 (U63018.1) and pMAMneoBlue (U02430.1). Restriction-enzyme recognition sites were located by exact string matching. The base-by-base alignments in Supplementary Table S5 were generated from the same reference intervals (NC_000068.8) and database entries.

## Supporting information

Supplementary Data 2. AG-28112 junction sequence (607 positions; FASTA format)

Supplementary Data 1. Source data, reference sequences, analysis scripts and outputs

## Data availability

Raw Sanger chromatograms and base calls, reference intervals, analysis scripts, and analysis outputs are assembled as Supplementary Data 1 (source-data.zip) for deposition with this manuscript. The complete 607-position junction sequence is printed in Table 1 and provided in FASTA format, with N at the discordant position 520, as Supplementary Data 2 (SupplementaryData2_AG28112_junction_607.fasta). The sequence has been submitted to GenBank (submission ID SUB16620894; accession number pending). Figure source data and vector/raster figures are packaged separately. The laboratory report and archival specimen metadata are not included in the public data package; access and reuse are subject to confirmation of custody and permissions. The contemporaneous records cited in the Results (progress-report presentations, manuscript drafts, correspondence and register extracts) are held by A.A. and can be made available to editors and investigators on request. Entries from N. Genoud’s histobook export (Genoud_HistobookMain.xls) are reproduced in Supplementary Tables S1 and S2. Page 7 of the investigation report of 26 November 2025, a printout of histobook entries, is reproduced as Supplementary Fig. S1. Supplementary Documents 1 and 2 contain the complete 2001 licence application and the 2003 registration confirmation of the Kantonales Veterinäramt Zürich, with personal e-mail addresses masked.

## Contributions and acknowledgments

Michele Biase performed the PCR optimisation and amplification, gel extraction, coordination of Sanger sequencing, manual sequence alignment and reference comparison, independent de novo reassembly of the Sanger chromatogram files, chromatogram inspection and BLAST analysis, annotation of the complete junction sequence (Table 1), and prepared the November 2025 laboratory report. Adriano Aguzzi conceived the retrospective analysis, compiled the historical records and prepared the manuscript. We thank Elena De Cecco for extracting and preparing DNA from archived paraffin-derived material; Jingjing Guo for DNA-quality assessment by agarose gel electrophoresis and image acquisition. Microsynth provided Sanger sequencing services.

## Use of AI tools

The BLASTN searches reported in the Methods were initially run through the NCBI BLAST web service by Claude on 7 November 2026. M.B. repeated the searches and inspected the alignments manually on 8 November 2026, with AI-assisted guidance. AI assistants were also used to help organise and draft text, including the summary of those searches and parts of this manuscript.

## Ethics and archived material

All animal experiments underlying the original study were approved by the Veterinary Office of the Canton of Zurich (Kantonales Veterinäramt Zürich), with the agreement of the cantonal animal-experimentation committee, and were performed in accordance with the Swiss Animal Protection Act of 1978 and its 1981 ordinance.^28^ The principal licence was 40/2002 (Bewilligungsnummer 2002040, the cantonal licence number; application signed 19 May 2001 and registered on 6 June 2001 under internal number 2283; approved April 2002; three years requested in the application; case officer Dr H. Sigg; subcommittee Dr F. Rey and Prof. J.-C. Perriard), which covered prion inoculation, transplantation, immunization, tail biopsy for genotyping, ear marking and, under its section IV, the generation and analysis of *Dpl*-deficient mice (Supplementary Fig. S4; Supplementary Documents 1 and 2). Further licences in force during the work were 109/99 (‘Transgene Mäuse zum Studium der Neurodegeneration und Neurokanzerogenese’, until 30 June 2002), 98/2001 (killing of untreated mice as organ donors, successor of 129/98, severity grade 0, notifiable) and 190/2002 (restoration of fertility in *Dpl*-deficient mice); the application for 40/2002 also lists the laboratory’s earlier licences 357/91, 144/92, 148/95, 207/96 and 203/98. The decision letter for 40/2002 has not been retrieved; its issue is documented by the laboratory’s notice of 5 April 2002 instructing that all cage cards carry the number 40/2002, by the veterinary office’s letters of 12 June and 20 August 2002 citing the Bewilligungsverfügung (licence decision) for 40/2002, and by N. Genoud’s addendum requests filed under 40/2002 in December 2002, January 2004 and October 2004. The present analysis used only archived paraffin blocks from animals killed more than twenty years ago; no living animals were used.

---

**Supplementary Figure S1.**
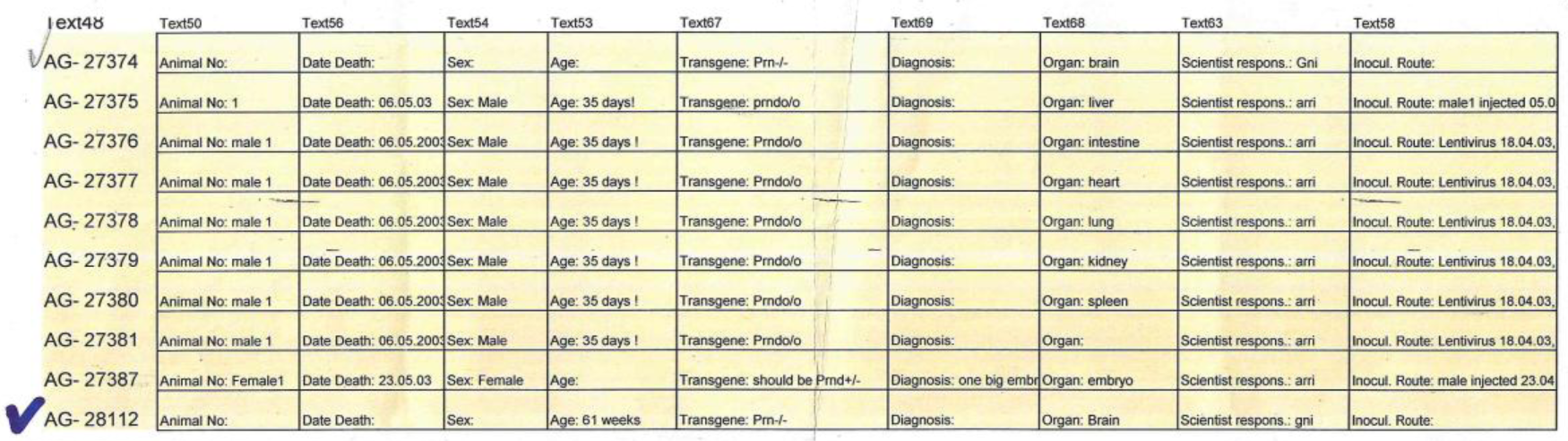
Histobook printout recording AG-28112. Printout of histobook entries (Microsoft Access report of the Institute of Neuropathology histobook; the fields are the same as in Genoud_HistobookMain.xls), scanned as page 7 of the investigation report of 26 November 2025 (M. Biase). The page, printed in landscape, was rotated by 90° and cropped to the table; no other changes were made. The rows list histobook numbers AG-27374 to AG-27381, AG-27387 and AG-28112; AG-27374 and AG-28112 carry hand-drawn ticks. AG-28112 is recorded with ‘Age: 61 weeks’, ‘Transgene: *Prn*^-/-^’, ‘Organ: Brain’ and ‘Scientist respons.: gni’, without a date of death; AG-27374 is recorded as a *Prn*^-/-^ brain (Gni) without age or date of death. AG-27375 to AG-27381 are entries of another scientist for *Prnd*^o/o^ mice aged 35 days that died on 6 May 2003, most of them recorded as injected with lentivirus on 18 April 2003; AG-27387 has a date of death of 23 May 2003. Column headers (Text48–Text69) are internal field names of the database report.

**Supplementary Figure S2.**
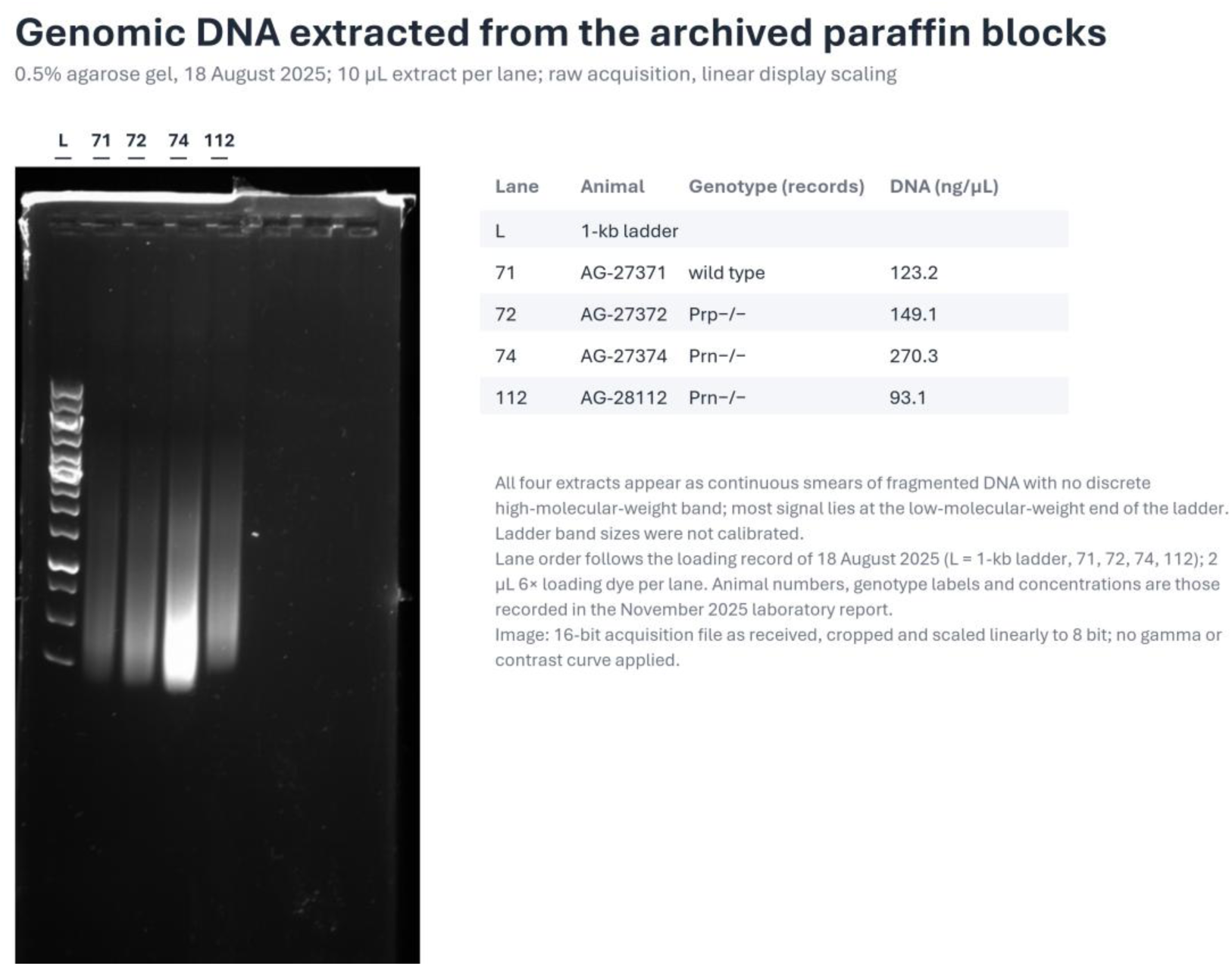
Integrity of genomic DNA extracted from the archived paraffin blocks. Extracts of samples 71 (AG-27371), 72 (AG-27372), 74 (AG-27374) and 112 (AG-28112) were run on a 0.5% agarose gel next to a 1-kb ladder on 18 August 2025 (10 µL extract with 2 µL 6× loading dye per lane). All four lanes show continuous smears of fragmented DNA with no discrete high-molecular-weight band, and most of the signal lies at the low-molecular-weight end of the ladder. Ladder band sizes were not calibrated. The image is the 16-bit acquisition file as received, cropped, labelled and scaled linearly to 8 bit; no gamma or contrast curve was applied. Animal numbers, genotype labels and DNA concentrations are those recorded in the November 2025 laboratory report.

**Supplementary Figure S3.**
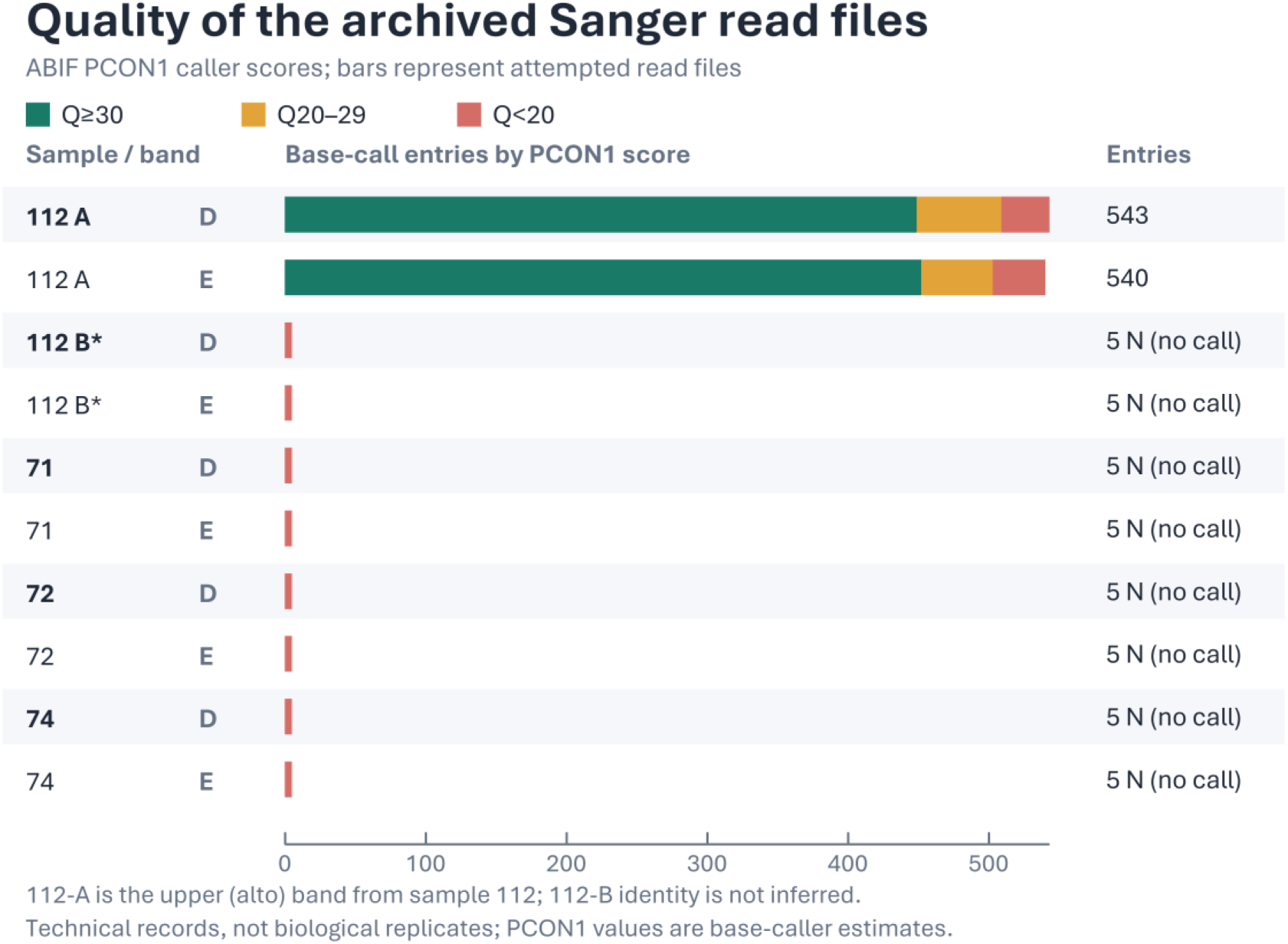
Base-call quality in archived Sanger read files. ABIF PCON1 scores (per-base quality values stored in the trace files) are grouped as Q≥30, Q20–29, and Q<20 for each D/E file. 112-A denotes the upper (“alto”, Italian for upper) band from sample 112; 112-B identity is not inferred. The other eight files contain five N entries each and contain no base calls. These ten files are technical records, not biological replicates; PCON1 values are base-caller estimates.

**Supplementary Figure S4.**
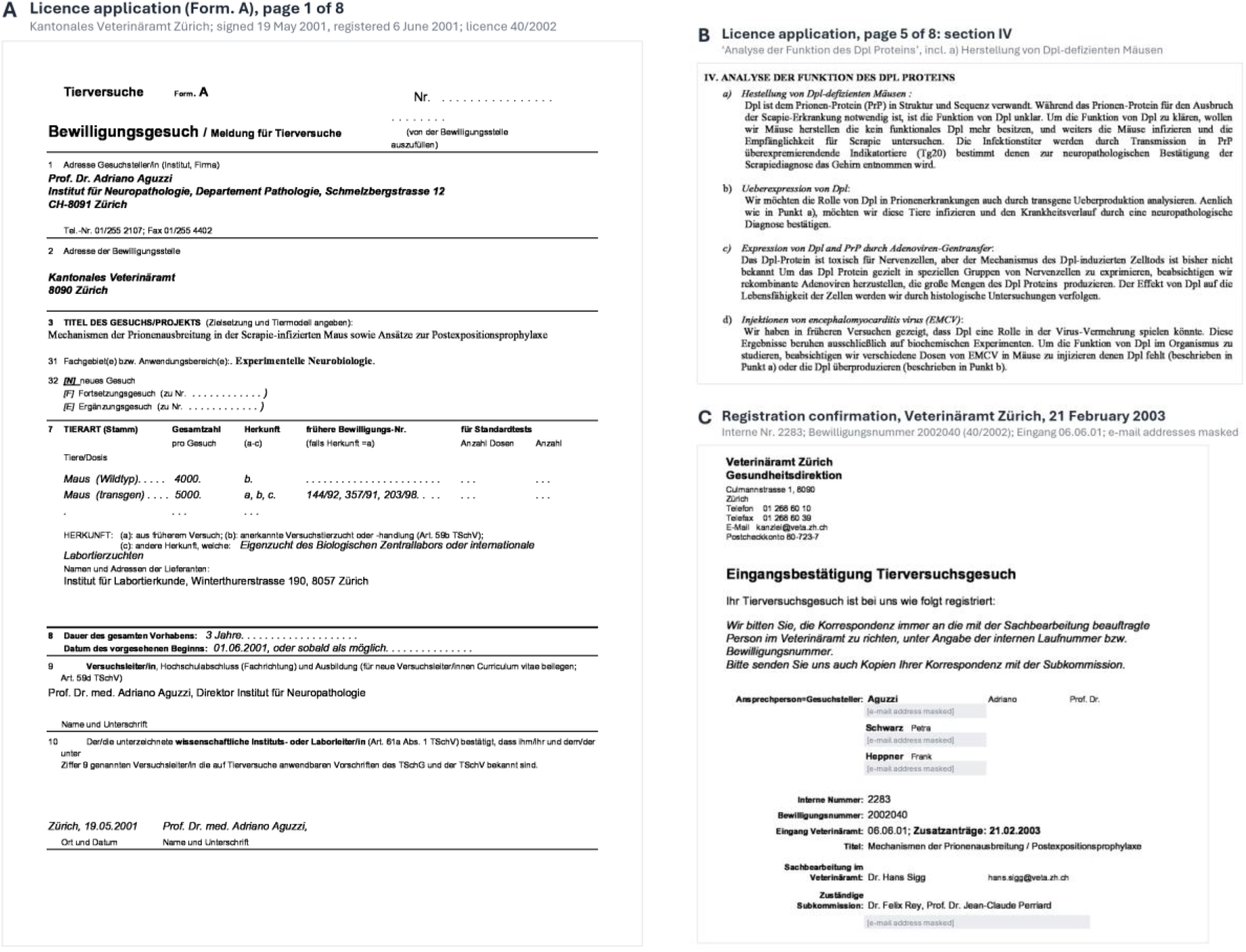
Animal-experimentation licence documents. (A, B) Pages 1 and 5 of the eight-page licence application (Form. A) submitted by A.A. to the Kantonales Veterinäramt Zürich, signed 19 May 2001 and registered by the office on 6 June 2001. Page 1 gives the applicant, the title (‘Mechanismen der Prionenausbreitung in der Scrapie-infizierten Maus sowie Ansätze zur Postexpositionsprophylaxe’), the animal numbers, the earlier licences 144/92, 357/91 and 203/98 and the intended start on 1 June 2001; page 5 contains section IV, ‘Analyse der Funktion des *Dpl* Proteins’, including the generation of *Dpl*-deficient mice. (C) Registration confirmation (‘Eingangsbestätigung Tierversuchsgesuch’) issued by the Veterinäramt Zürich on 21 February 2003 for this application, giving the internal number 2283, the licence number 2002040 (40/2002), the date of receipt (6 June 2001), the receipt of addendum requests on 21 February 2003, the case officer and the responsible subcommittee of the cantonal animal-experimentation committee. Personal e-mail addresses are masked. The complete application and the confirmation are provided as Supplementary Documents 1 and 2.

**Supplementary Table S1.** Histobook entries for the archived specimens analysed. Genotype, organ and scientist are given exactly as entered in the export of N. Genoud’s entries in the histobook, the Institute of Neuropathology’s shared record of the provenance and experimental details of each paraffin block (Microsoft Access report rptHistobookMain, exported as Genoud_HistobookMain.xls; two copies with identical content). The four entries were created together on 28 May 2003 and last modified at 15:44:40 on the same day; their fields for animal number, date of death, sex, age, cage, diagnosis, inoculation and notes are empty.

| Histobook no. | Recorded genotype (as entered) | Organ | Scientist | Entry created | Analysed here (lane) | DNA (ng/µL) |
|---|---|---|---|---|---|---|
| AG-27371 | wild-type | Brain | Gni | 28.05.2003 15:44:38 | Yes (71); wild-type control; no band | 123.2 |
| AG-27372 | PrP<sup>-/-</sup> | Brain | Gni | 28.05.2003 15:44:38 | Yes (72); Prp<sup>−/−</sup> control; no band | 149.1 |
| AG-27373 | Lox1<sup>+/-</sup> | Brain | Gni | 28.05.2003 15:44:38 | No | — |
| AG-27374 | *Prn*<sup>-/-</sup> | brain | Gni | 28.05.2003 15:44:38 | Yes (74); faint ~600-bp band, no sequence obtained | 270.3 |
| AG-28112 | Not in the export | — | — | — | Yes (112); ~600-bp band sequenced (112A) | 93.1 |

Lox1 presumably denotes the *Prnp*<sup>*lox1*</sup> allele (Figure 1A). AG-28112 is absent from the export; the nearest entries are AG-27374 (created 28 May 2003) and AG-28341 (brain from an unrelated PrP-Fc experiment, Plp Cre × Tg550Stop, created 12 November 2003). The histobook printout reproduced in Supplementary Fig. S1 records AG-28112 as a *Prn*<sup>−/−</sup> brain from a 61-week-old animal, with N. Genoud (gni) as responsible scientist and no date of death. Lane numbers refer to Figure 2 and Supplementary Fig. S2; DNA concentrations are those recorded in the November 2025 laboratory report.

**Supplementary Table S2.**
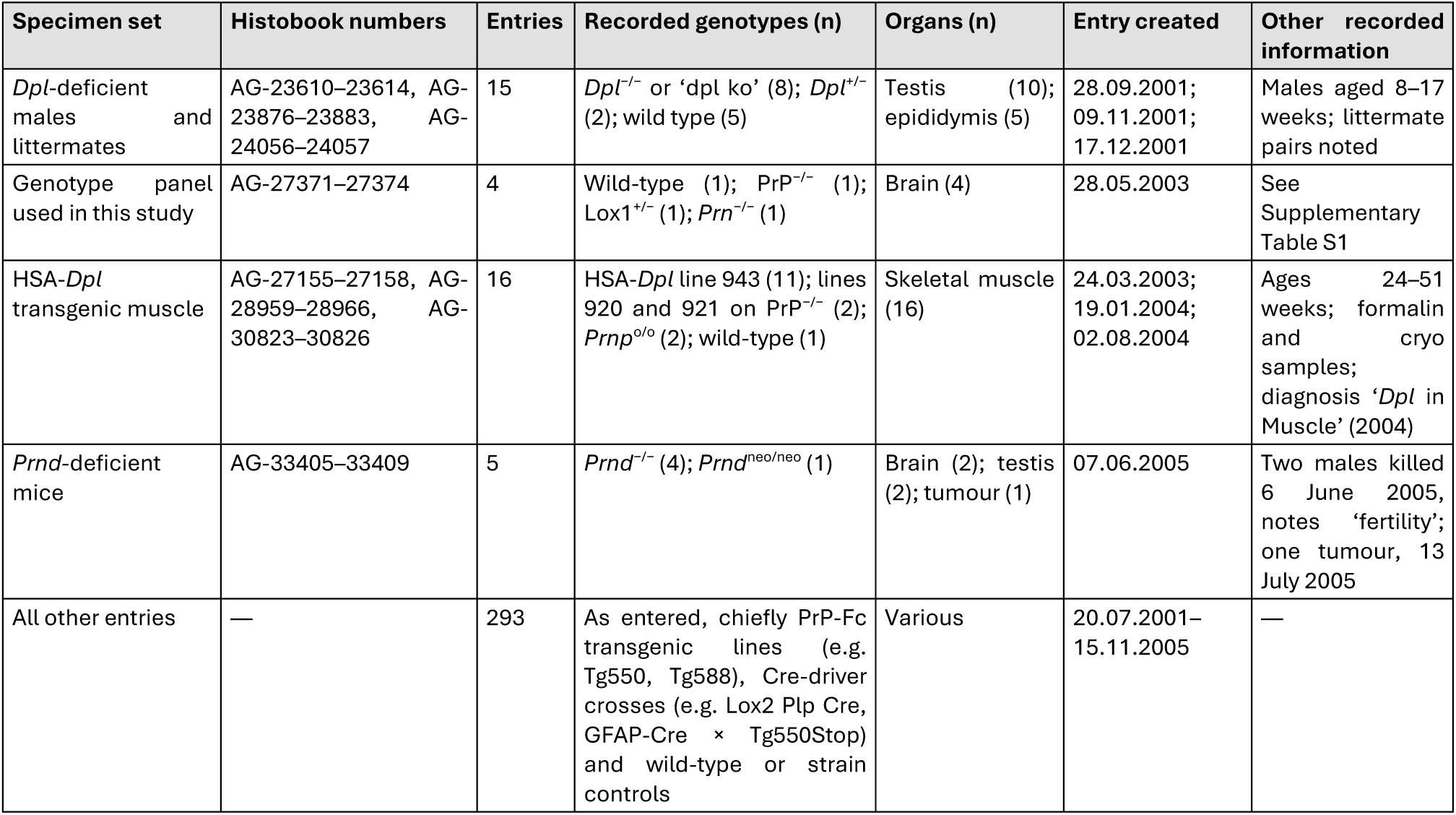
Doppel- and *Prn*-related entries in N. Genoud’s histobook export. The export lists 333 entries (AG-22951 to AG-35348) created between 20 July 2001 and 15 November 2005, all with N. Genoud (initials code GNI) recorded as responsible scientist, alone or with colleagues. Entries were grouped by recorded genotype and creation date; genotype spellings are normalized (as-entered variants: ‘dpl ko’, ‘*Dpl* knockout out’, ‘*Prnp* 00’). AG-27374 is the only entry recorded as *Prn*^−/−^. HSA-*Dpl*, Doppel transgene expressed in skeletal muscle; PrP-Fc, soluble dimeric PrP fused to an immunoglobulin Fc domain; Cre-driver crosses, mice expressing Cre recombinase in defined cell types crossed with Cre-activatable transgenic lines.

**Supplementary Table S3.**
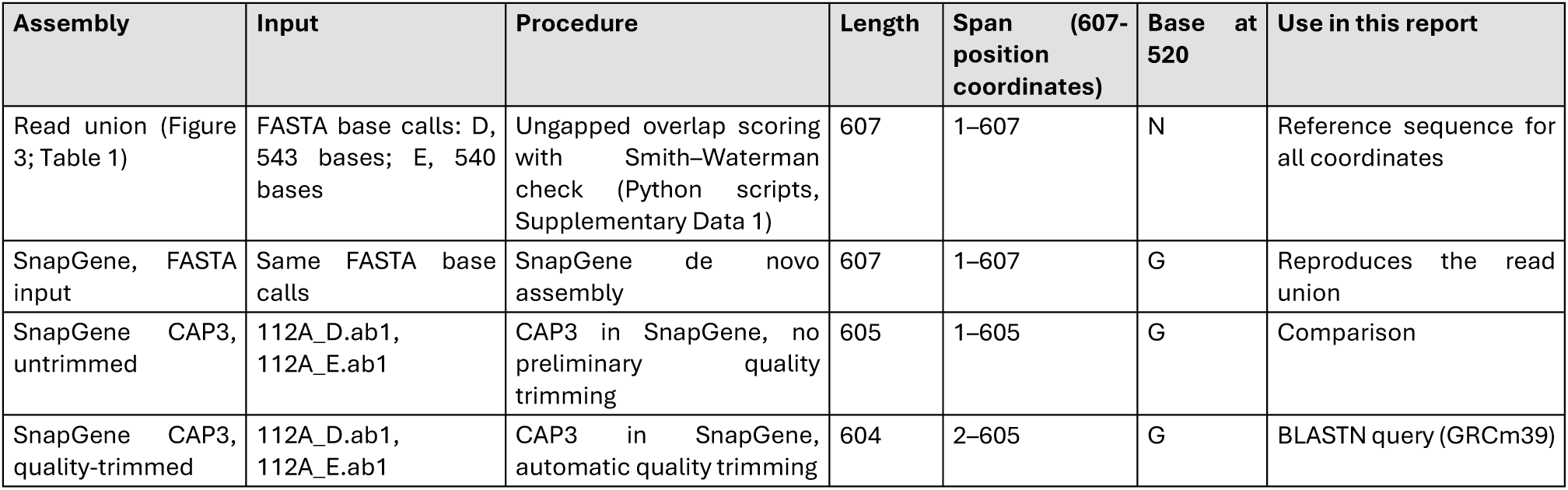
Assemblies of reads 112A_D and 112A_E. Overlap of the FASTA reads: 476 positions, 475 identical, one disagreement (position 520), no gaps; 543 + 540 − 476 = 607. The two CAP3 assemblies were generated in opposite orientations and agree at all 604 shared positions after reverse complementation of one of them; the 605-position assembly carries one additional terminal G (position 1). The 604-position assembly is in reverse orientation relative to Figure 3; its position p corresponds to position 606 − p. SnapGene assemblies and chromatogram inspection by M.B.

**Supplementary Table S4.** Positions with reduced base-call confidence or differences from the reference. Base calls are given in the reference orientation of Figure 3 (read D reverse-complemented). Q, Phred base-call quality (Q20 and Q30 correspond to estimated error probabilities of 1% and 0.1%). Read D covers positions 1–543 and read E positions 68–607. Quality values at positions 1, 520, 606 and 607 are from M.B.’s chromatogram inspection in SnapGene; those at 370 and 544 are the ABIF PCON1 values reported in the Results. No variant catalogued by Ensembl coincides with positions 370, 520 or 544.

| **Position** | **Region** | **GRCm39 chr2 (GRCm38)** | **Reference** | **Read D call (Q)** | **Read E call (Q)** | **607-position sequence** | **SnapGene consensus** | **Retained in 605 / 604 assemblies** |
|---|---|---|---|---|---|---|---|---|
| 1 | Primer D site, *Prnp* flank | 131,777,897 (131,935,977) | G | G (15) | Not covered | G | G (605 only) | Yes / No |
| 370 | *Prnd* flank | 131,795,672 (131,953,752) | C | T (59) | T (59) | T | T | Yes / Yes |
| 520 | *Prnd* flank | 131,795,822 (131,953,902) | G | G (56) | A (17) | N | G | Yes / Yes |
| 544 | *Prnd* flank | 131,795,846 (131,953,926) | C | Not covered | T (51) | T | T | Yes / Yes |
| 606 | Primer E site, *Prnd* flank | 131,795,908 (131,953,988) | A | Not covered | A (9) | A | Excluded | No / No |
| 607 | Primer E site, *Prnd* flank | 131,795,909 (131,953,989) | A | Not covered | A (12) | A | Excluded | No / No |

**Supplementary Table S5.**
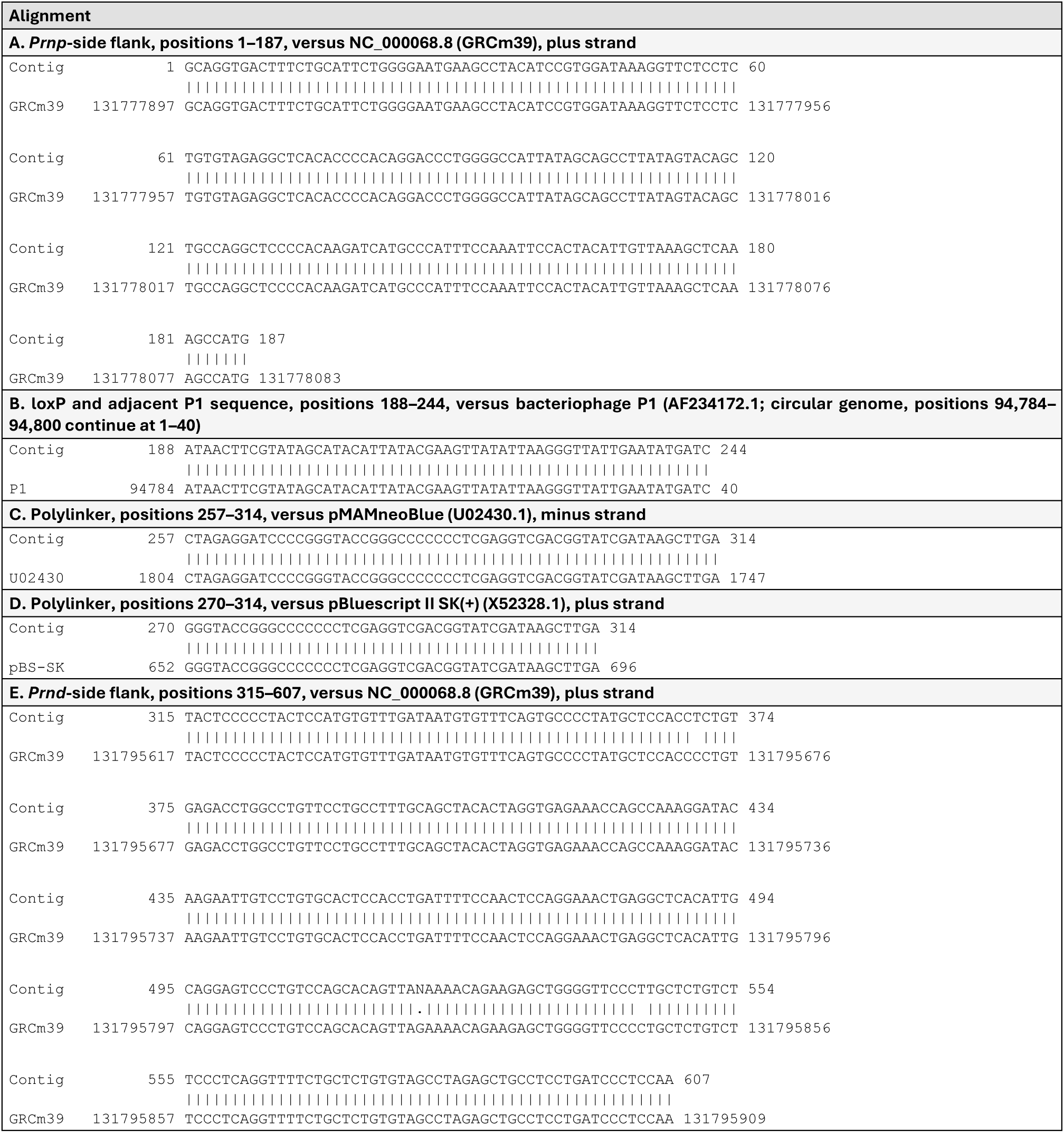
Base-by-base alignments of the junction sequence. Contig positions refer to the 607-position sequence (Table 1). | identical; blank, mismatch; ‘.’, N at position 520 (reference G). The mismatches in panel E are at positions 370 and 544 (C>T). Panel B shows that loxP and the following 23 bases are continuous P1 sequence; panels C and D show the plasmid polylinker. Panels A and E are identical to the corresponding GRCm38 intervals (NC_000068.7 chr2:131,935,977–131,936,163 and 131,953,697–131,953,989).

**Supplementary Table S6.** Database matches of the intervening interval. BLASTN searches of the NCBI core nucleotide database (core_nt) on 7 November 2026 (Methods). Representative top-scoring hits are shown; all significant hits for positions 222–314 were cloning vectors or synthetic or transgenic constructs, and none was natural mouse sequence. For the complete 607-position query (G at 520), the top mouse genomic hits cover only the *Prnp*-side (1–187) and *Prnd*-side (315–607) flanks; the two mismatches in the *Prnd* flank are at positions 370 and 544. Matched positions are given in 607-position coordinates. Subject coordinates in descending order indicate a minus-strand match.

| Query (positions) | Program | Subject | Accession | Matched positions | Identical/aligned | Subject coordinates | E value |
|---|---|---|---|---|---|---|---|
| 222–314 | blastn | Cloning vector pTF1 | S52444.1 | 257–314 | 58/58 | 323–380 | 8.15e-19 |
| 222–314 | blastn | Cloning vector pMAMneoBlue | U02430.1 | 257–314 | 58/58 | 1,804–1,747 | 8.15e-19 |
| 222–314 | blastn | Cloning vector pT3/T7-LUC | U02437.1 | 262–314 | 53/53 | 1,923–1,975 | 4.22e-16 |
| 222–314 | blastn | pBluescript II KS(−) derivative | MT856193.2 | 270–314 | 45/45 | 1,212–1,256 | 9.30e-12 |
| 222–256 | blastn-short | Cloning vector pRAY-2 | U63120.1 | 222–256 | 35/35 | 2,300–2,266 | 1.34e-07 |
| 222–256 | blastn-short | Cloning vector pRAY-1 | U63018.1 | 222–256 | 35/35 | 2,300–2,266 | 1.34e-07 |
| 222–256 | blastn-short | Cloning vector pRAY-Cre | AJ627603.1 | 222–256 | 35/35 | 2,300–2,266 | 1.34e-07 |
| 1–607 | megablast | Mouse BAC RP23-401J24, chromosome 2 | AL833794.9 | 1–187 | 187/187 | 134,070–134,256 | 3.18e-90 |
| 1–607 | megablast | Mouse BAC RP23-401J24, chromosome 2 | AL833794.9 | 315–607 | 291/293 | 151,790–152,082 | 8.18e-146 |
| 1–607 | megablast | Cloning vector pRAY-1 | U63018.1 | 188–257 | 70/70 | 4,728–4,659 | 3.49e-25 |

