## Supplementary figures and images for "Sequencing of the *Prnp–Prnd* deletion junction in *Prn*^-/-^ mouse tissues after formaldehyde fixation and >22 years at room temperature"

### image(20261106-184624).png

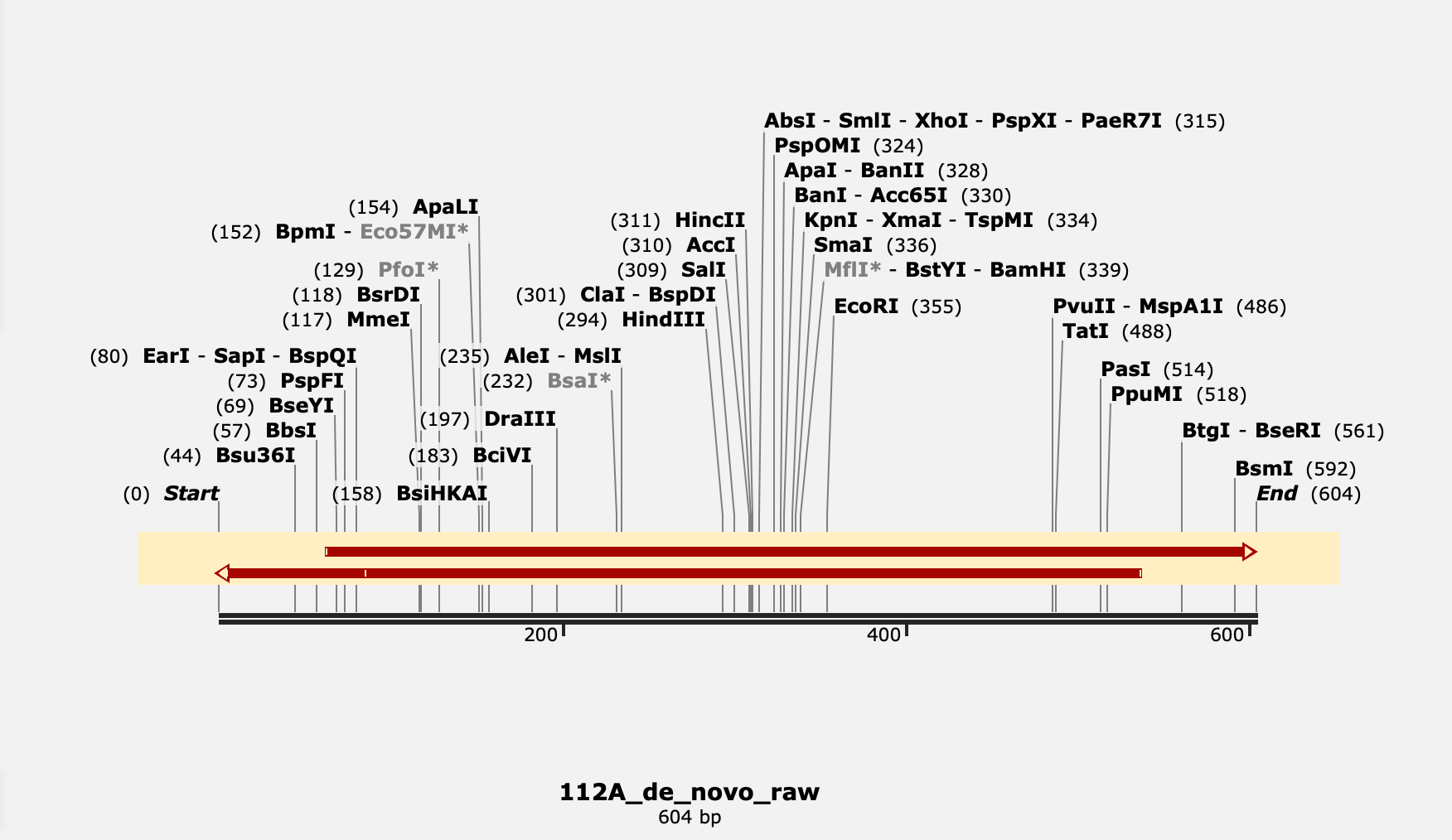

### image(20261106-185051).png

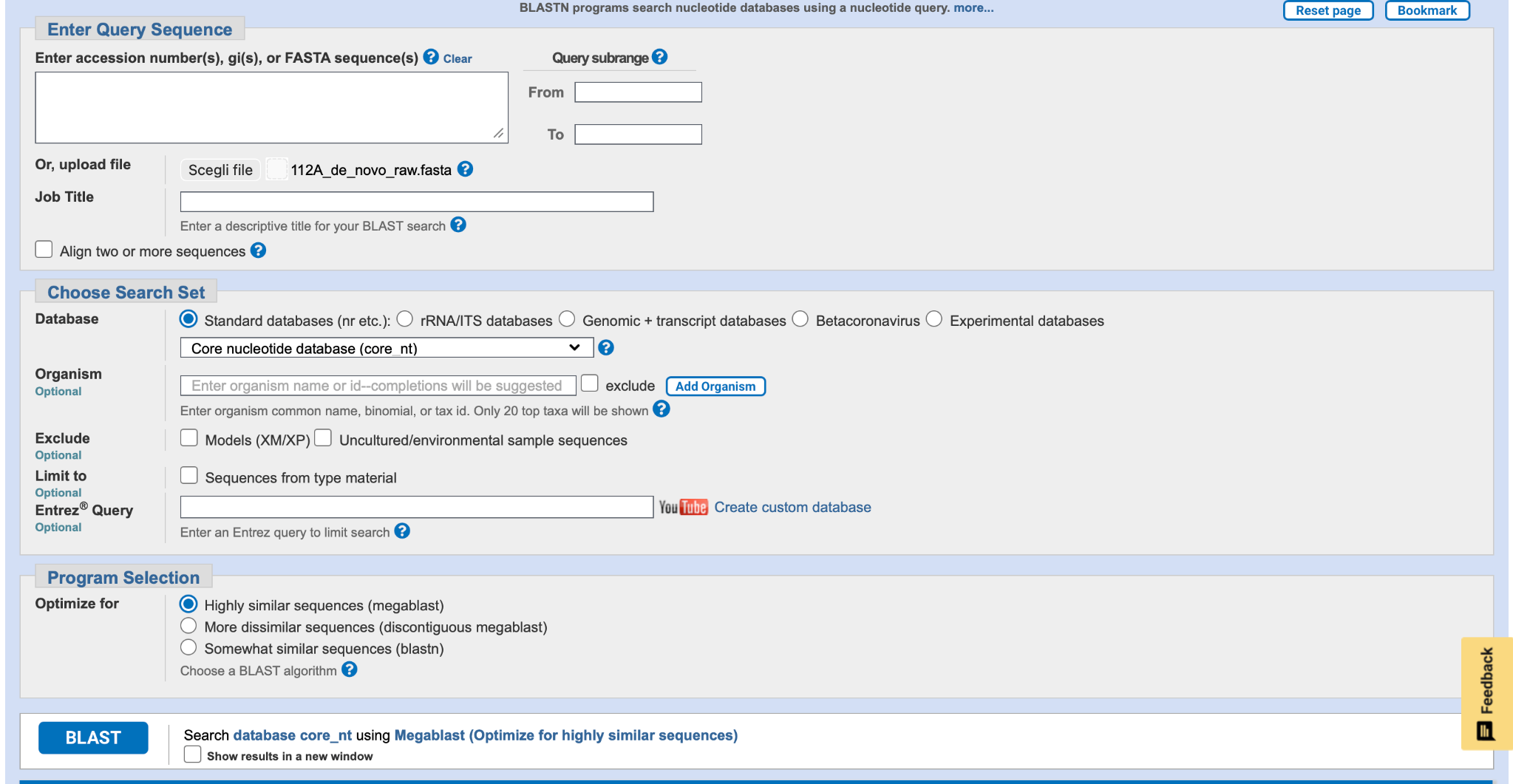

### image(20261106-185217).png

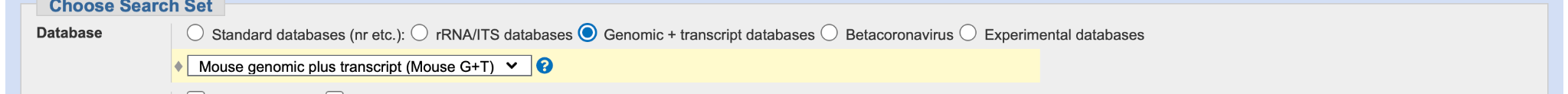

### image(20261106-185417).png

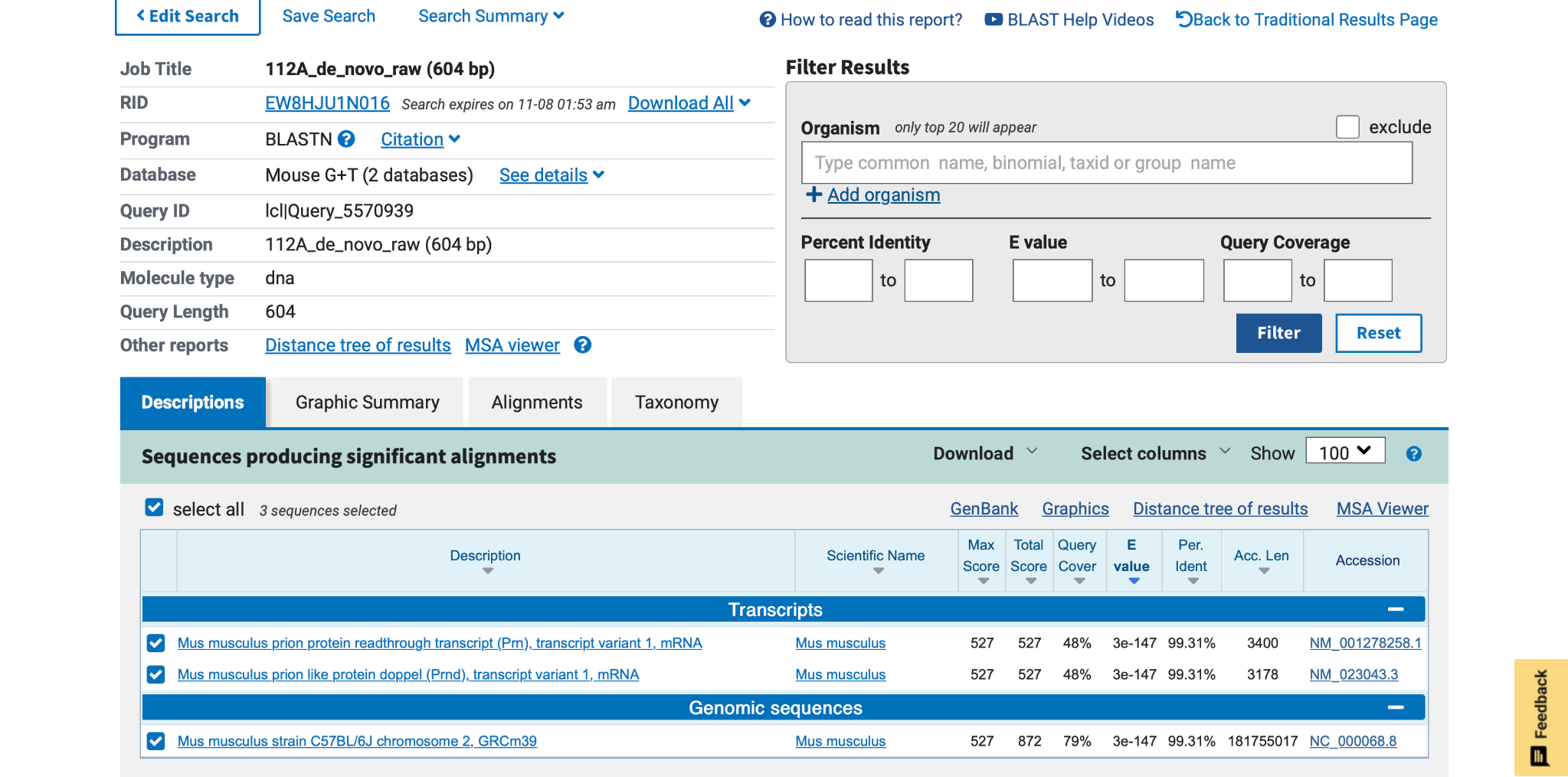

### image(20261106-185957).png

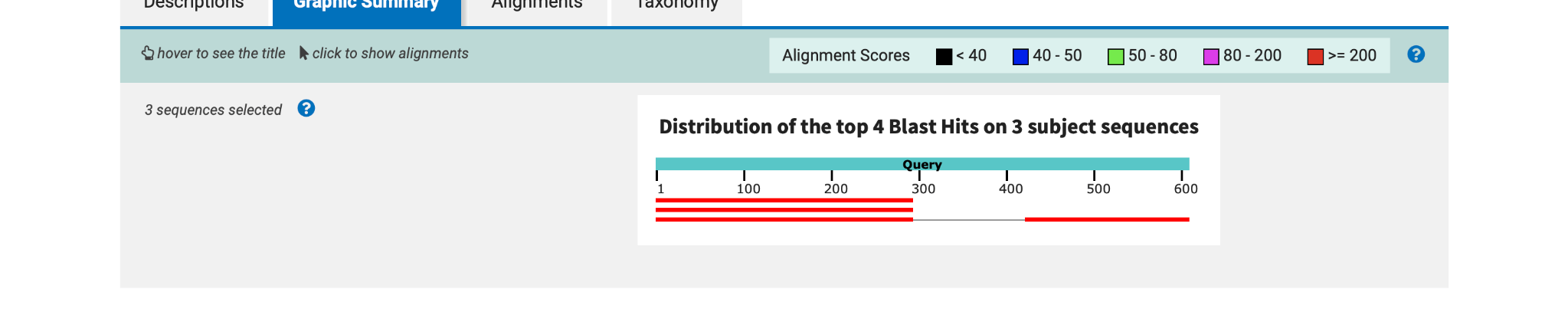

### image(20261106-190030).png

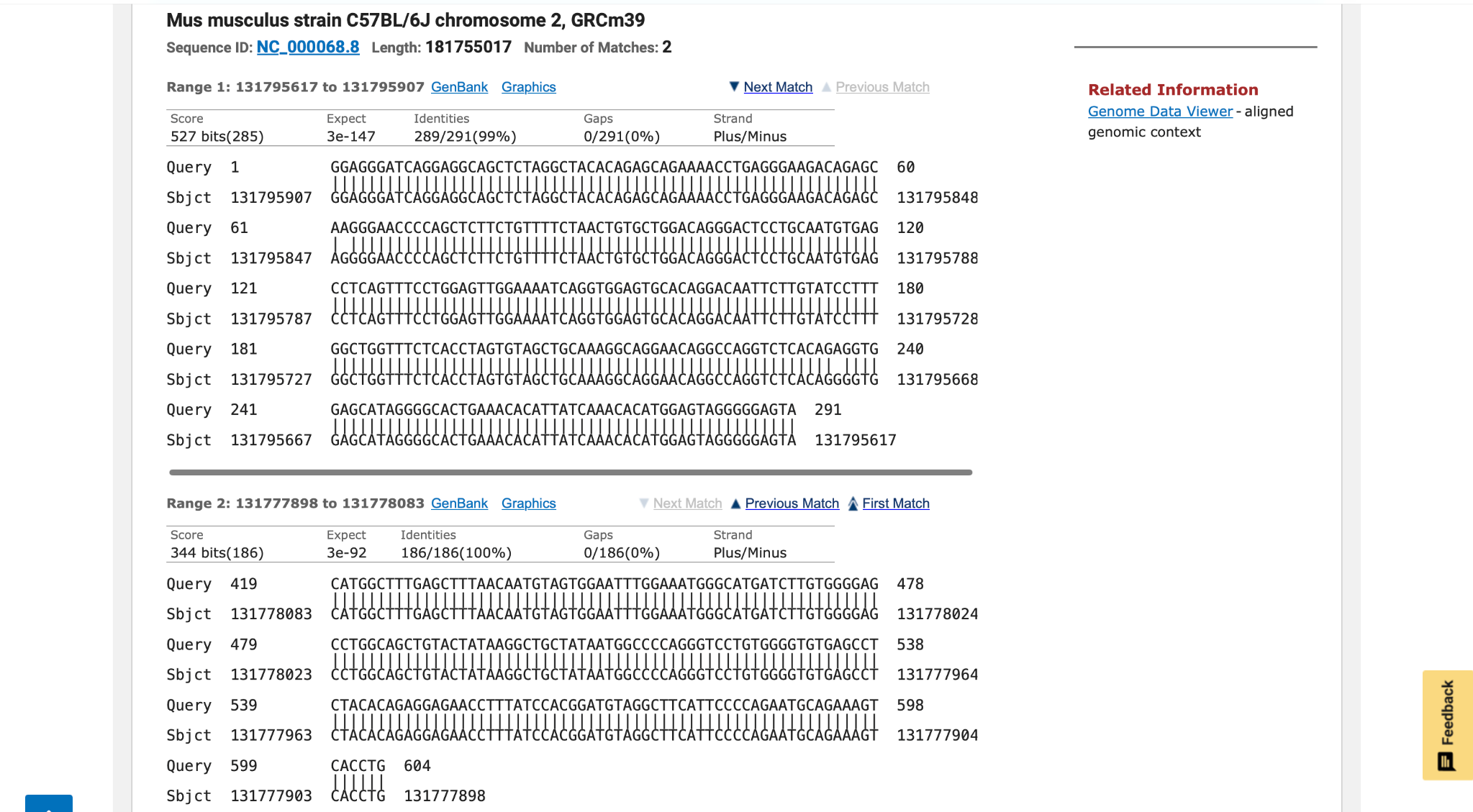

### image(20261106-190616).png

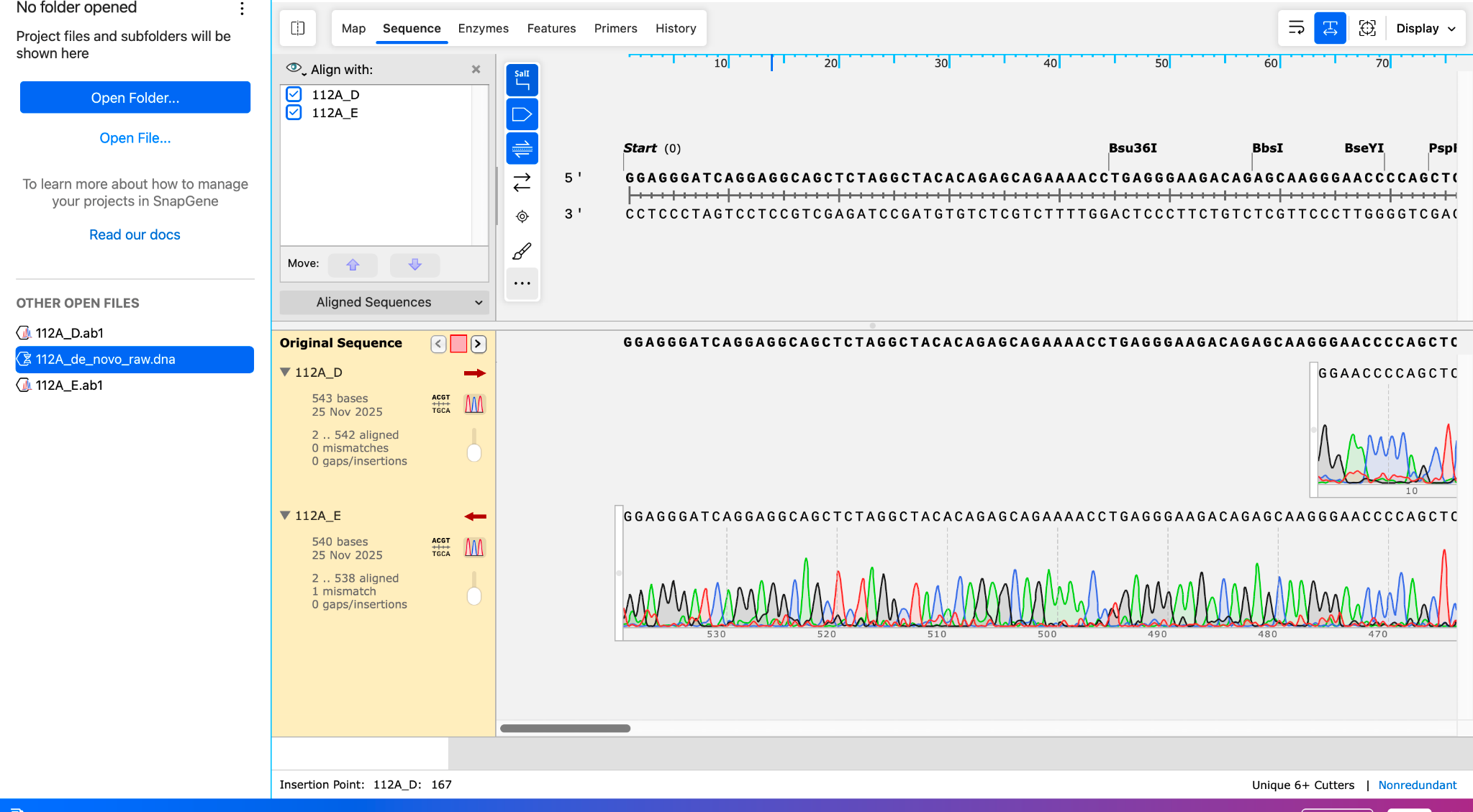

### image(20261106-190634).png

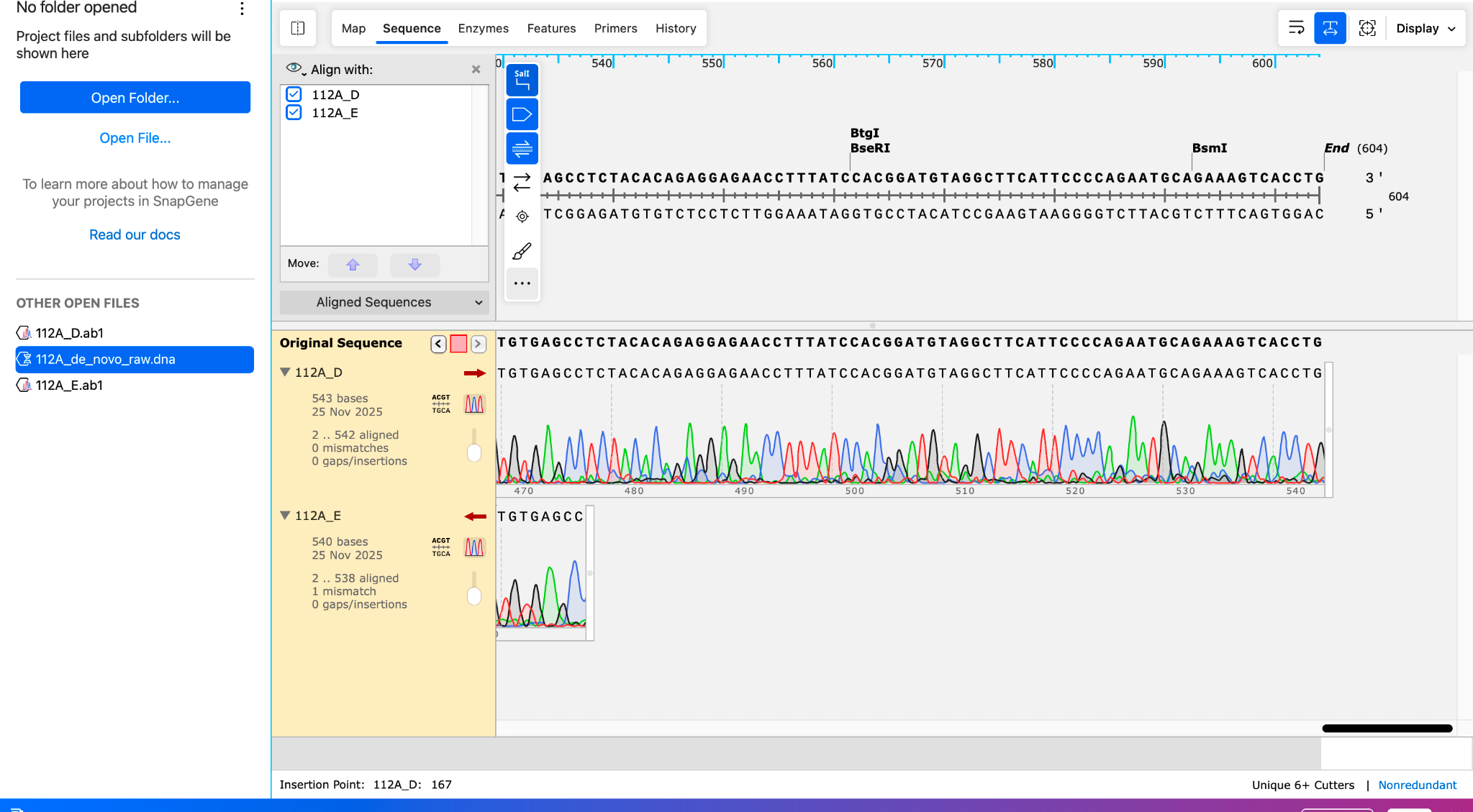

### image(20261106-190842).png

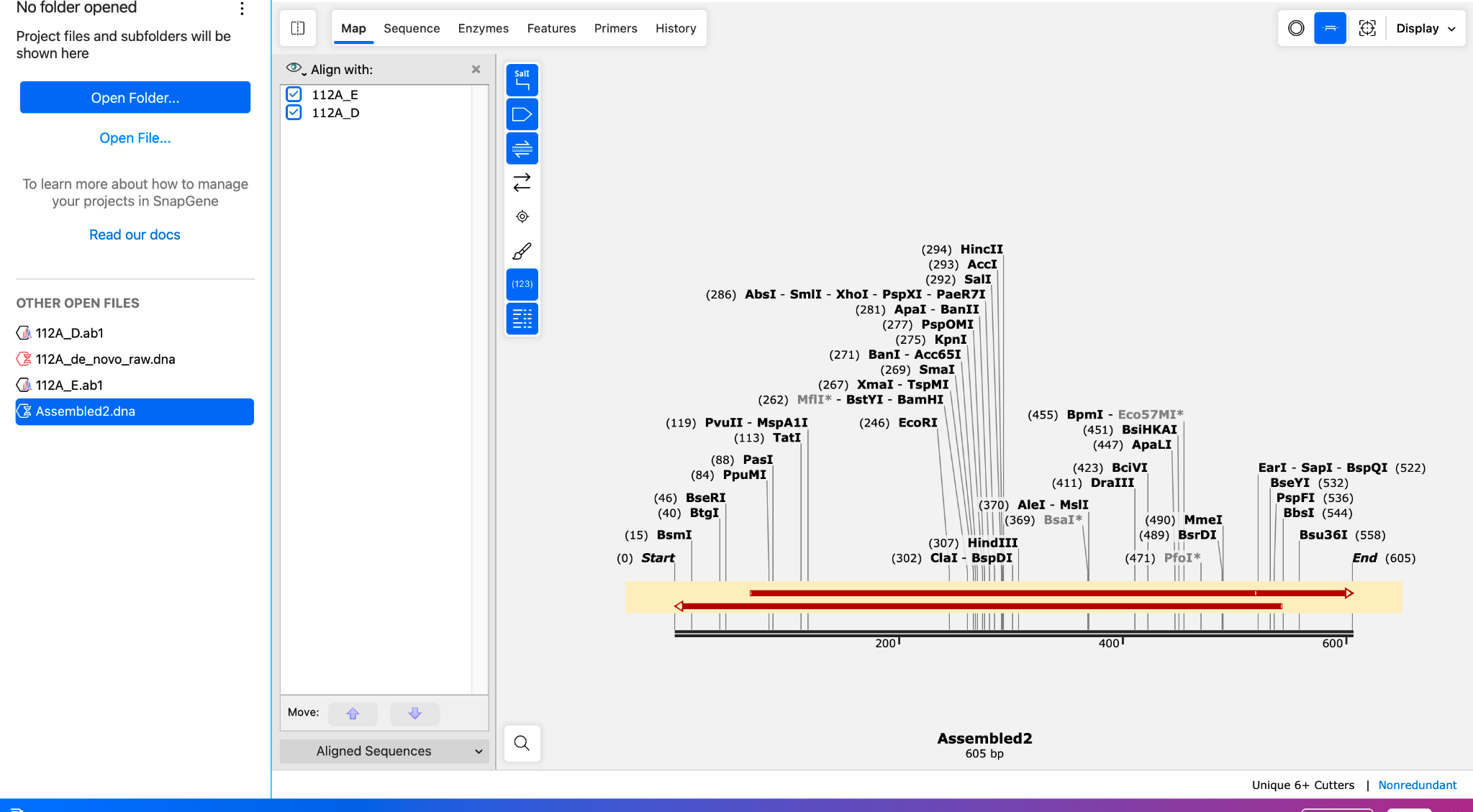

### image(20261106-192015).png

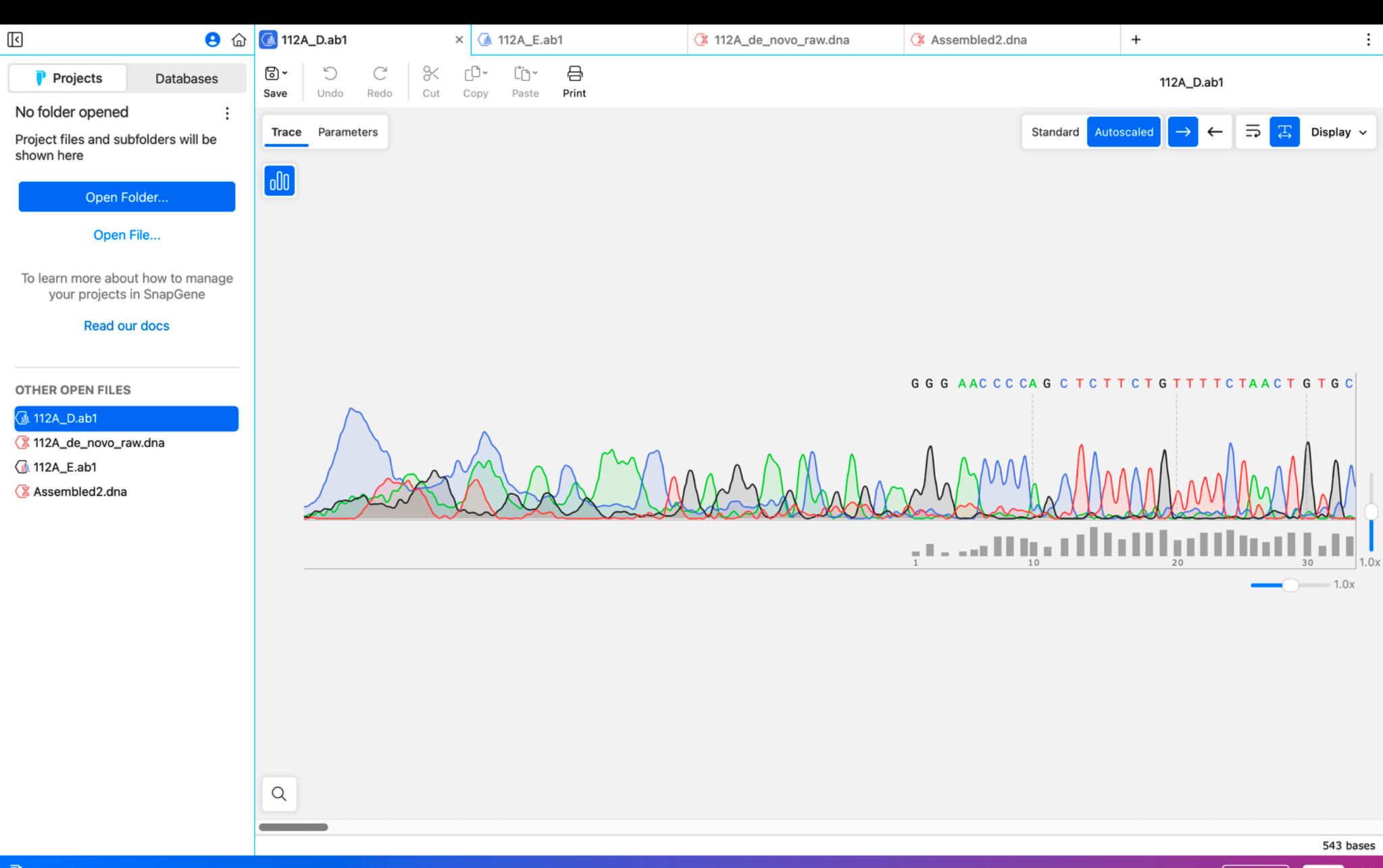

### image(20261106-192130).png

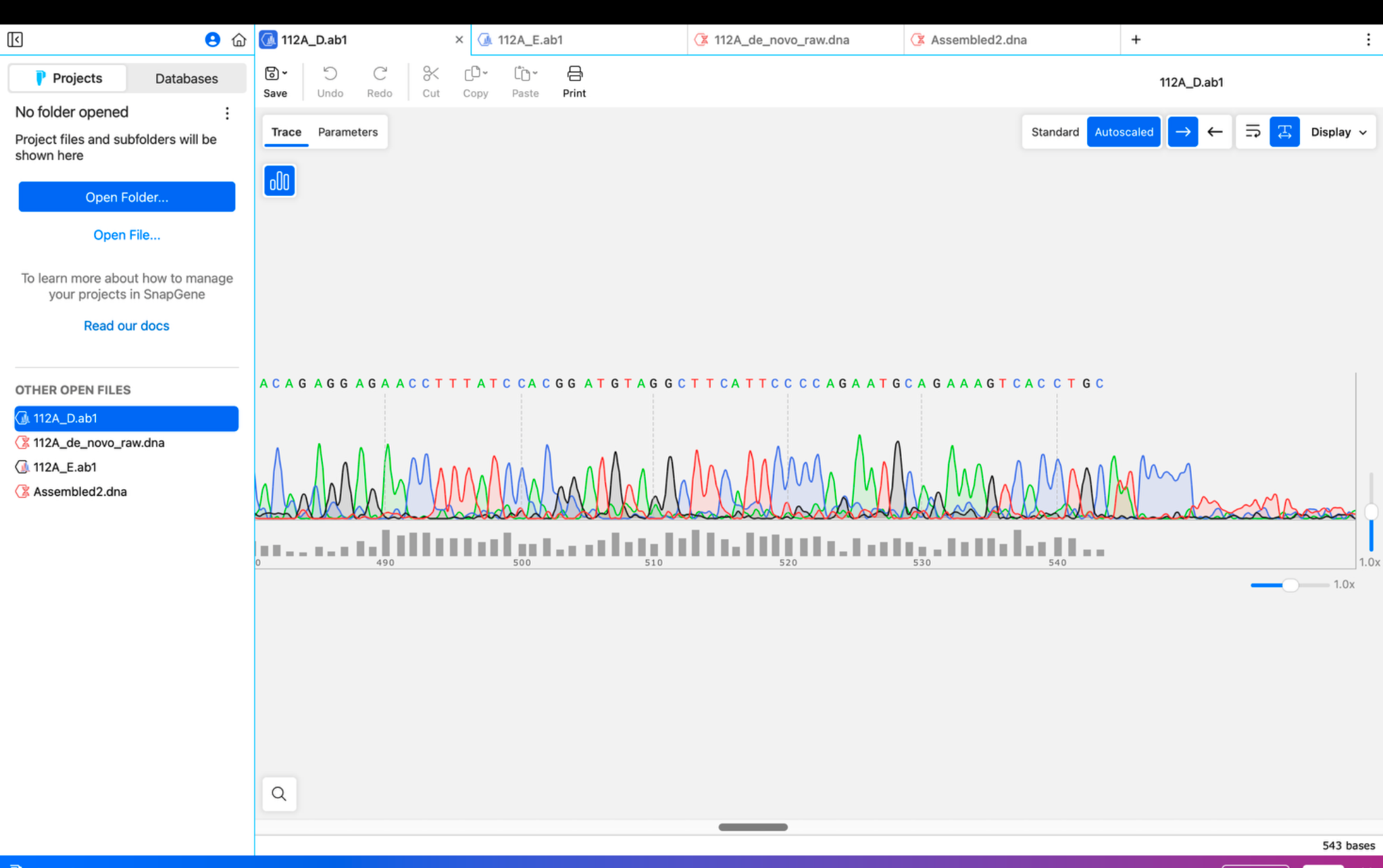

### image(20261106-192909).png

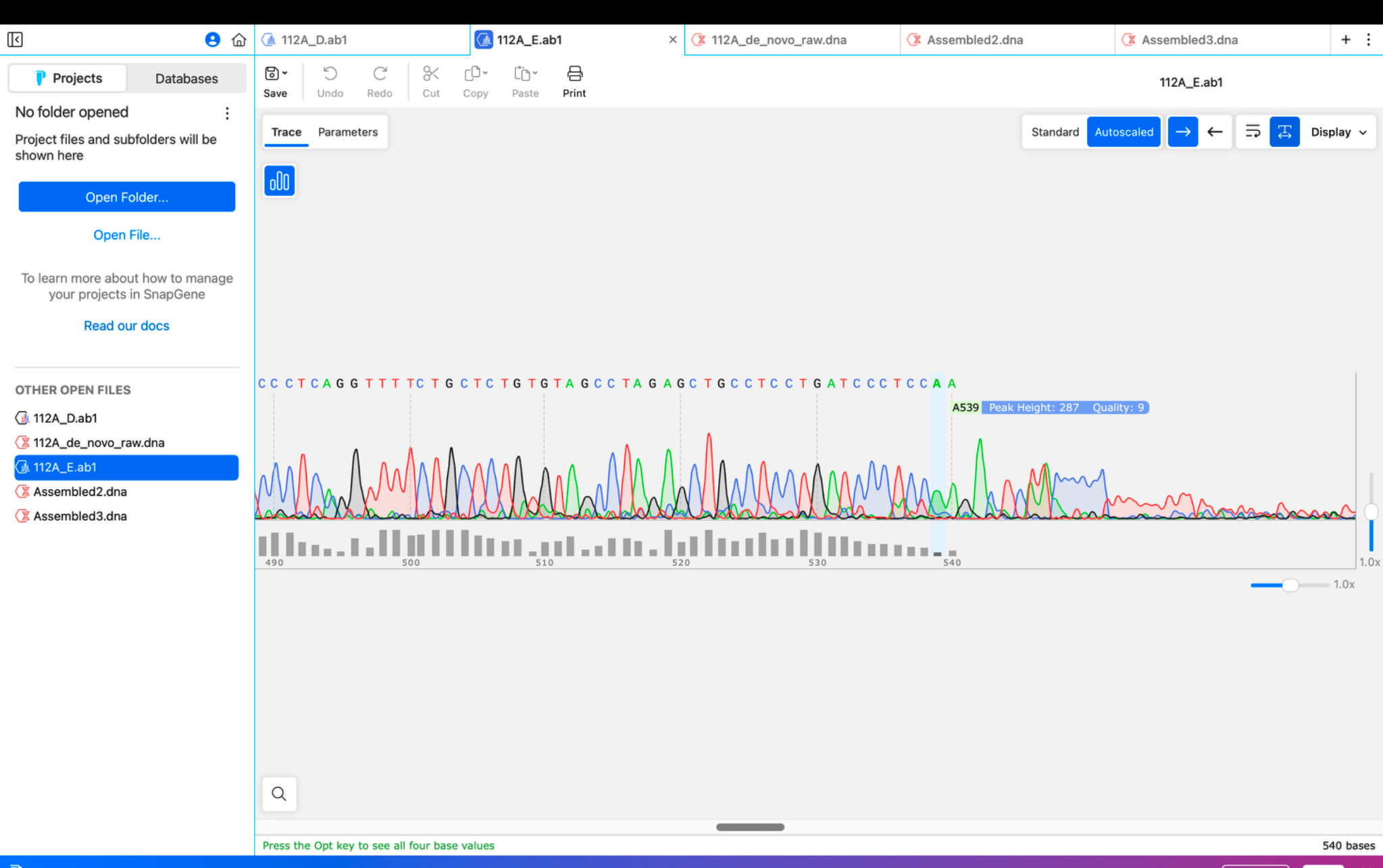

### image(20261106-192918).png

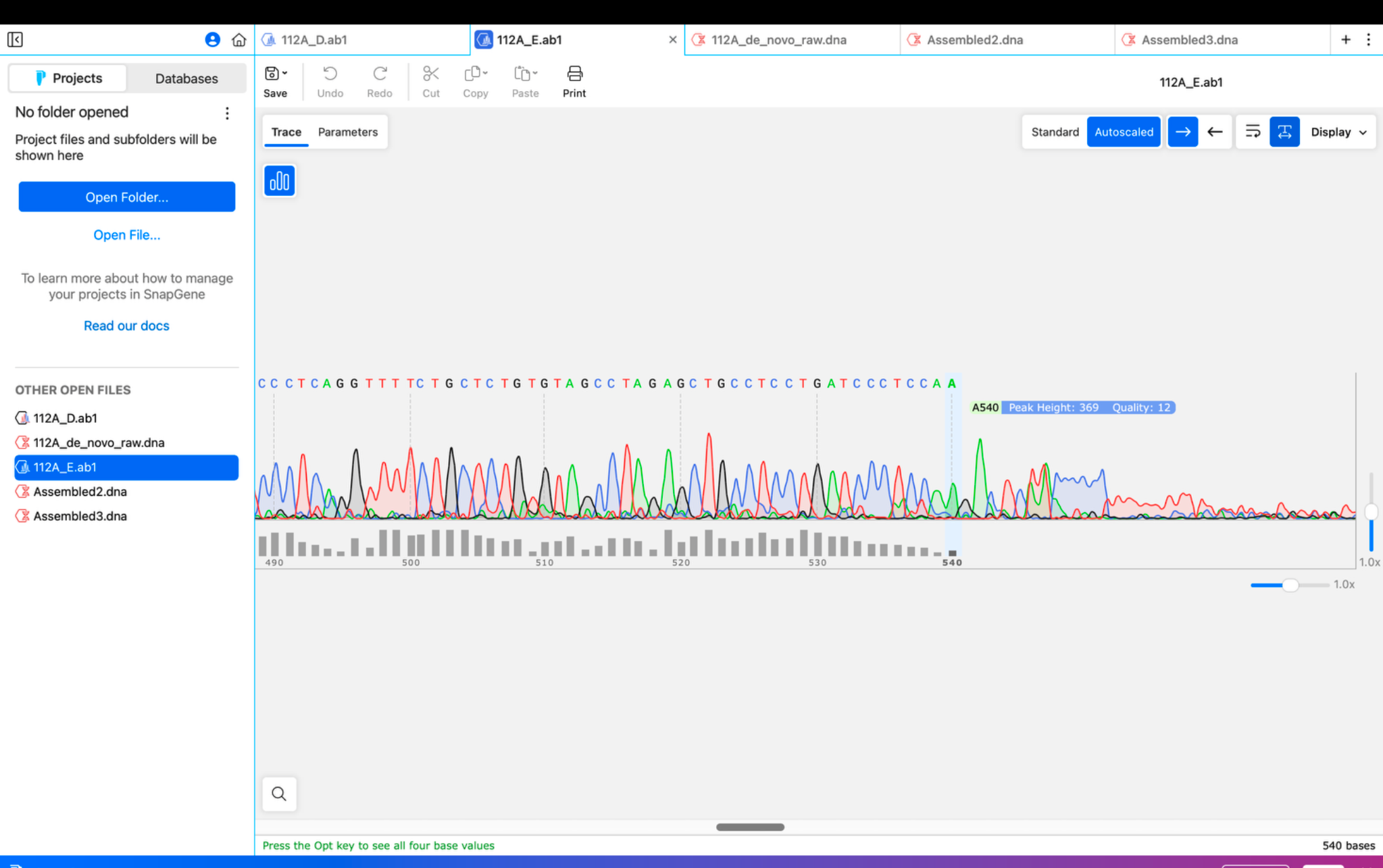

### image(20261106-193200).png

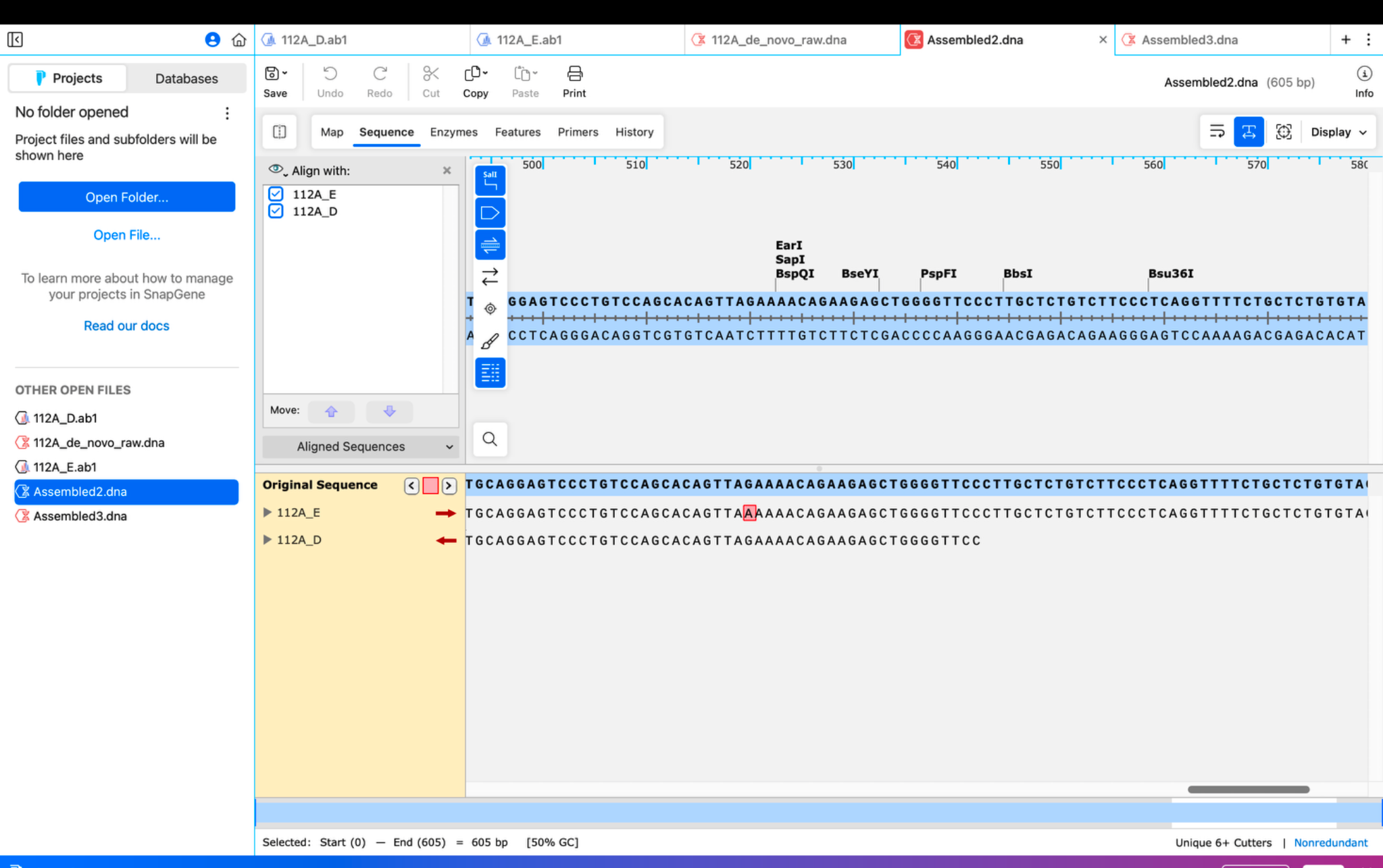

### image(20261106-193210).png

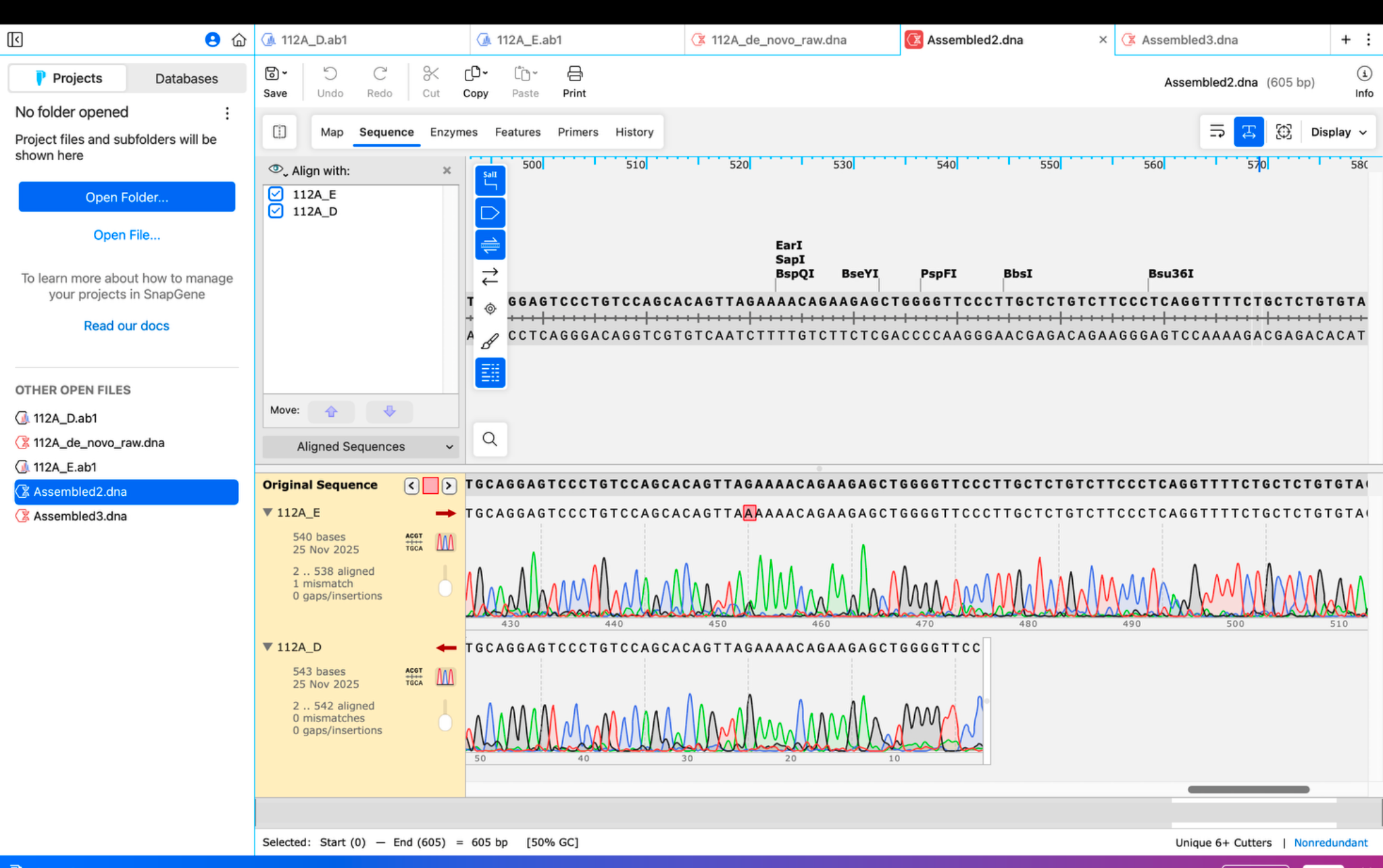

### image(20261106-193545).png

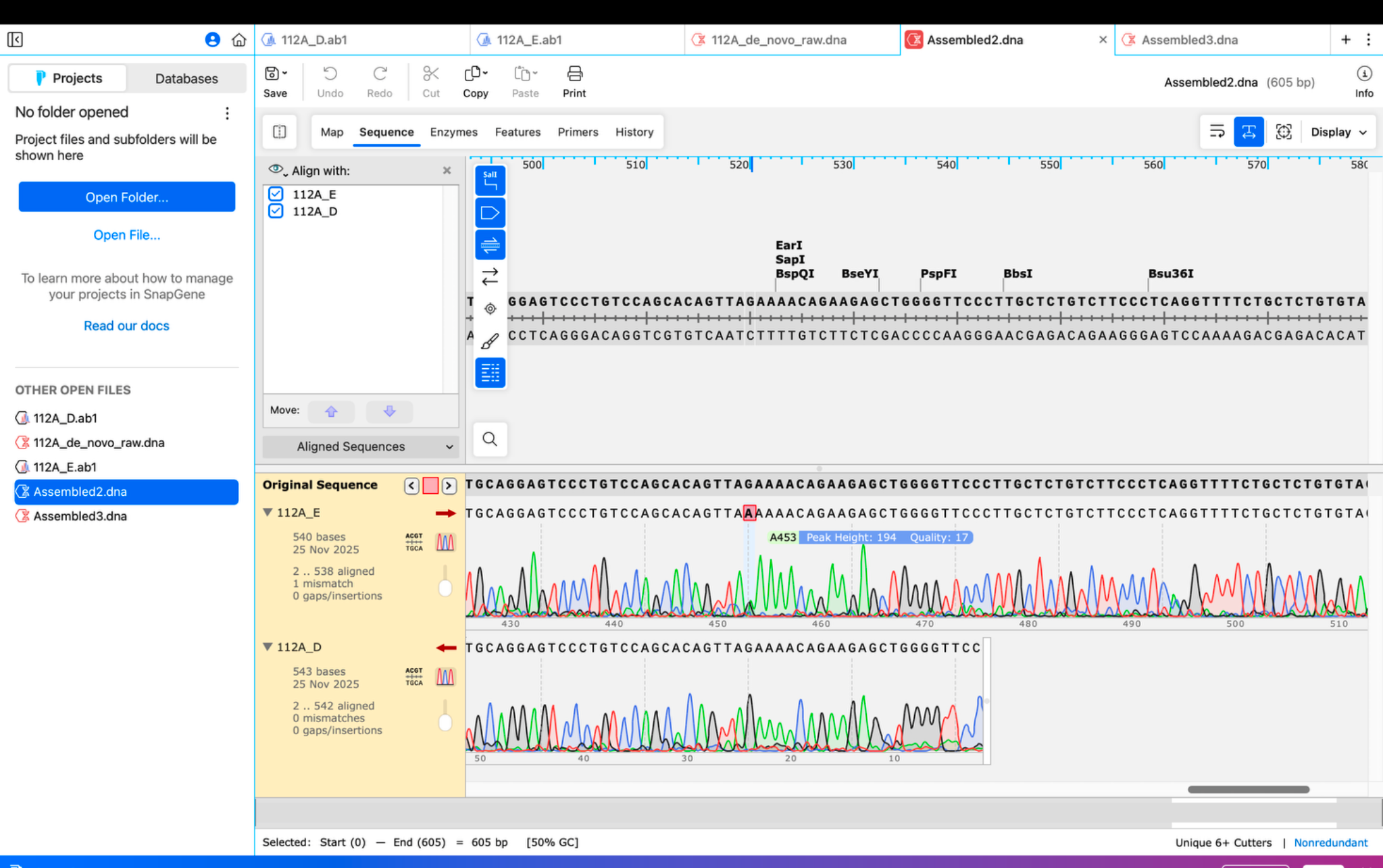

### image(20261106-193555).png

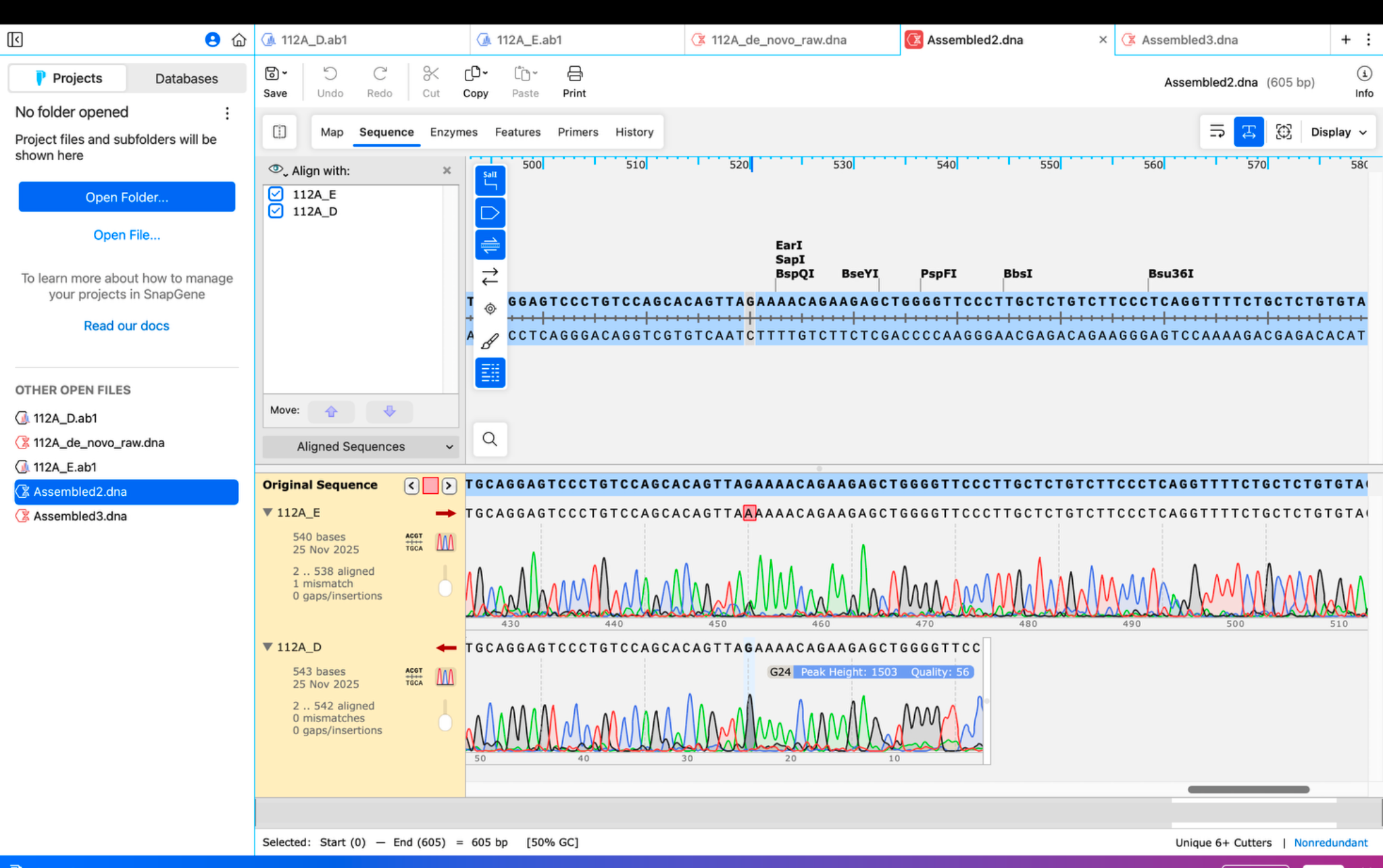

### image(20261108-173722).jpeg

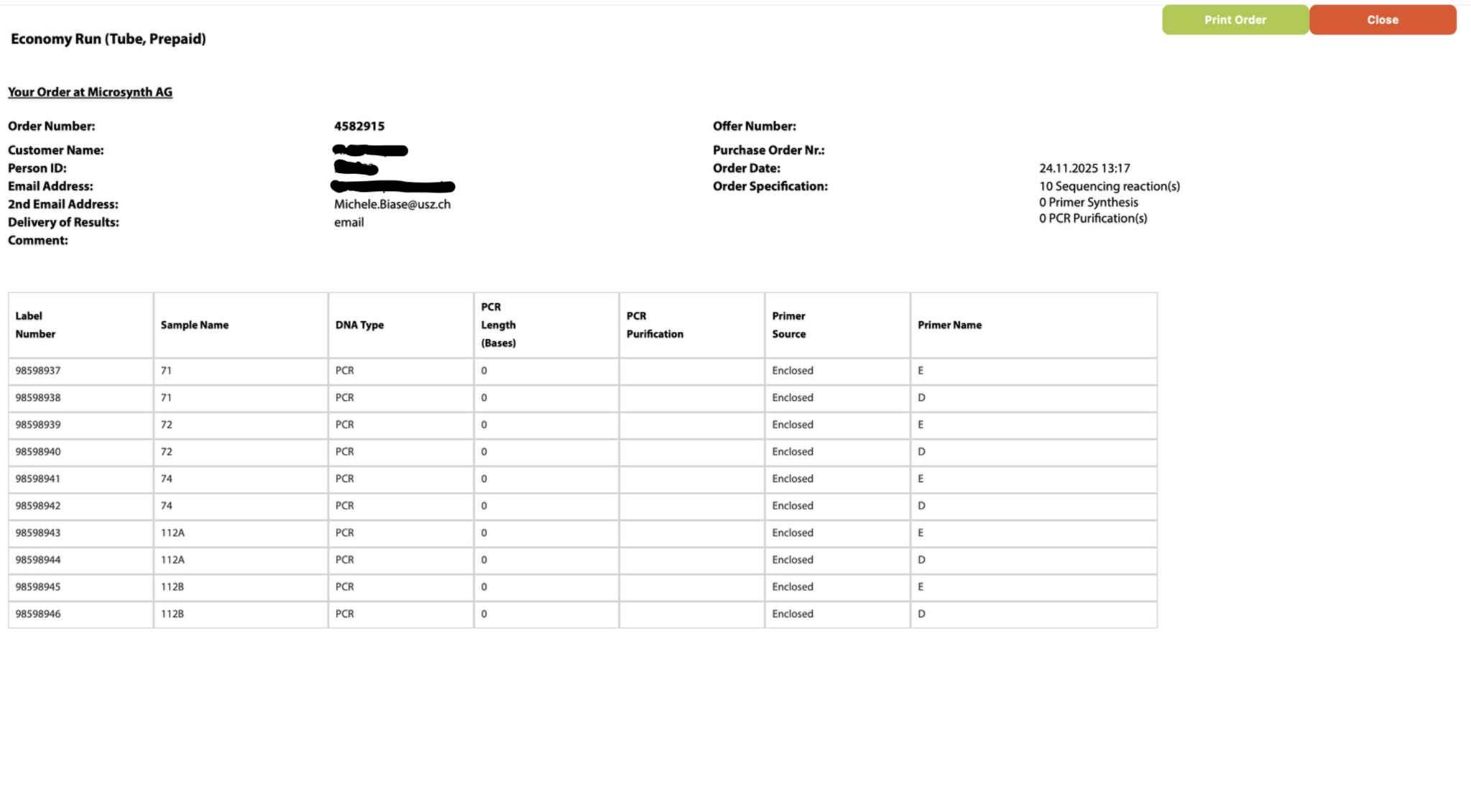

### image(20261108-175836).png

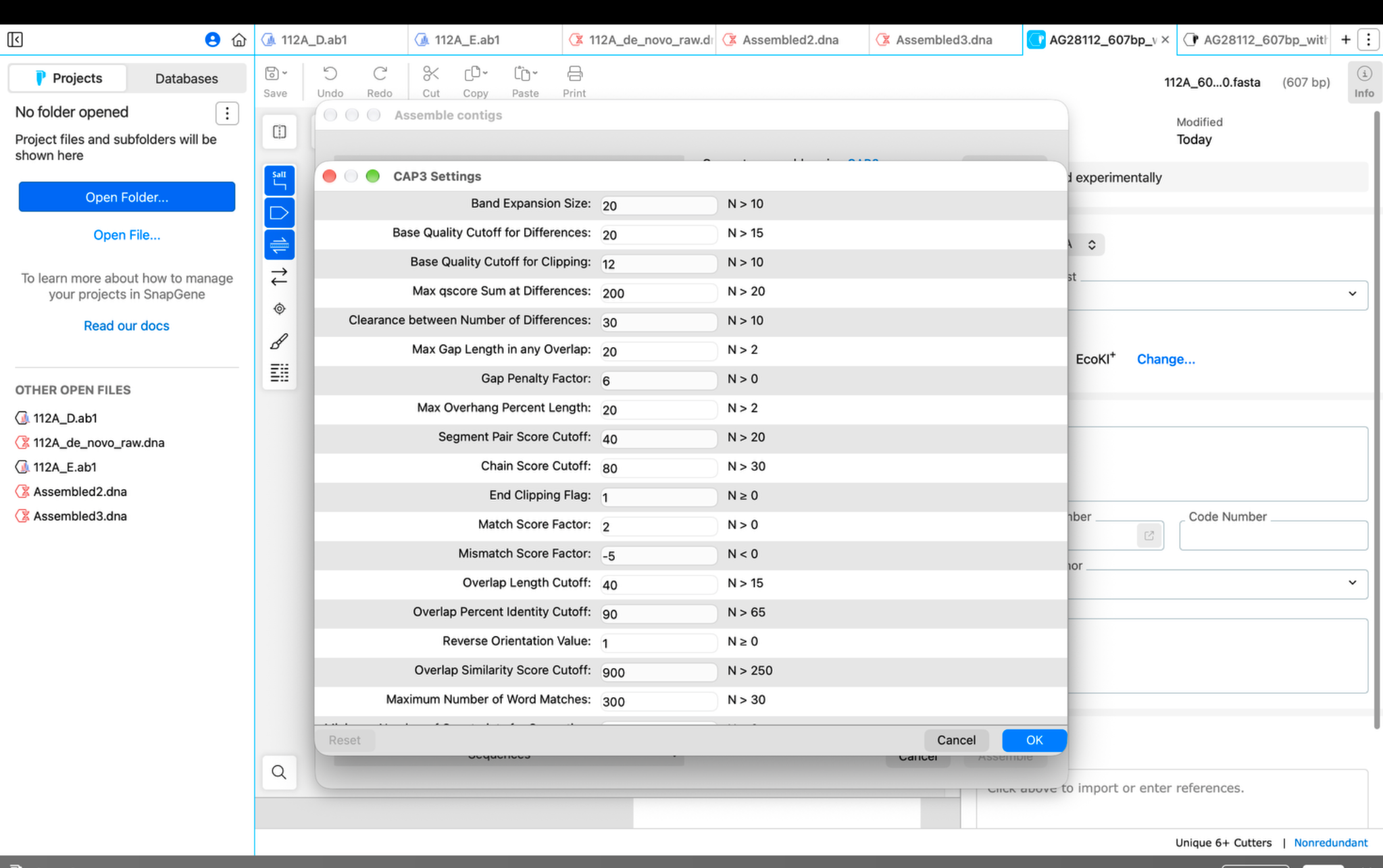

### image(20261108-175845).png

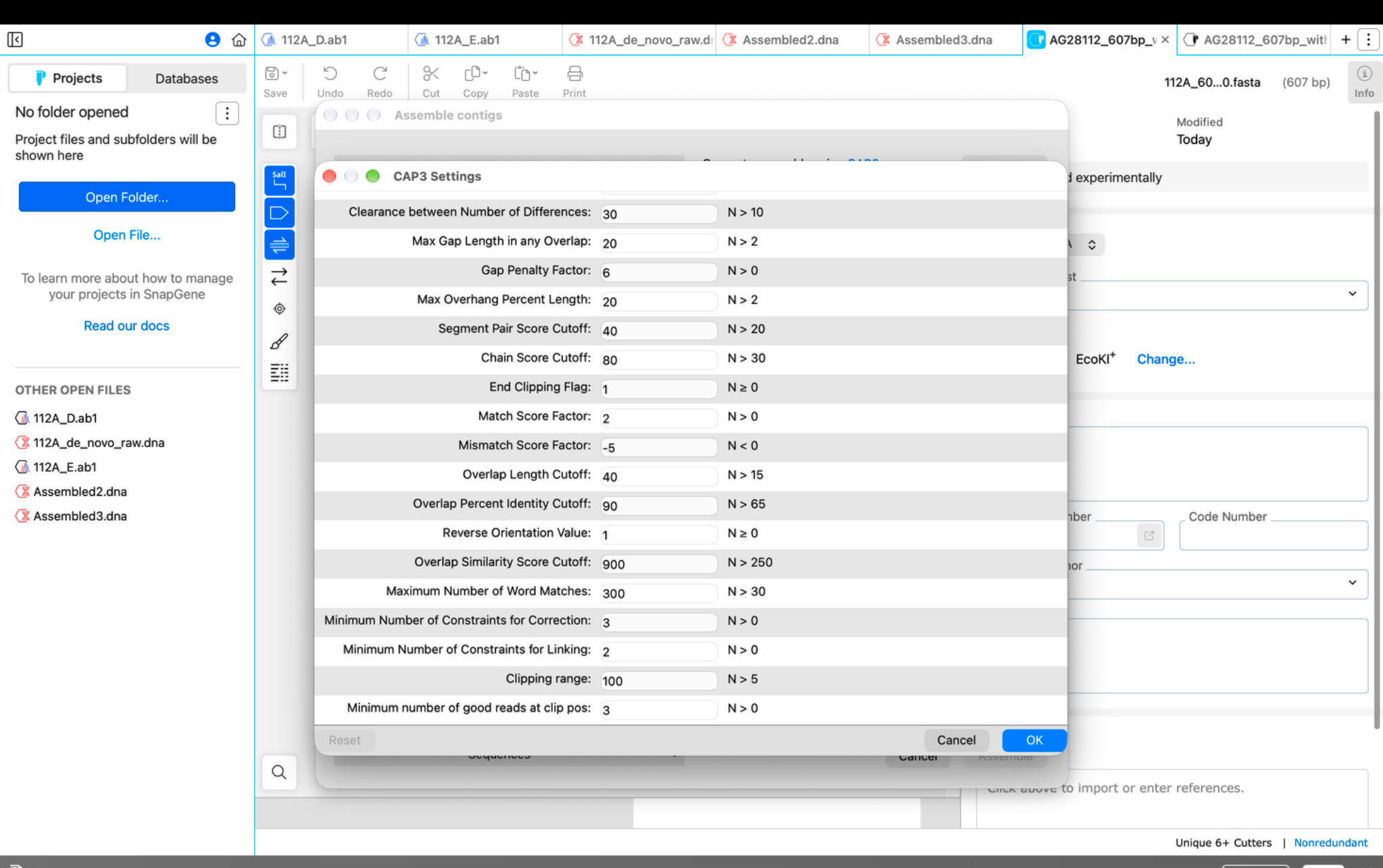

### image(20261108-180017).png

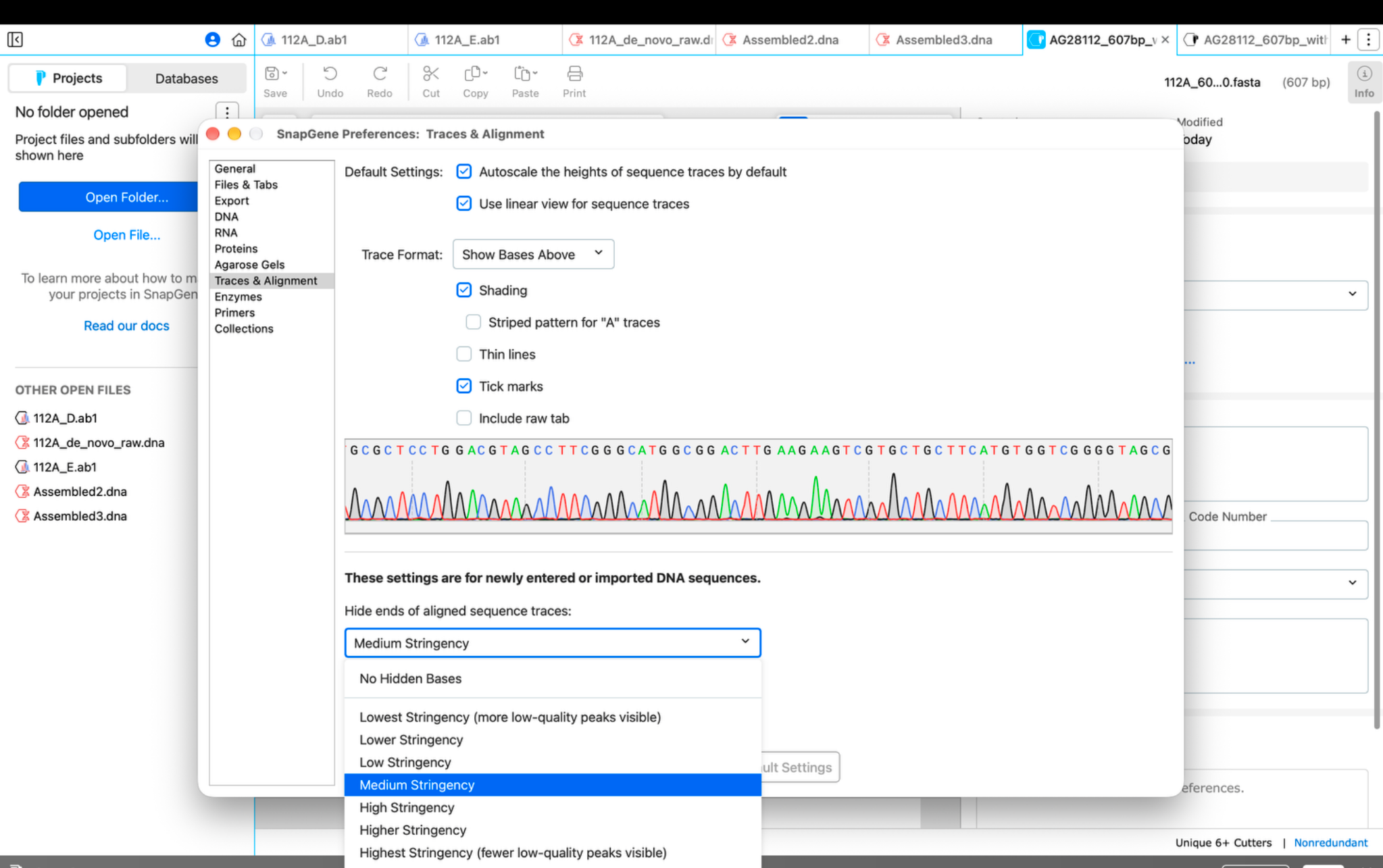

### image(20261108-182339).png

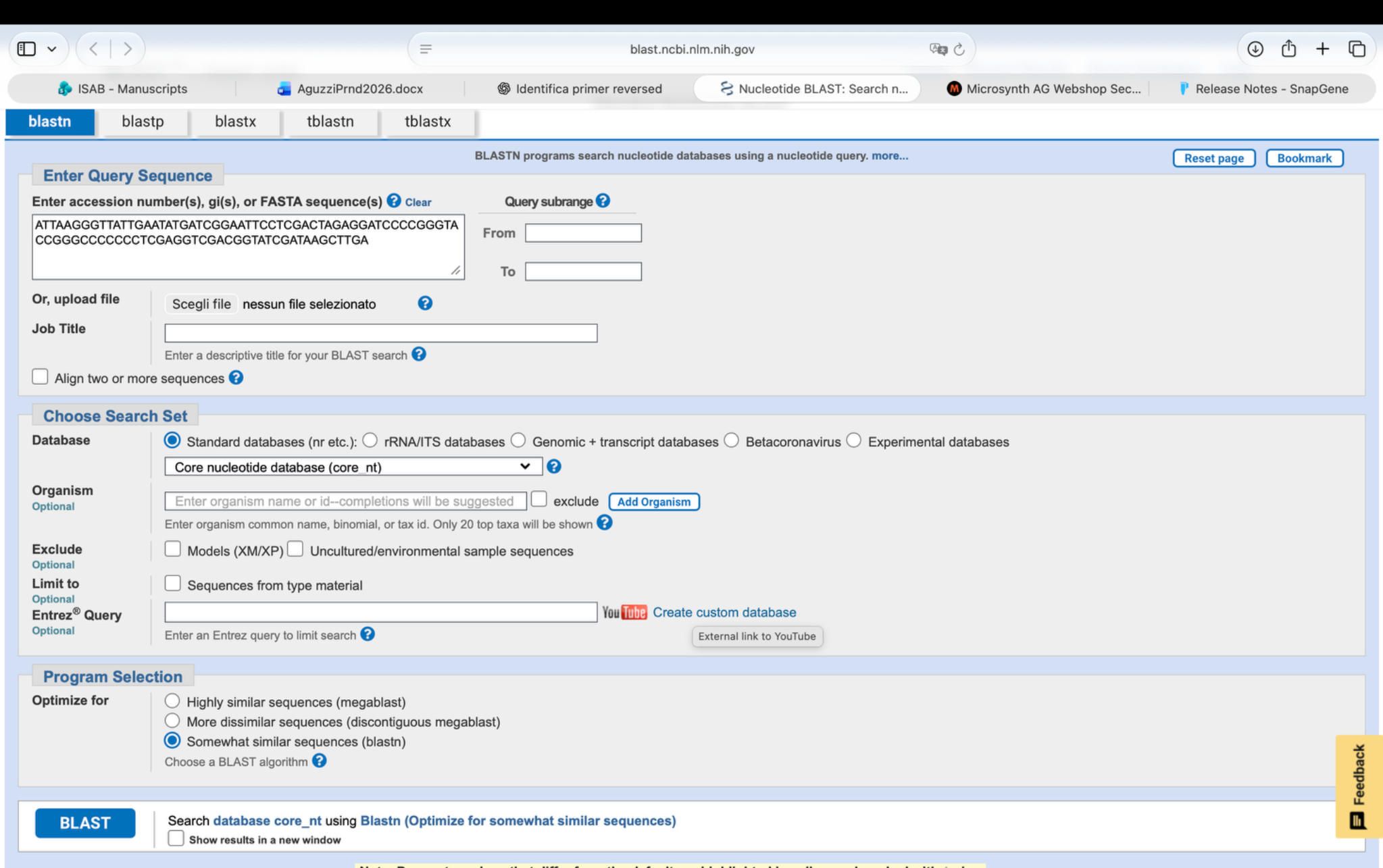

### image(20261108-182348).png

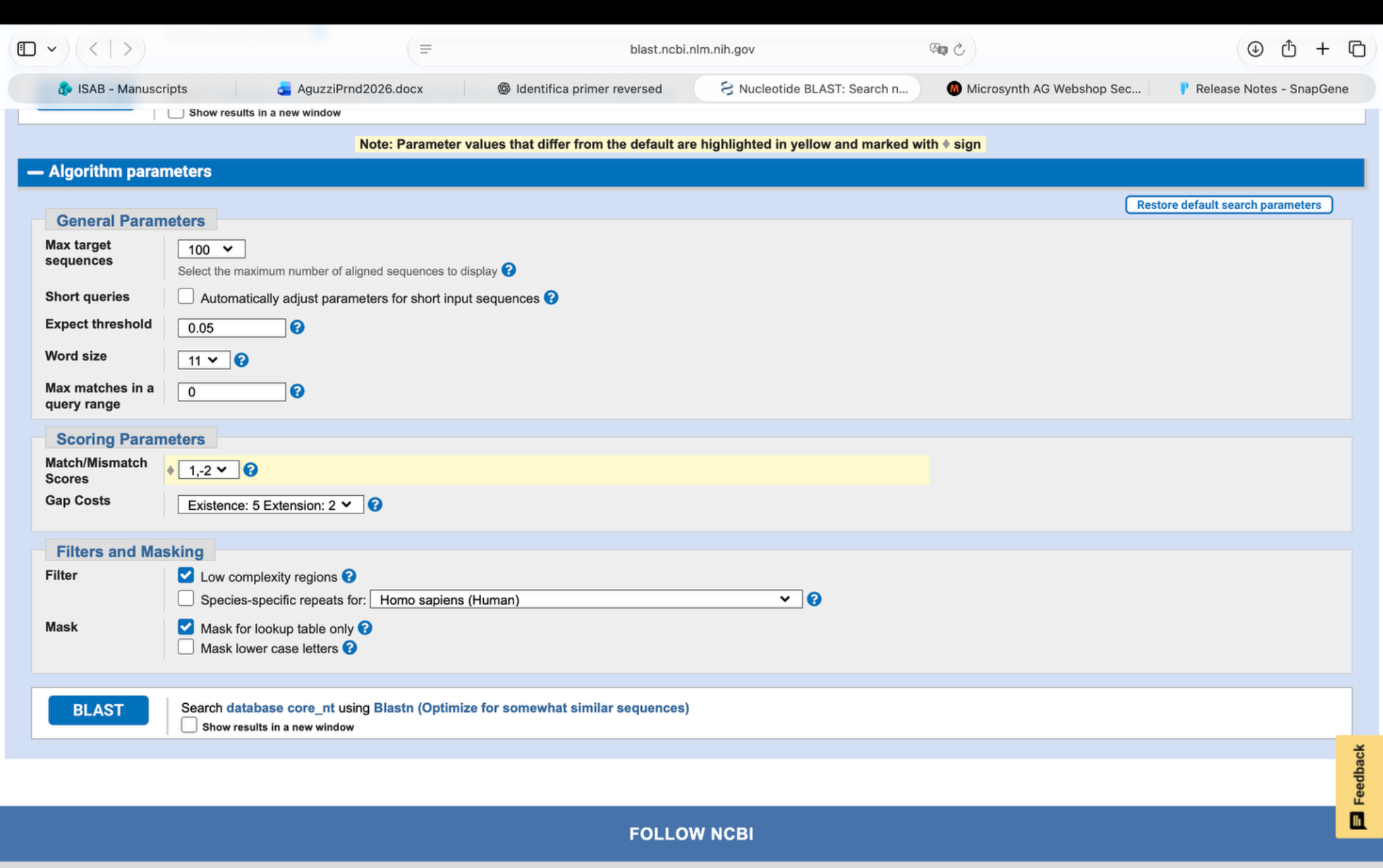

### image(20261108-182630).png

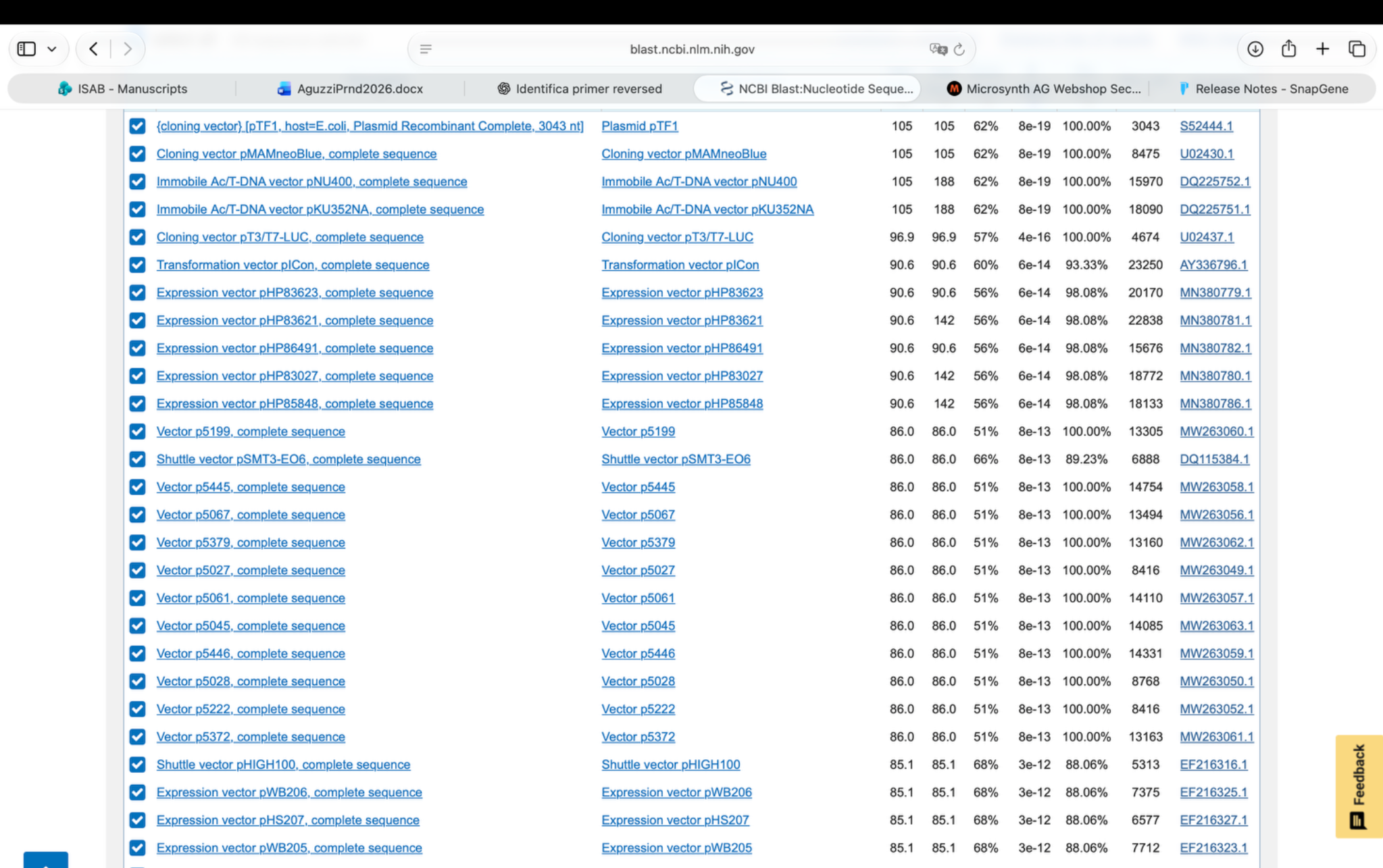

### image(20261108-182653).png

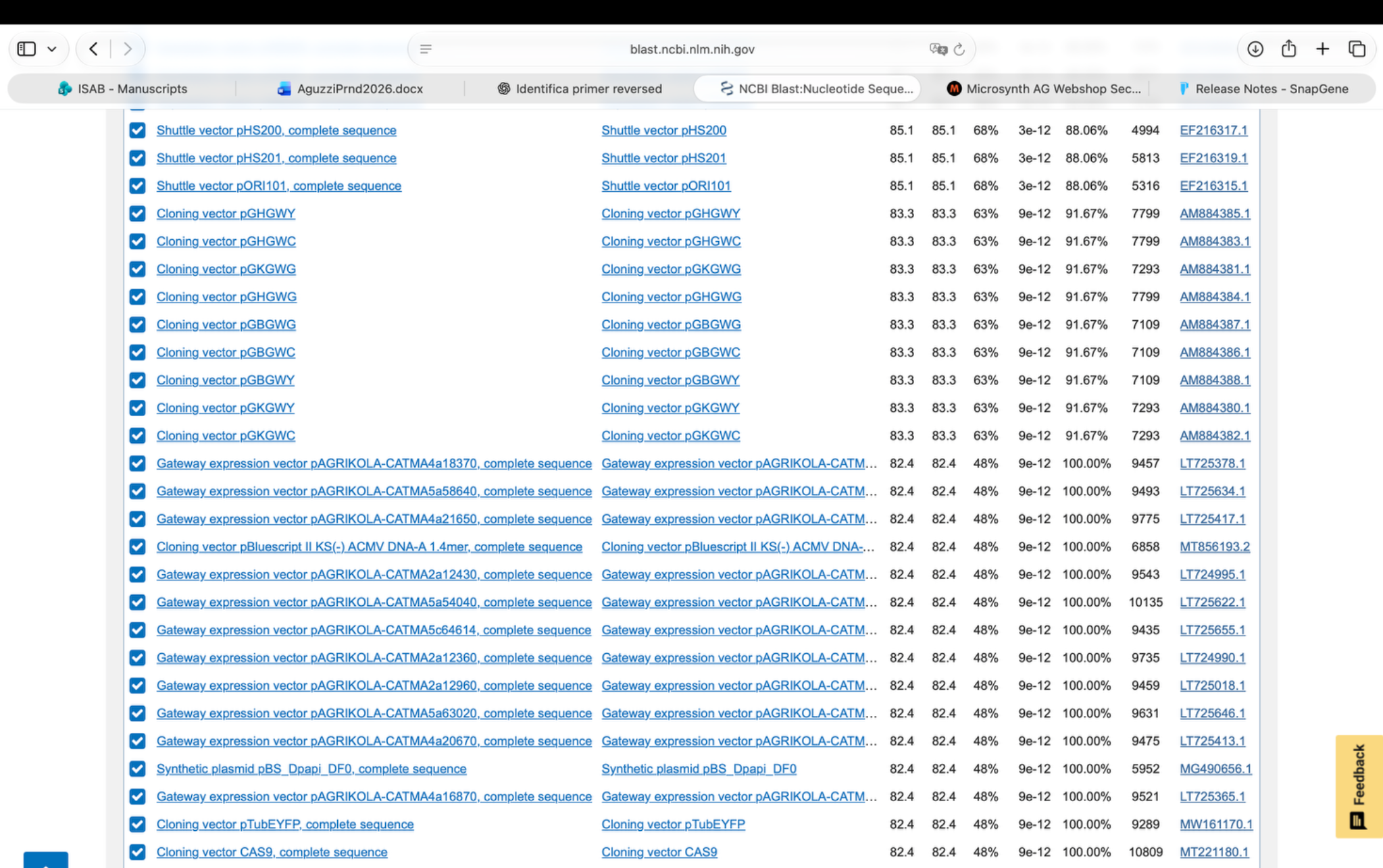

### image(20261108-182745).png

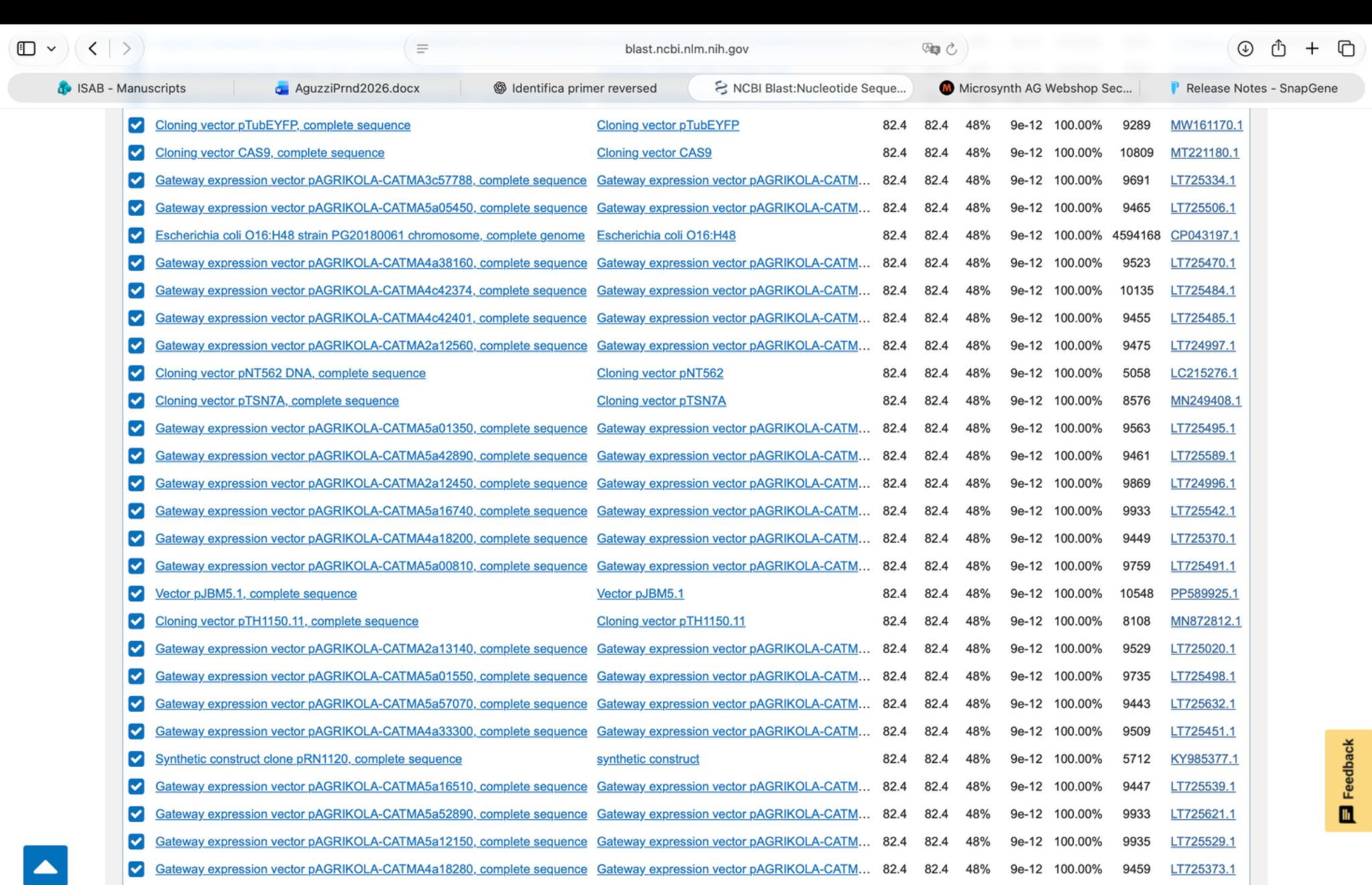

### image(20261108-182751).png

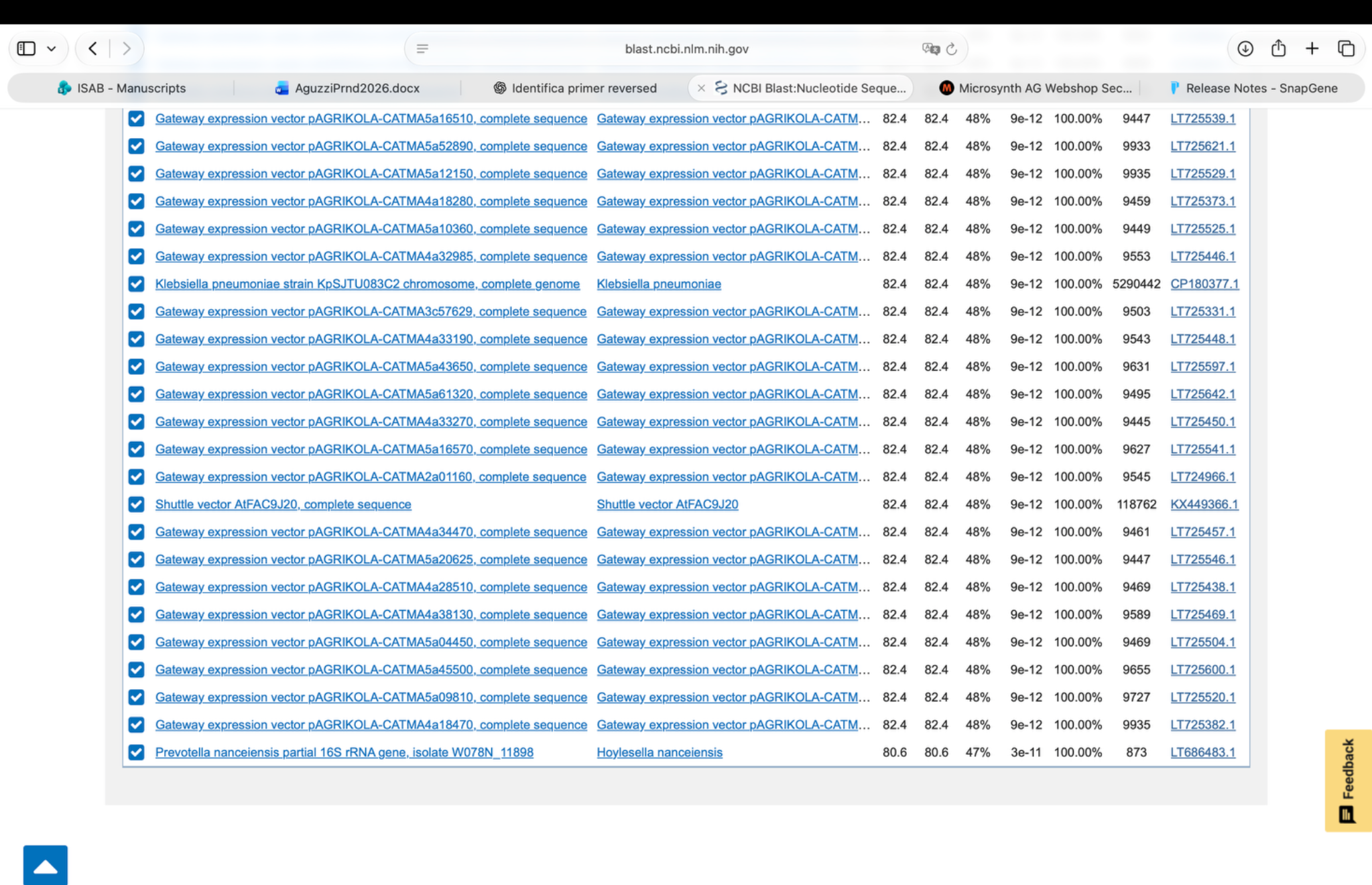

### image(20261108-182814).png

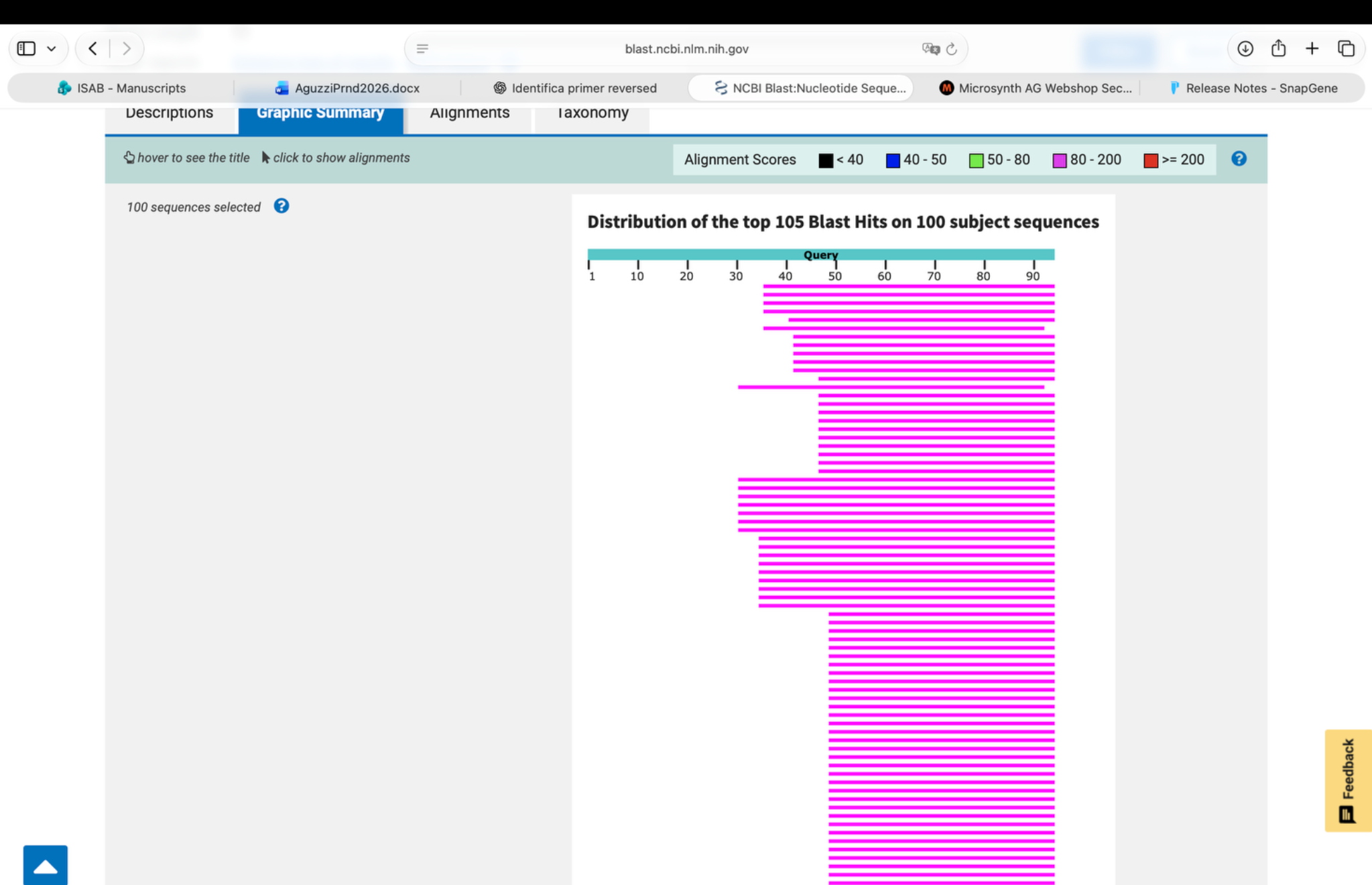

### image(20261108-182821).png

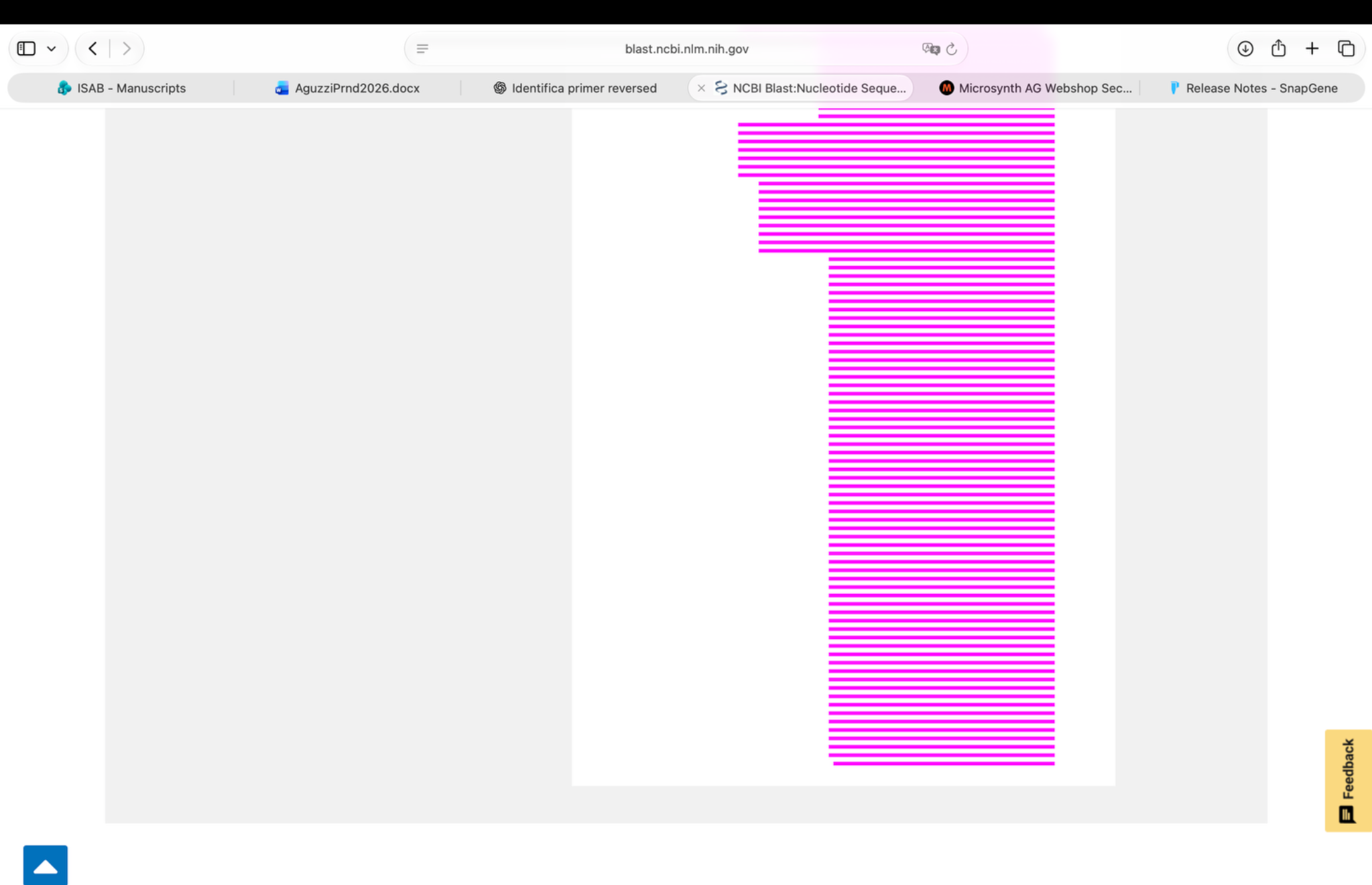

### image(20261108-184119).png

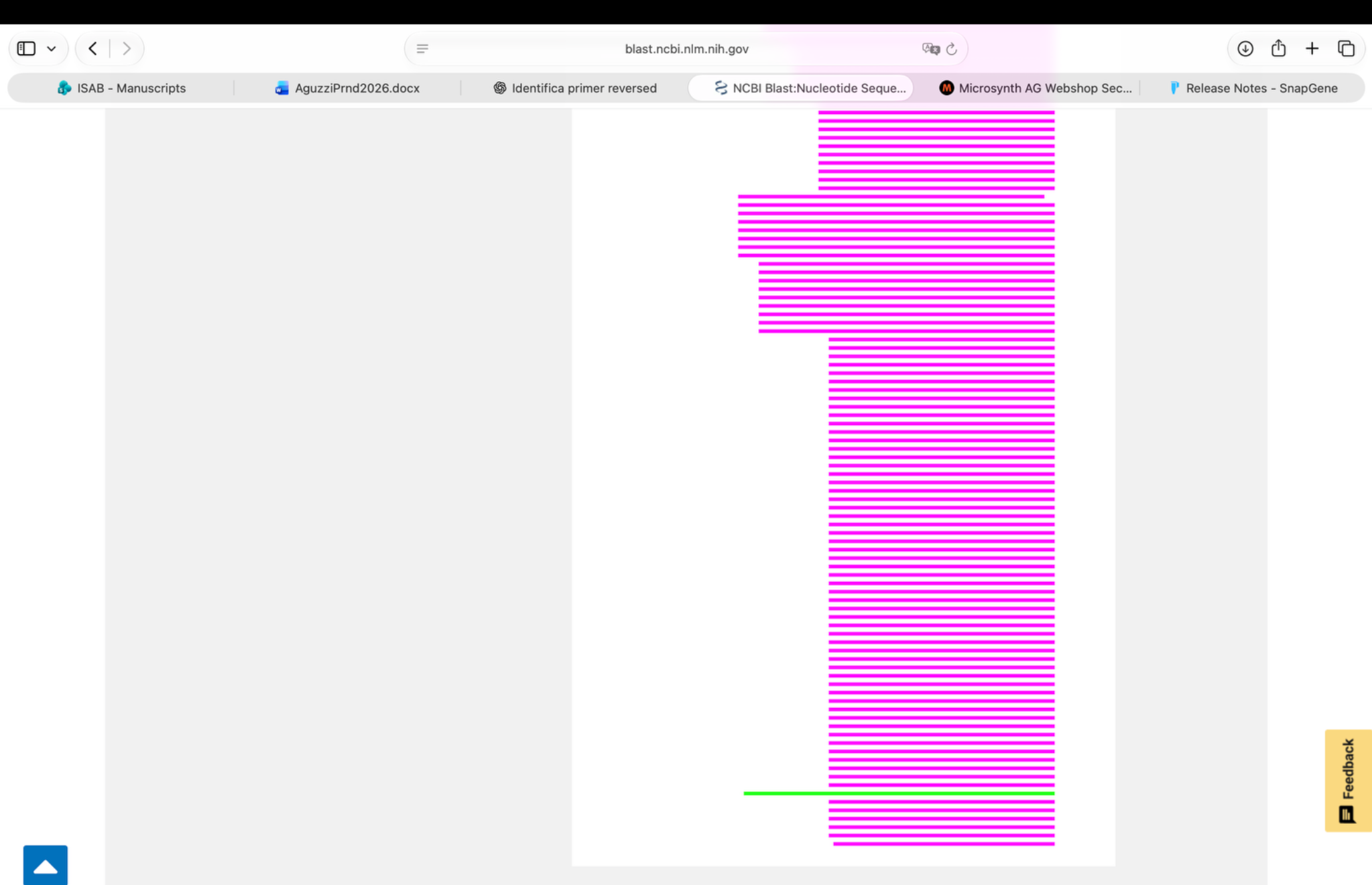

### image(20261108-184243).png

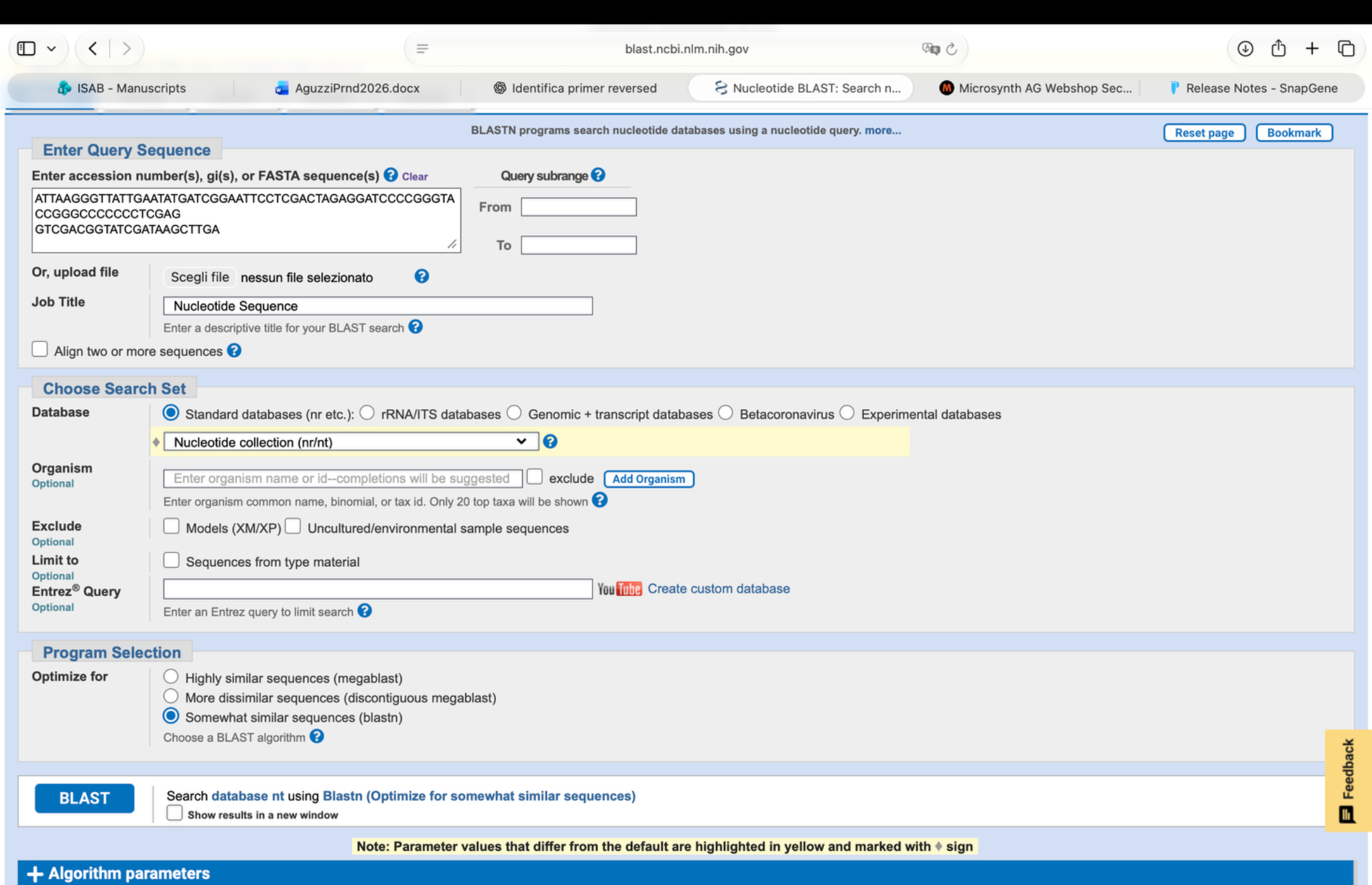

### image(20261108-184252).png

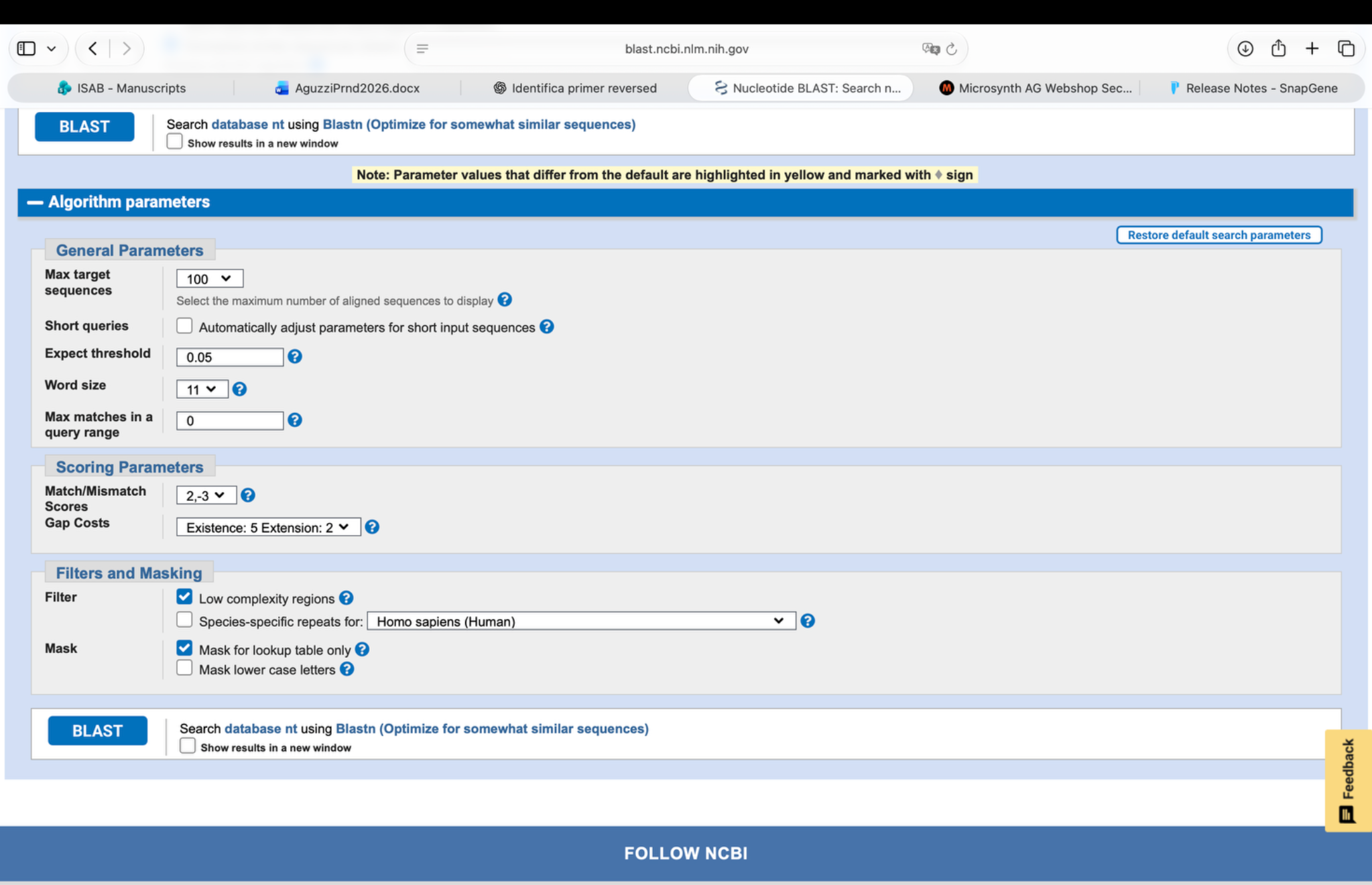

### image(20261108-185027).png

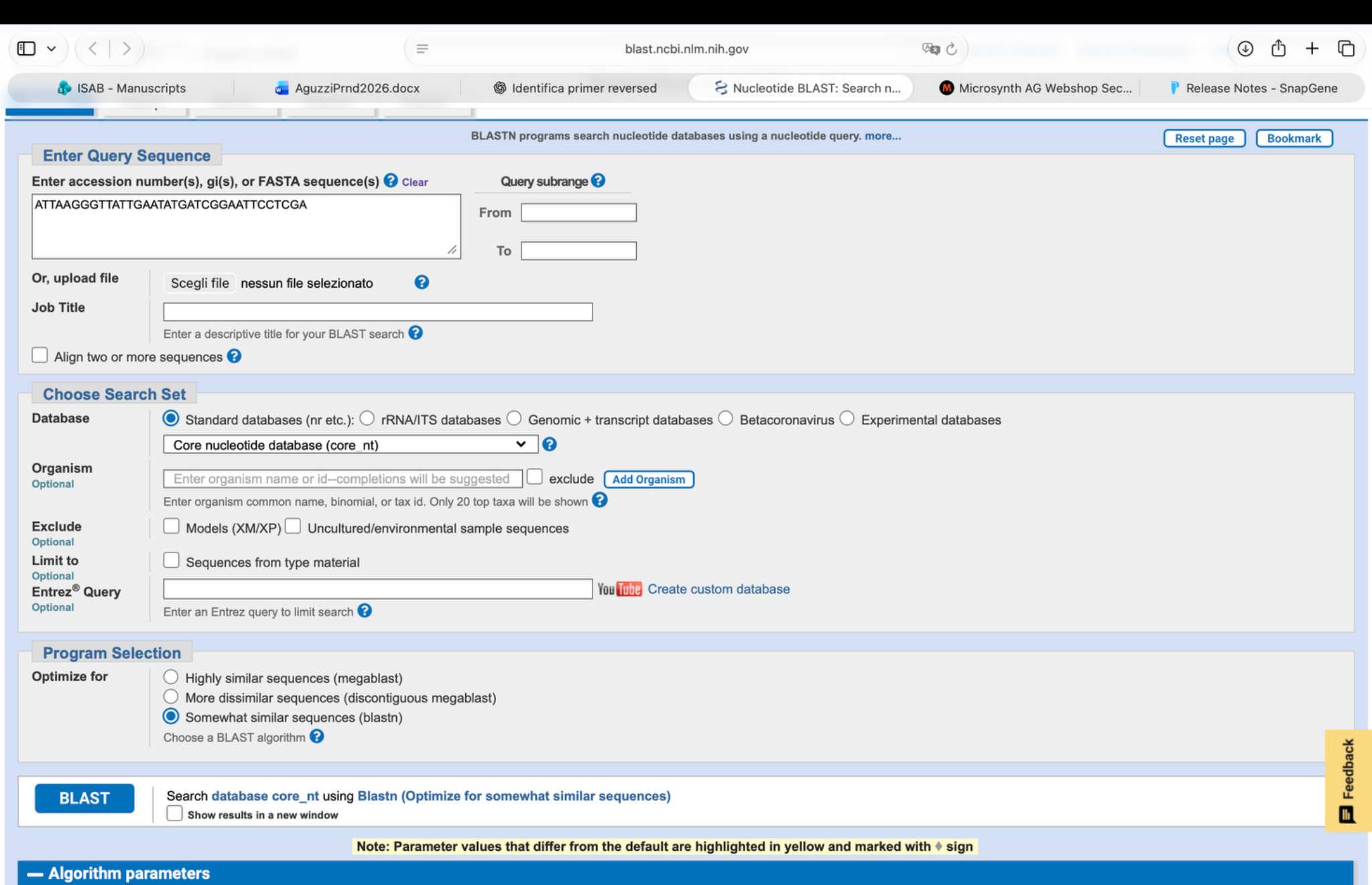

### image(20261108-185038).png

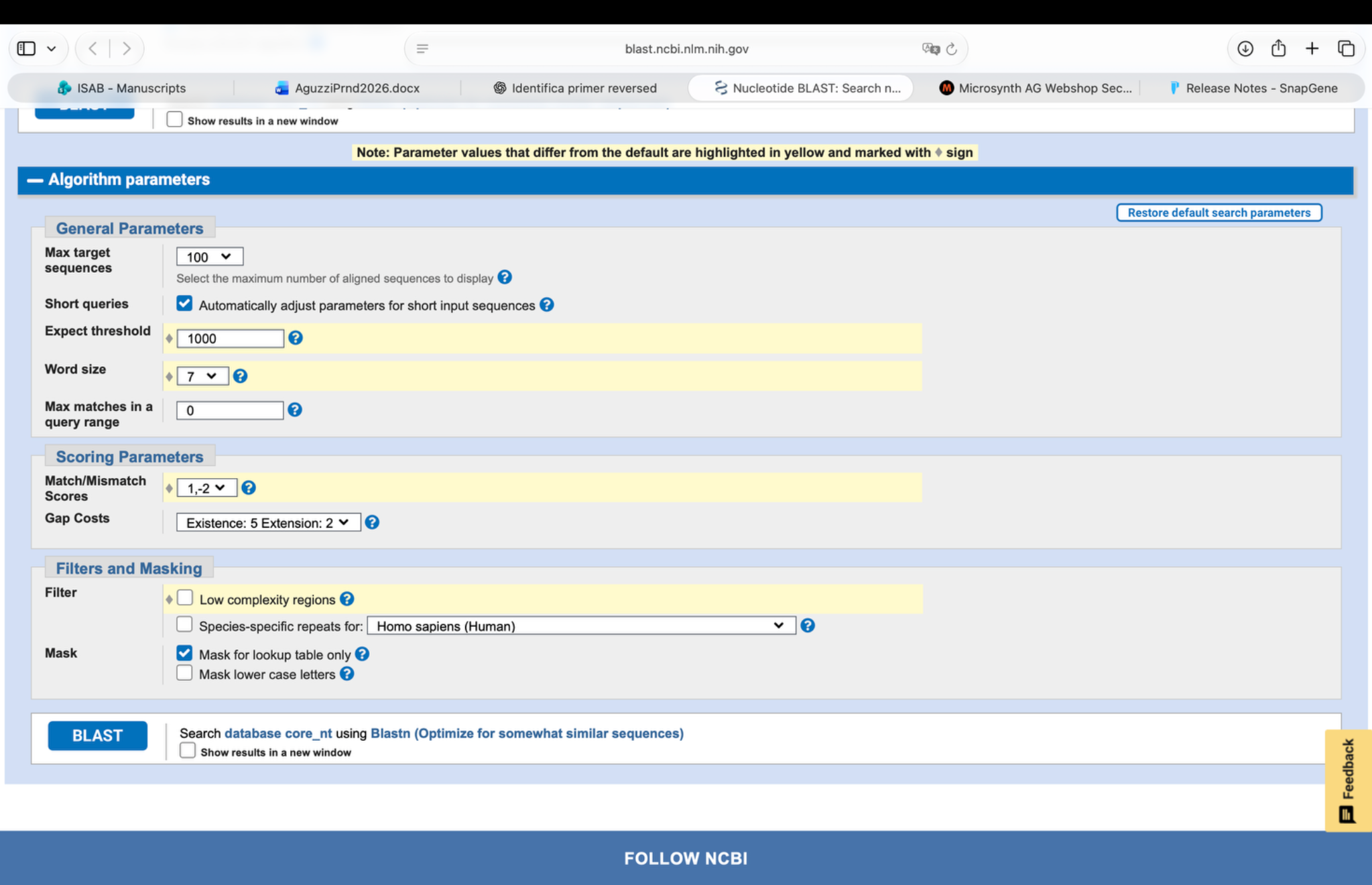

### image(20261108-185518).png

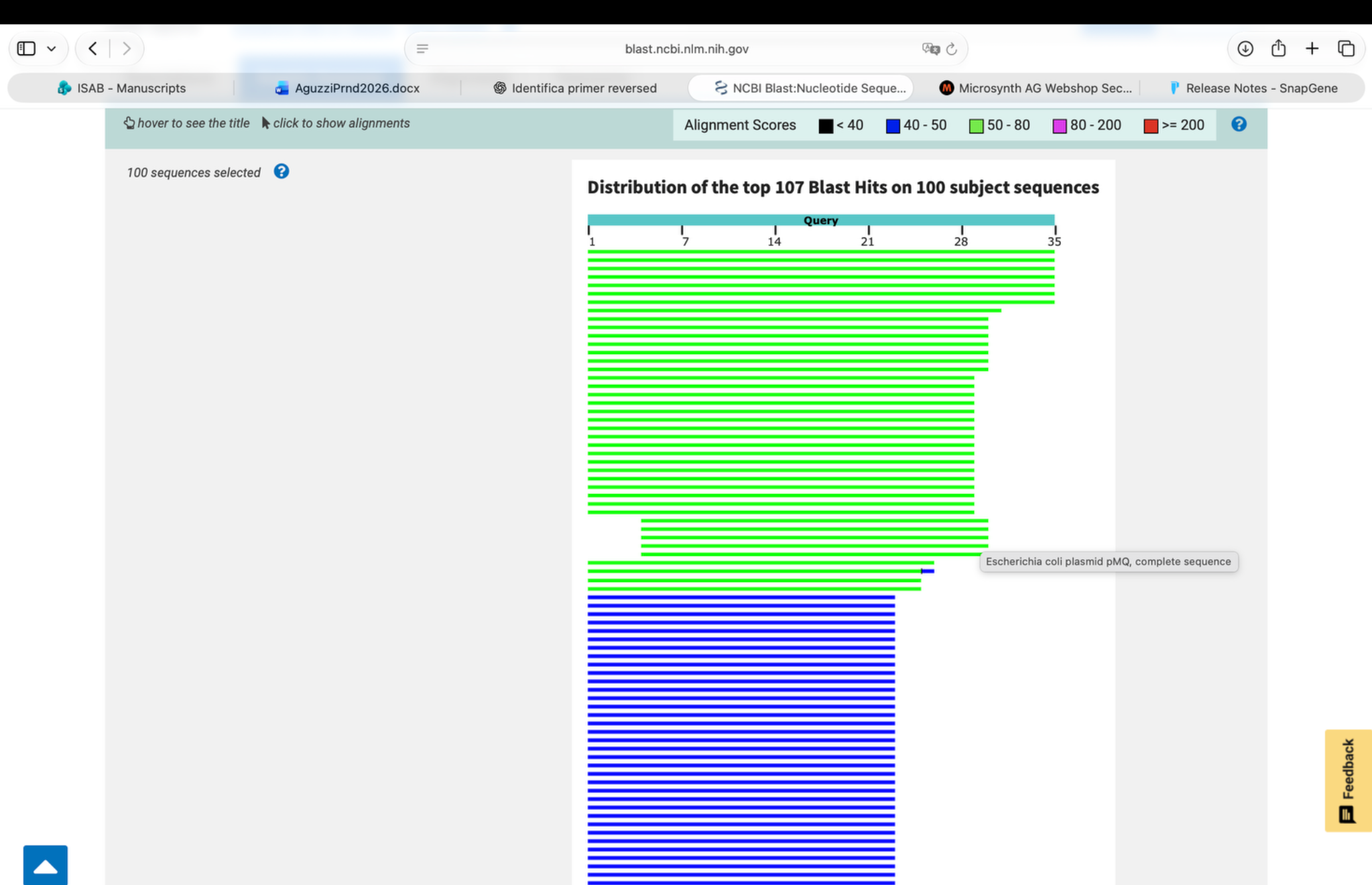

### image(20261108-185524).png

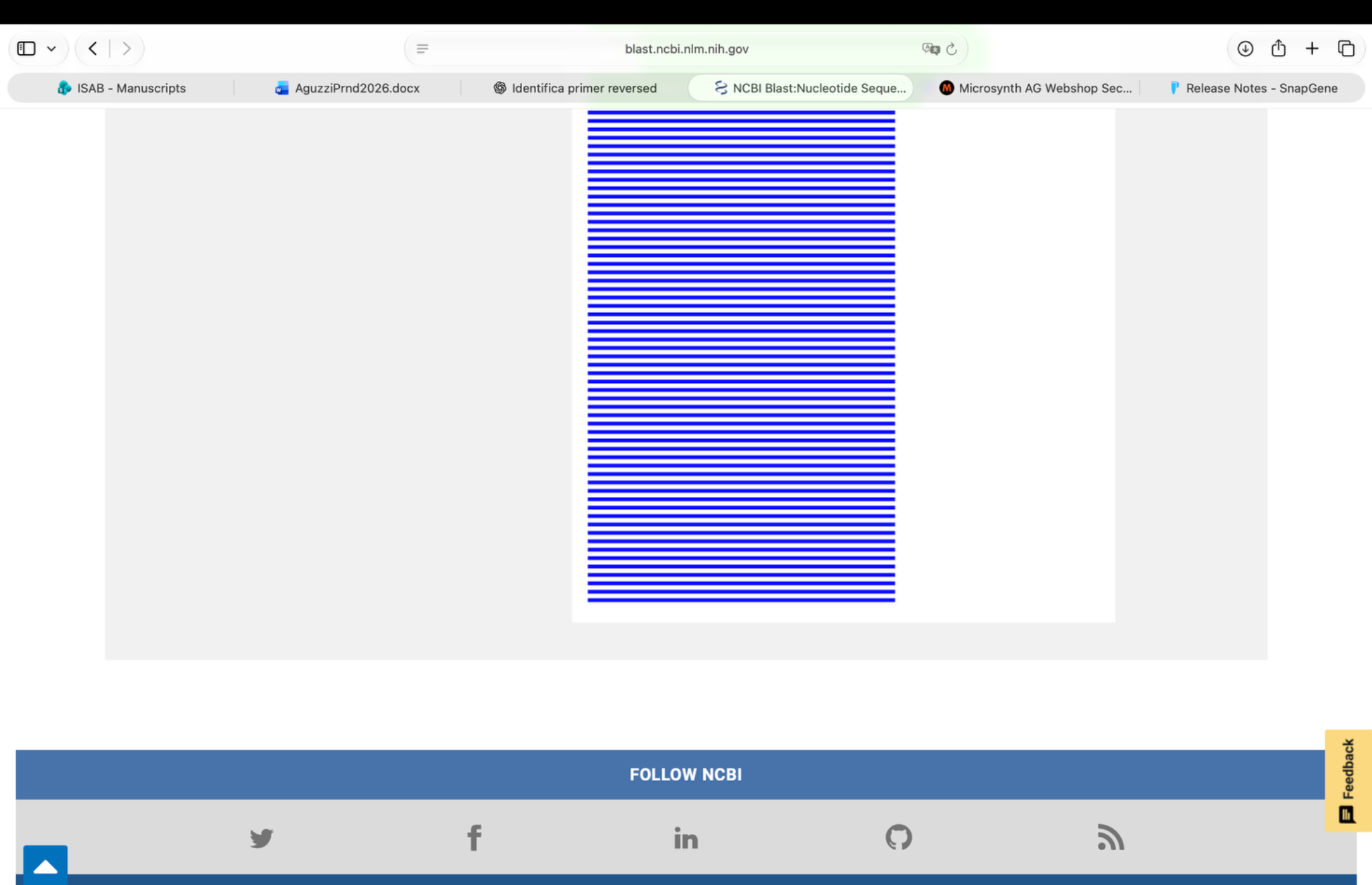

### image(20261108-185748).png

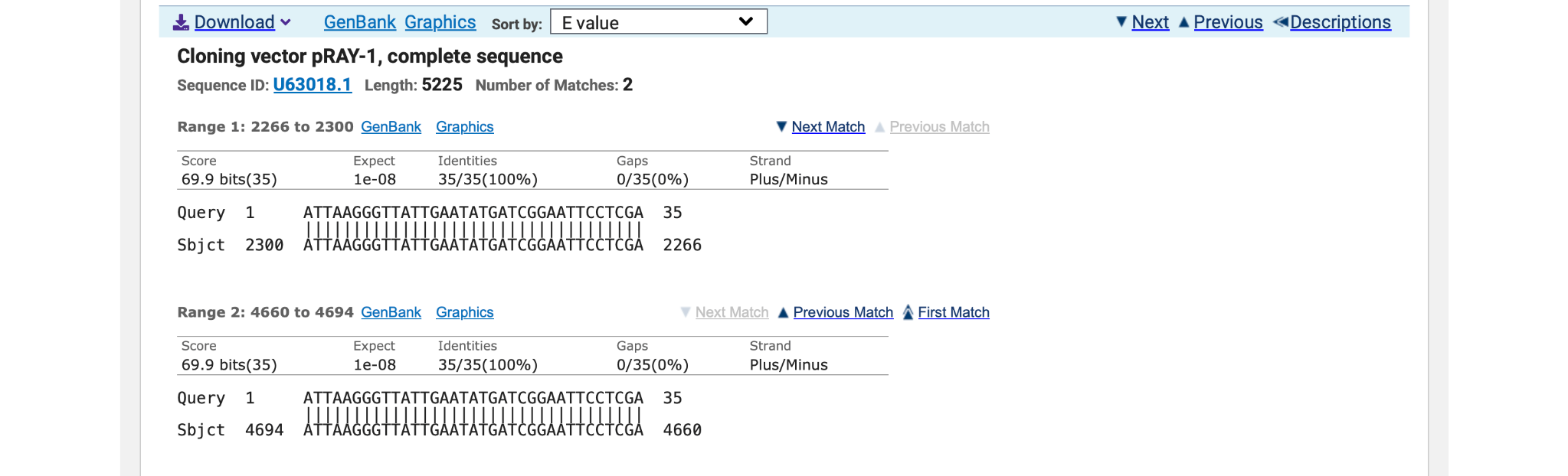

### image(20261108-190129).png

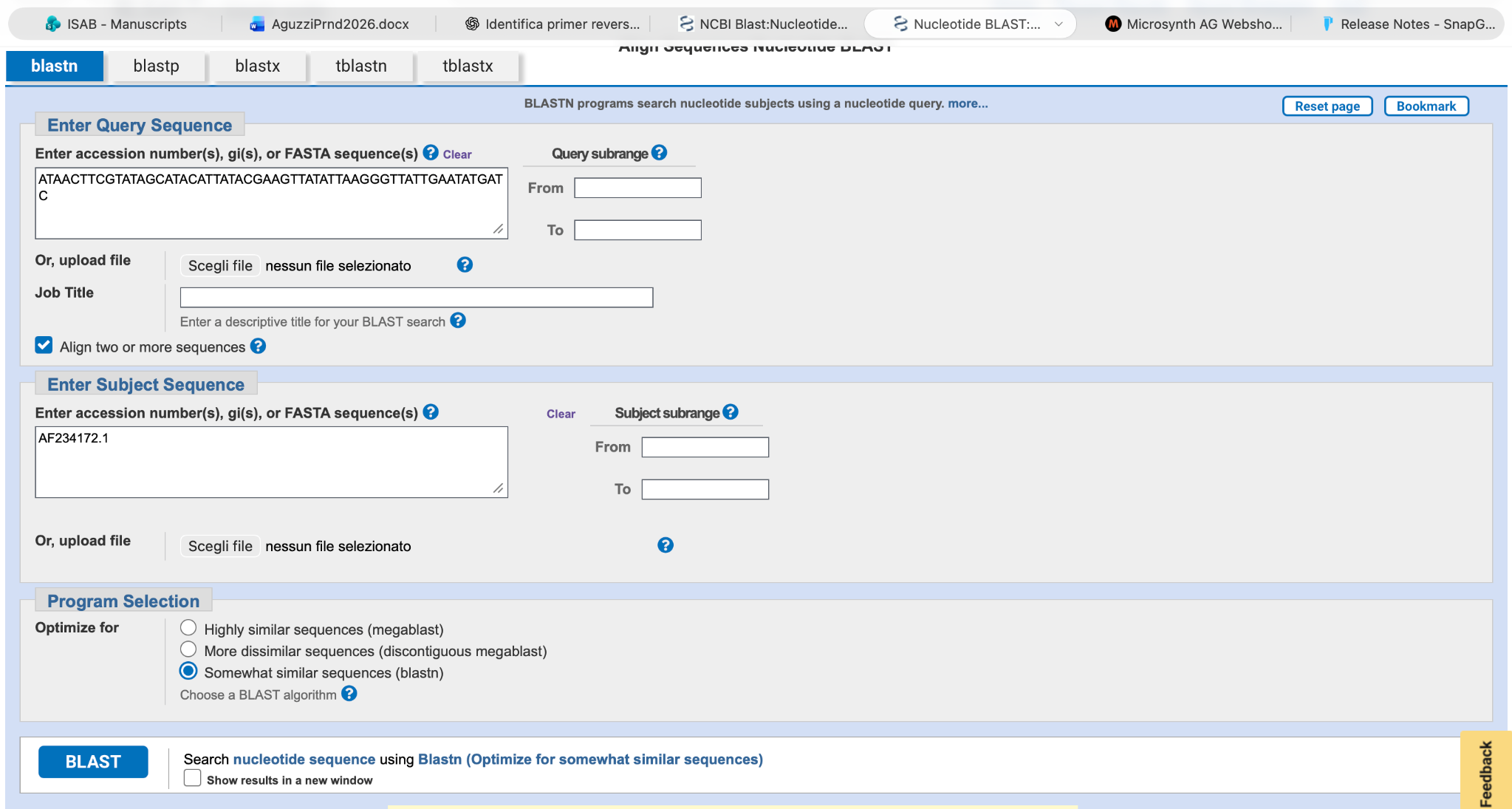

### image(20261108-190137).png

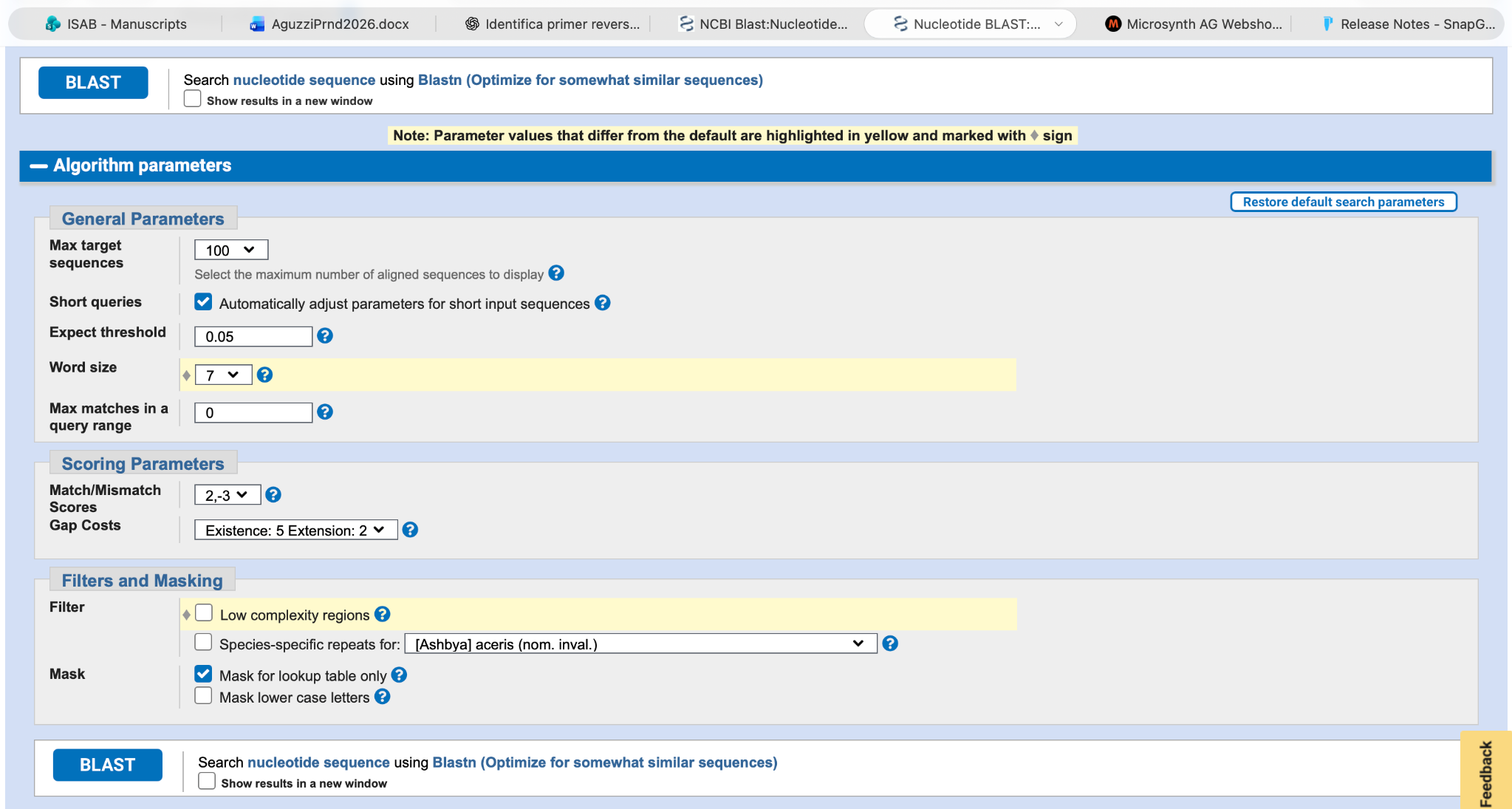

### image(20261108-190432).png

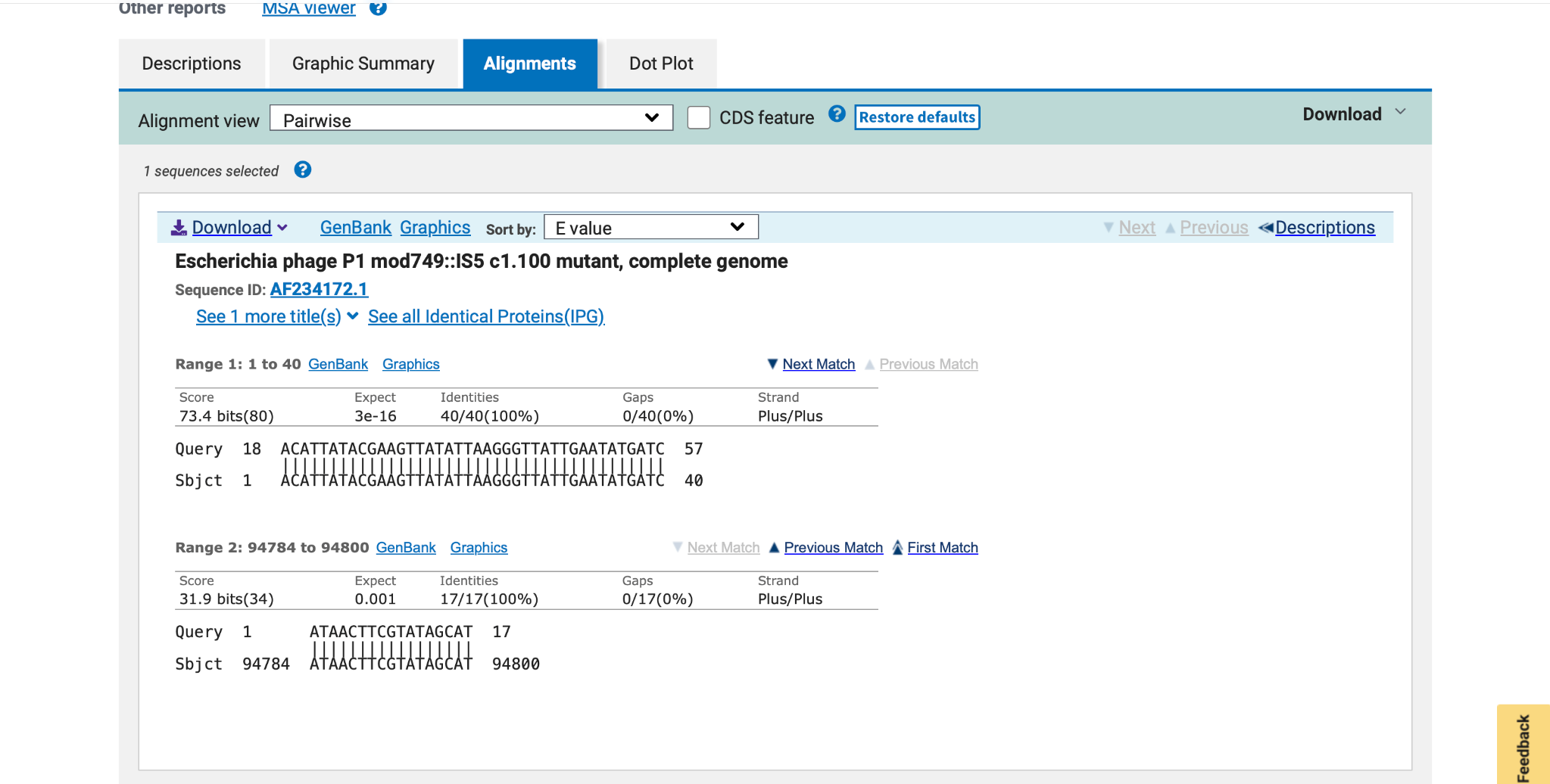

### image(20261108-190712).png

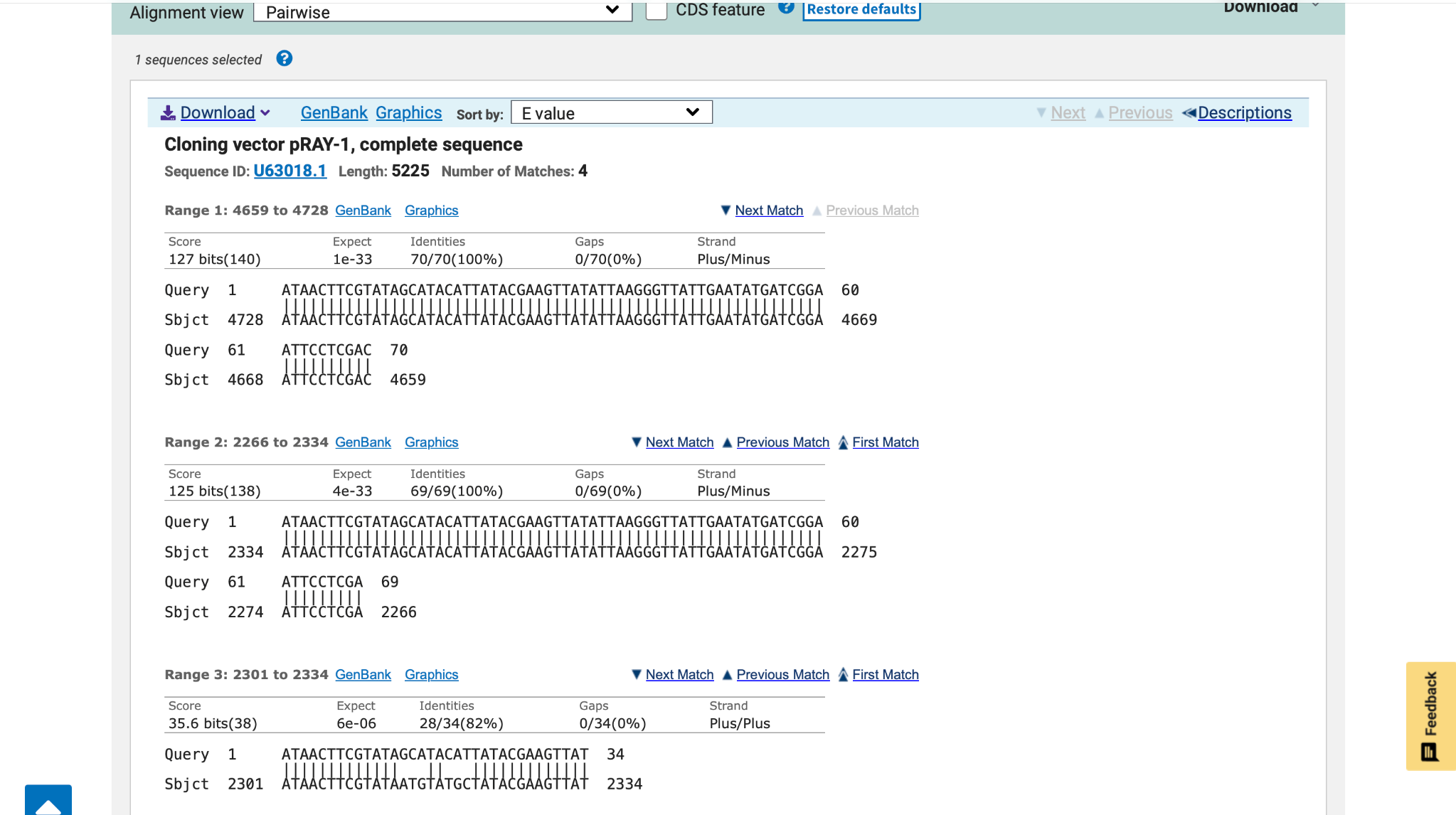

### image(20261108-190725).png

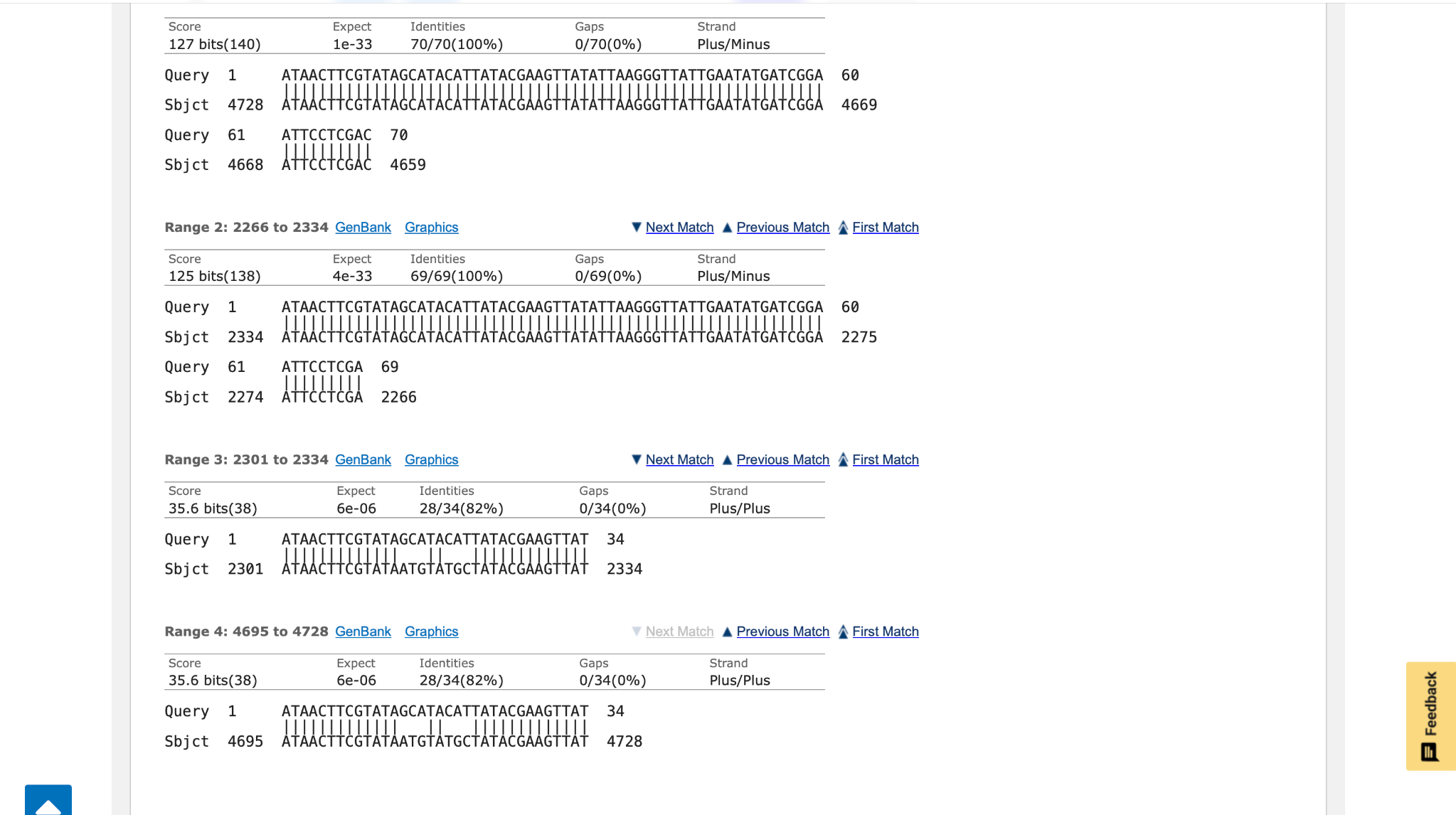

### image(20261108-191124).png

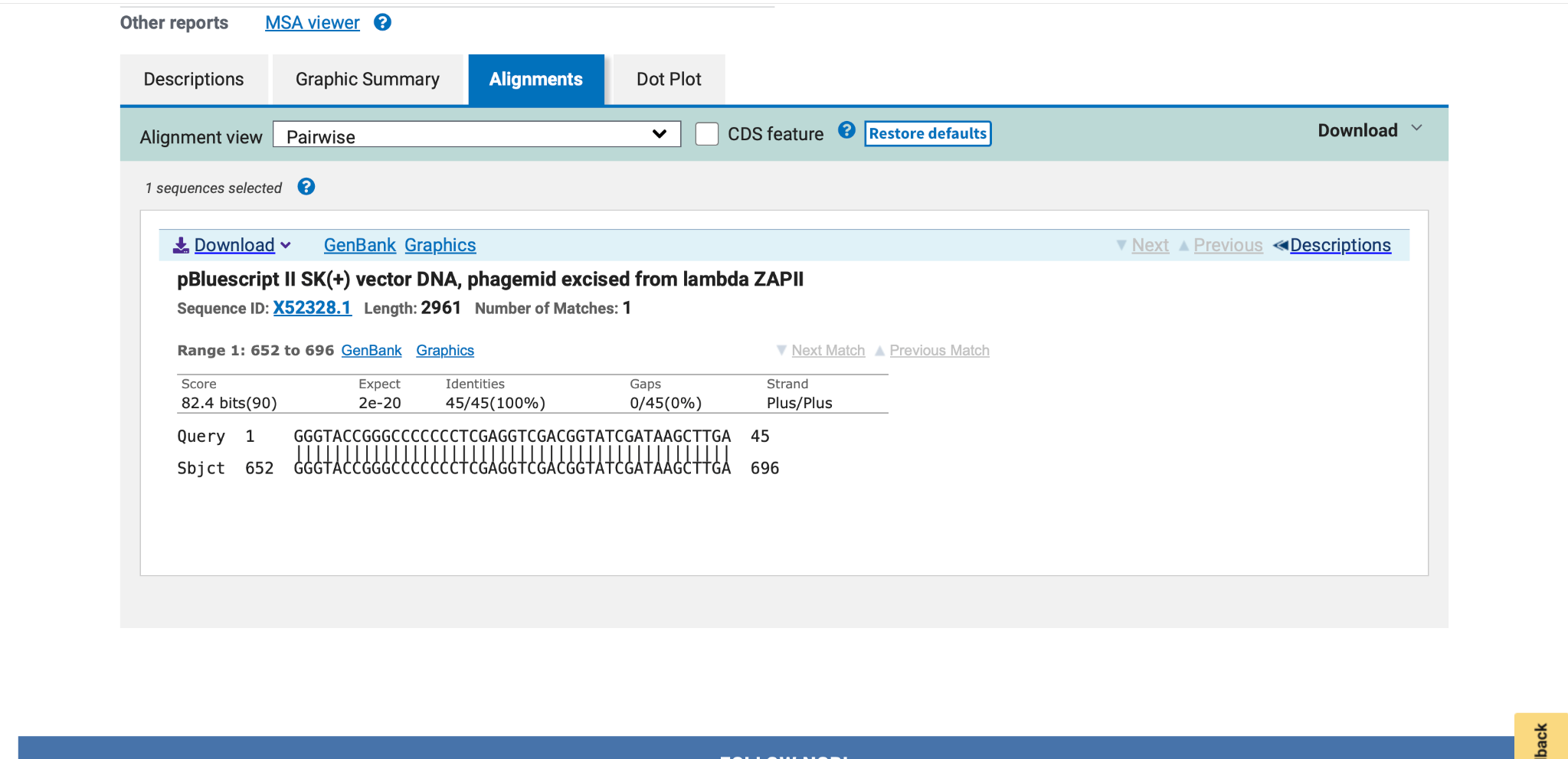

### image(20261108-191349).png

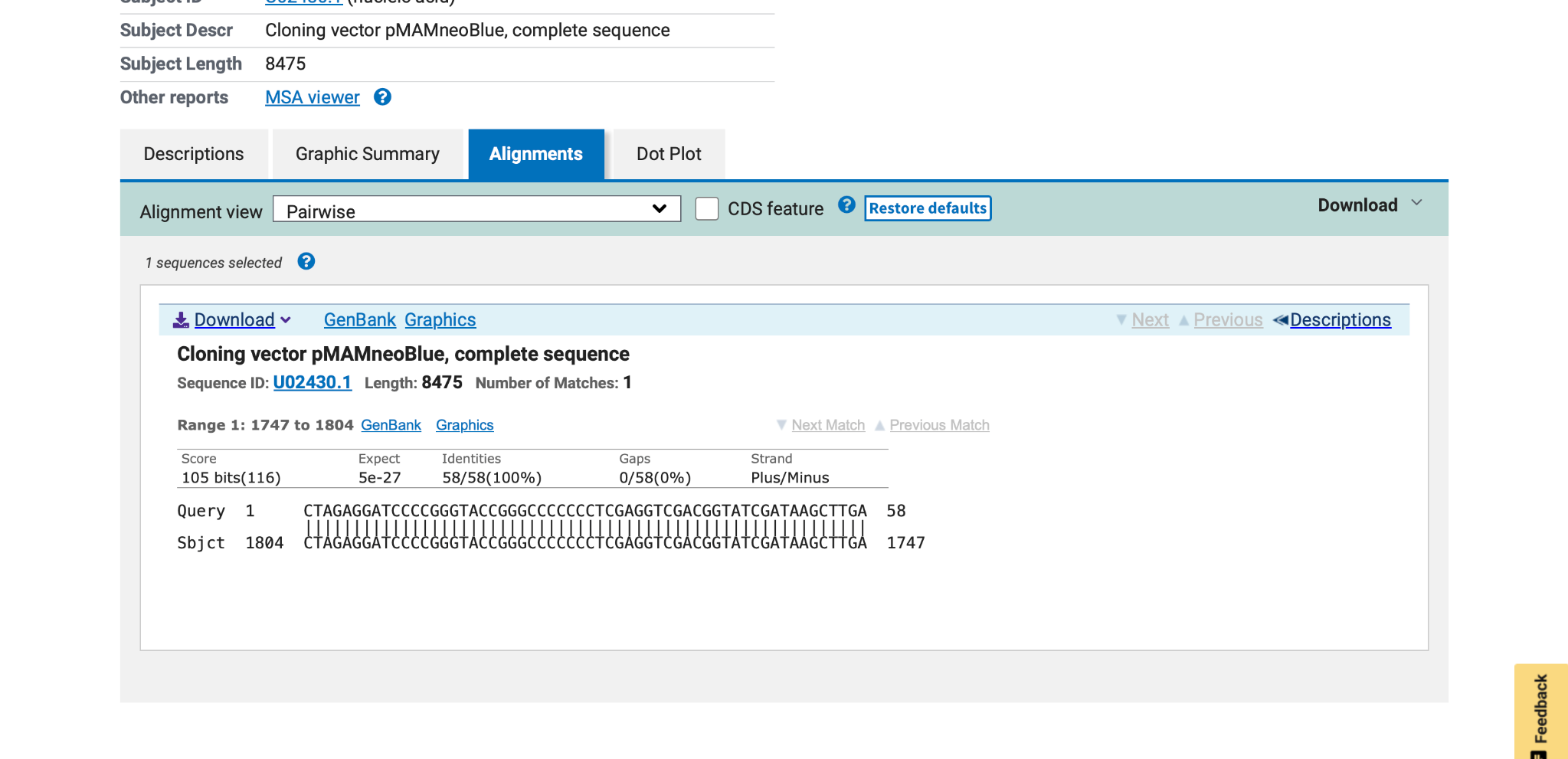

### image(20261108-191952).png

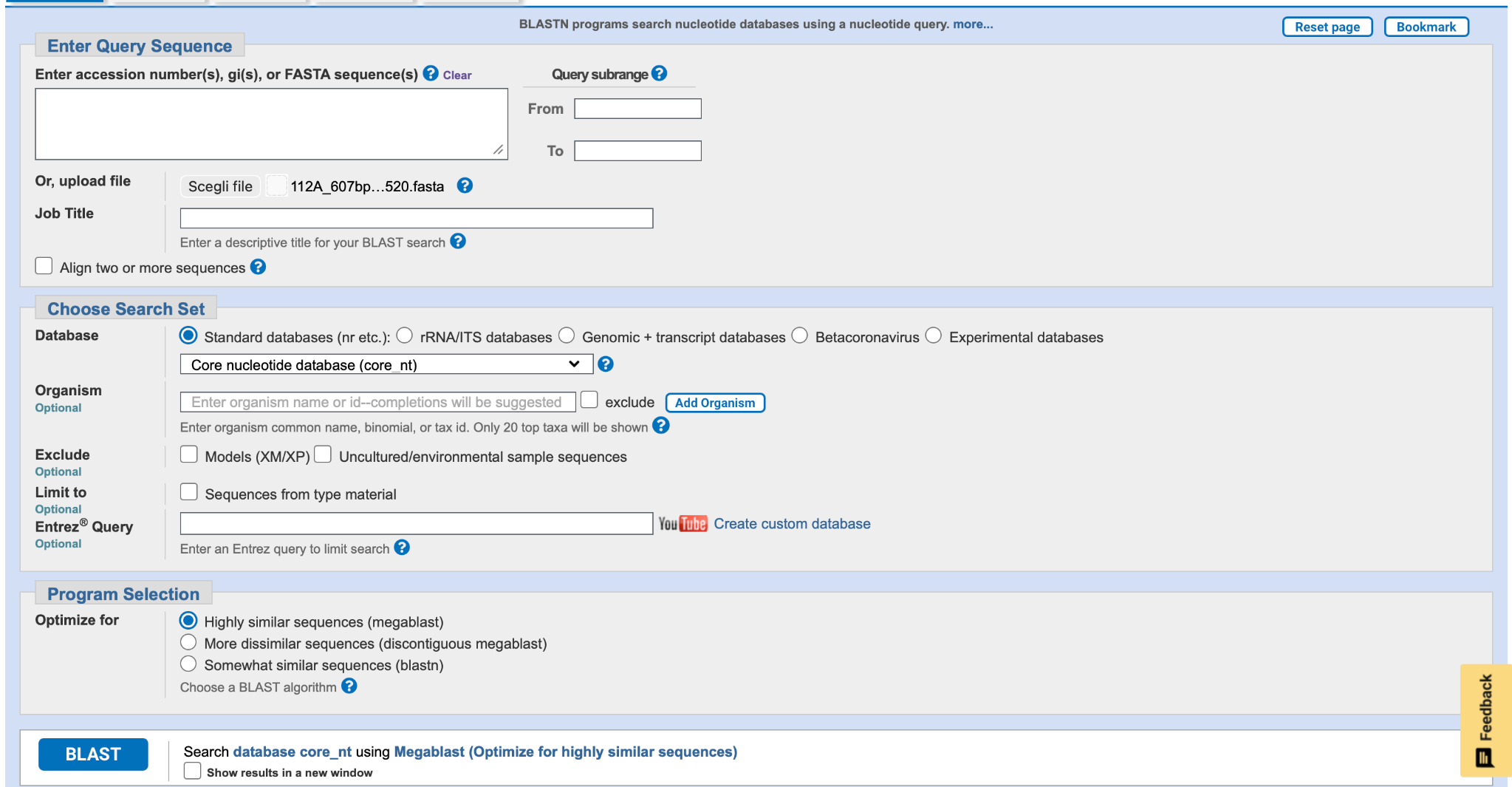

### image(20261108-192019).png

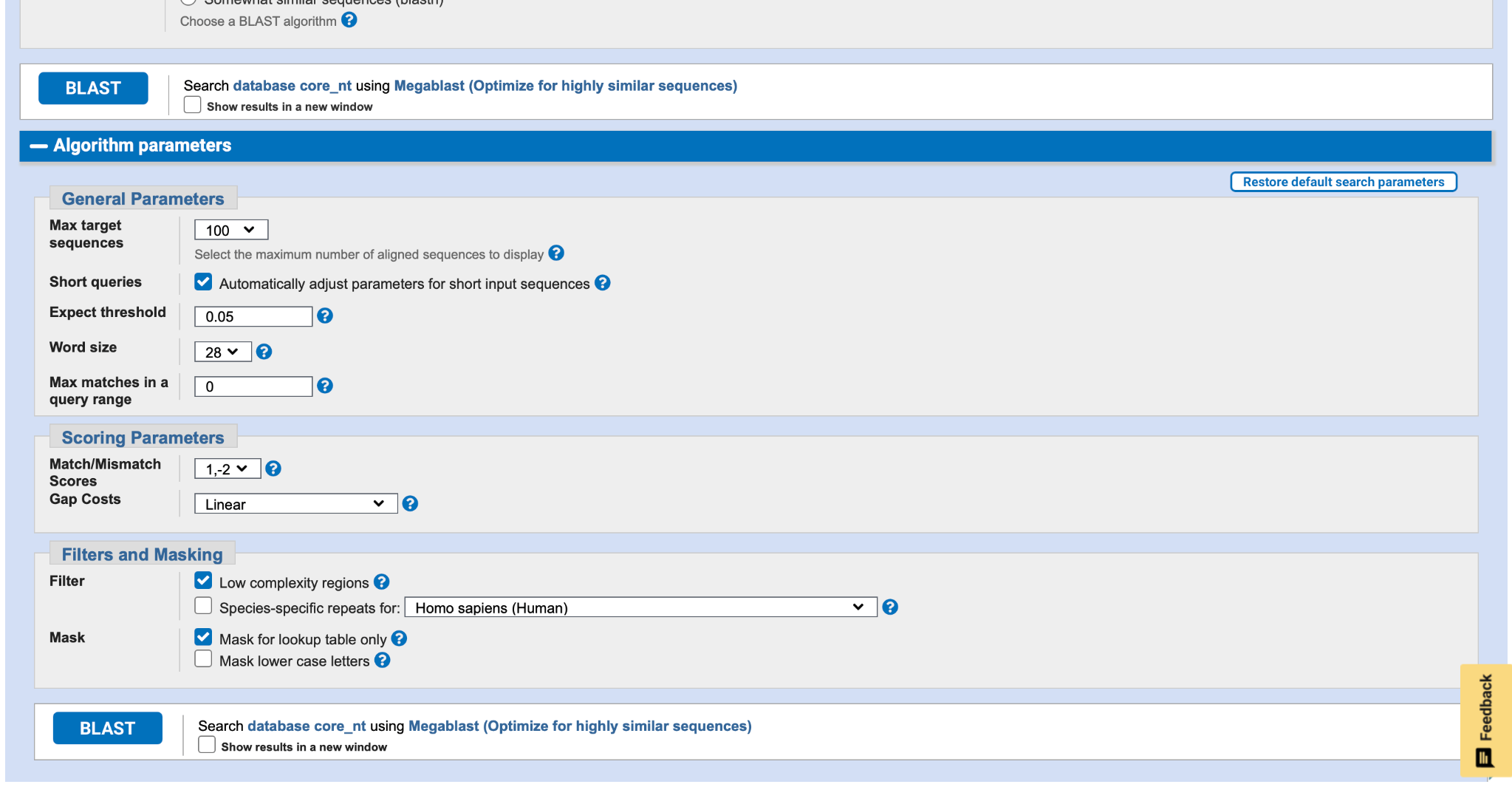

### image(20261108-192226).png

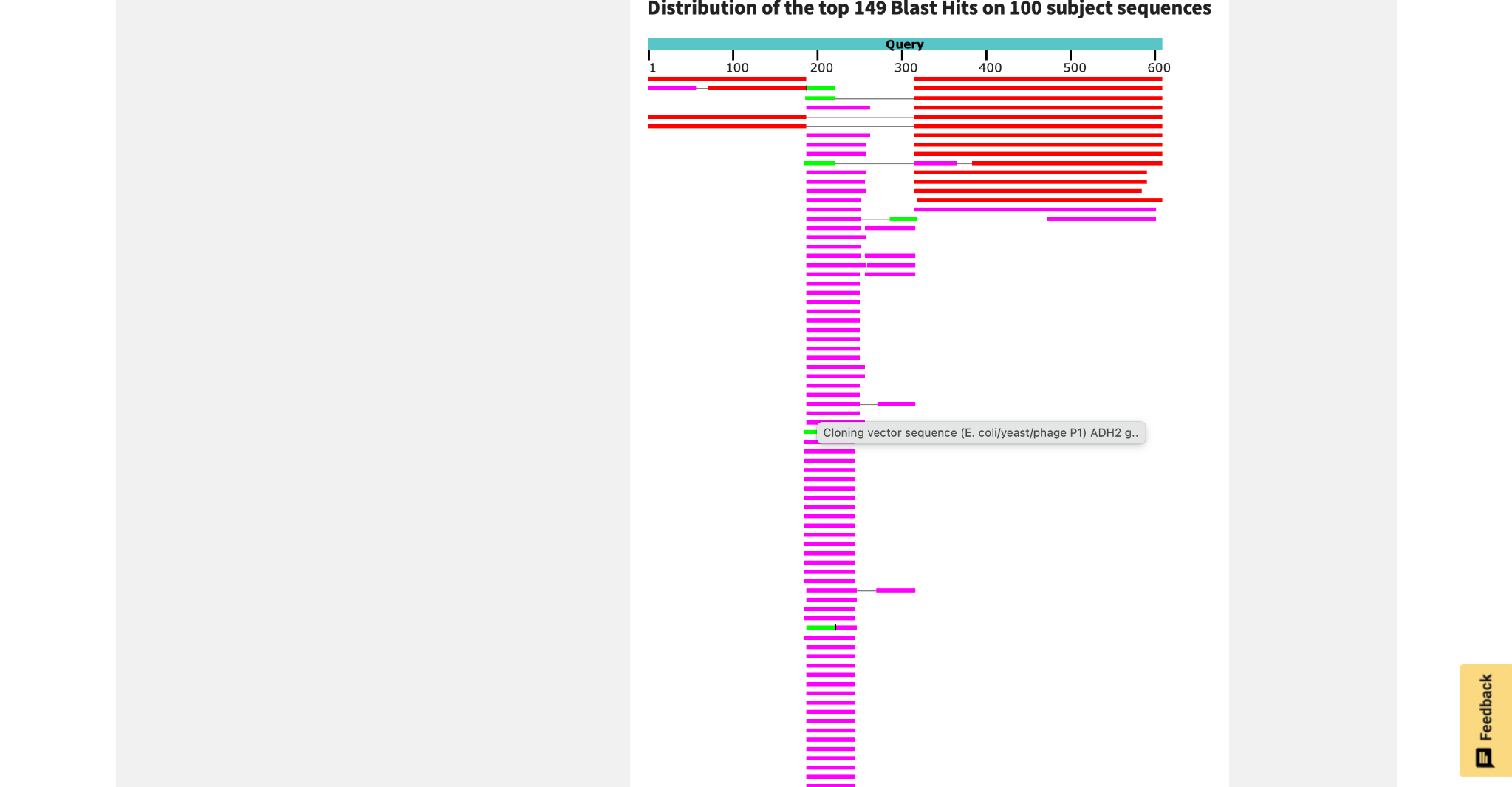

### image(20261108-192234).png

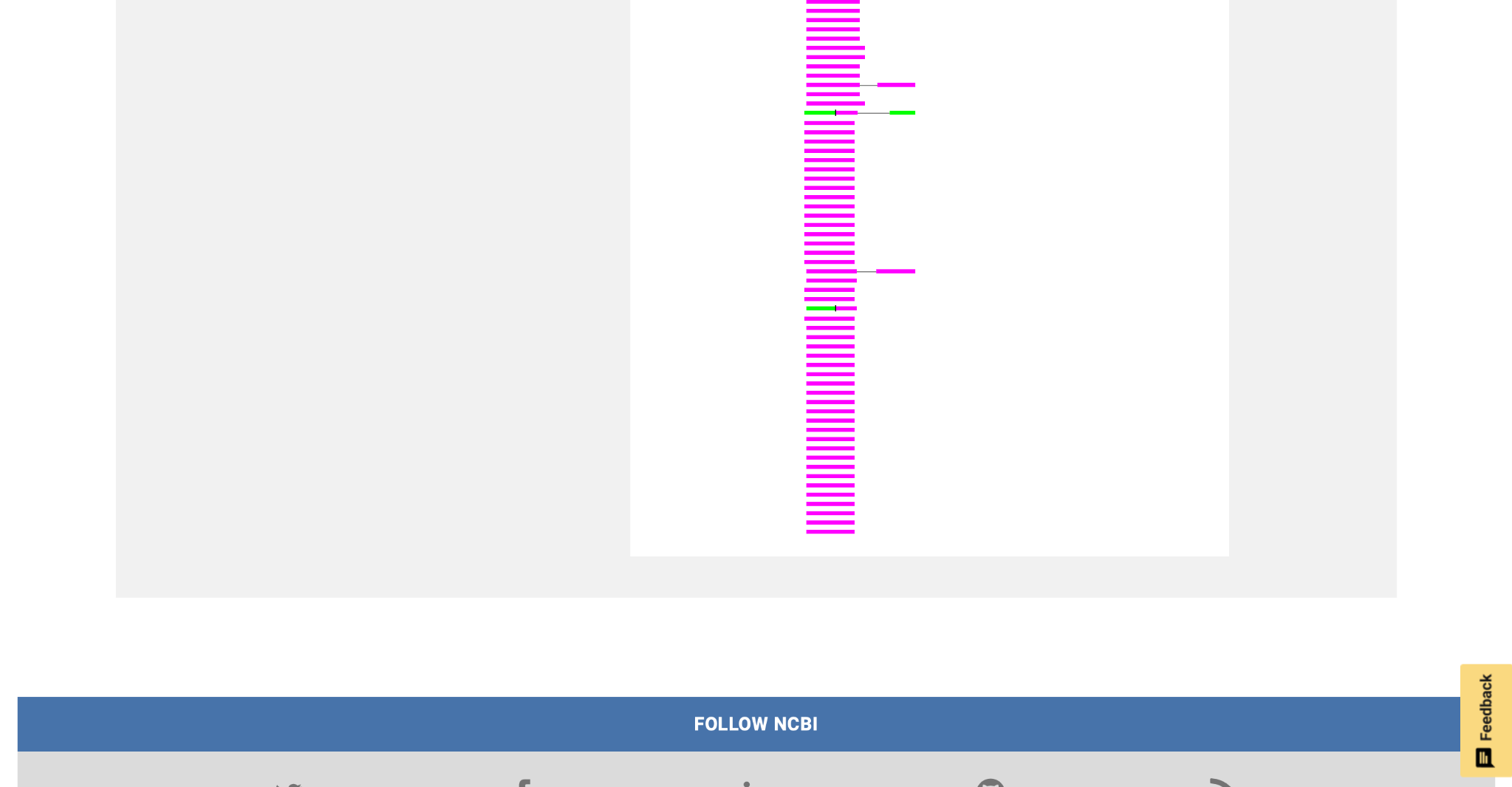

### image(20261108-192511).png

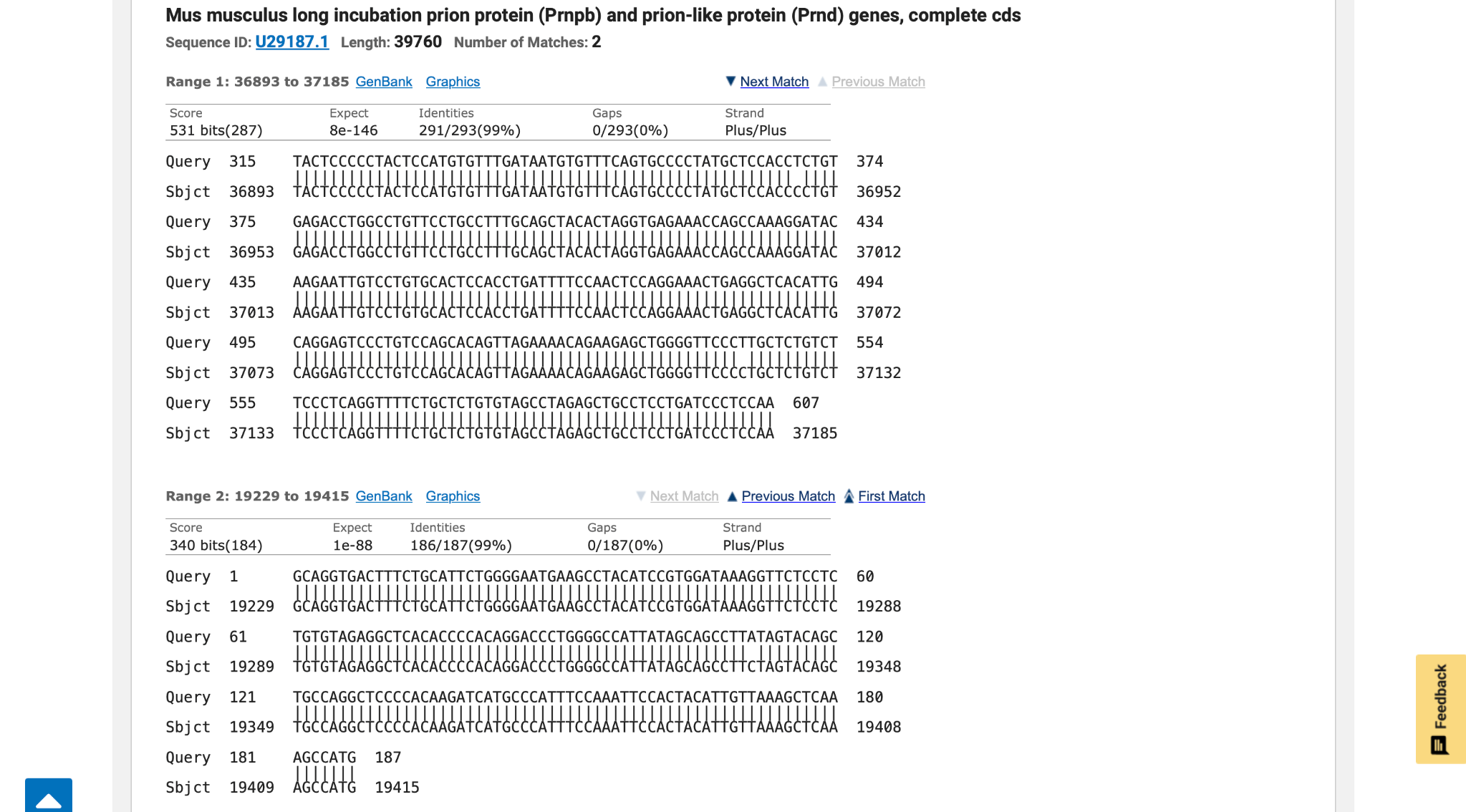

### image(20261110-135433).png

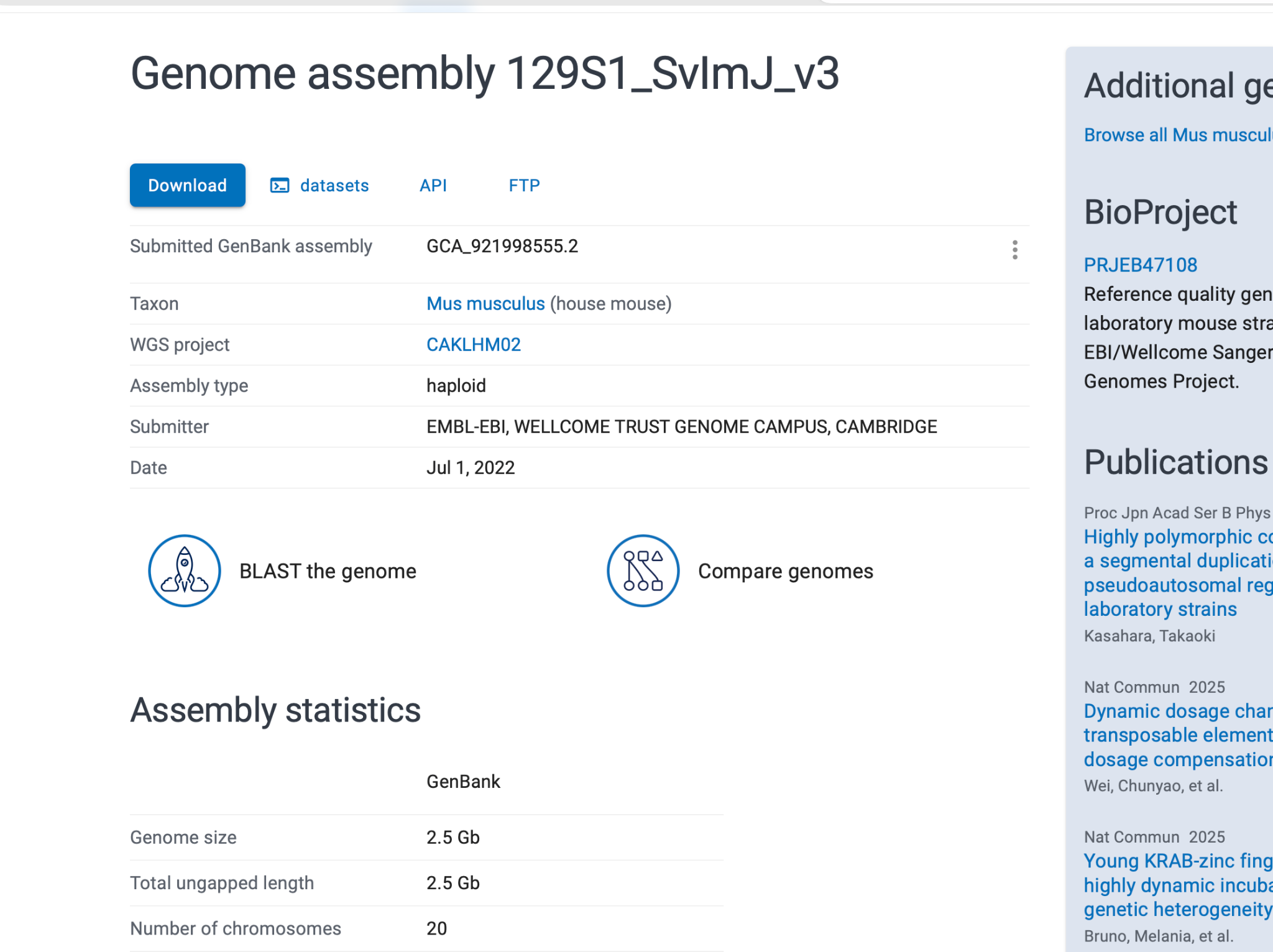

### image(20261110-135537).png

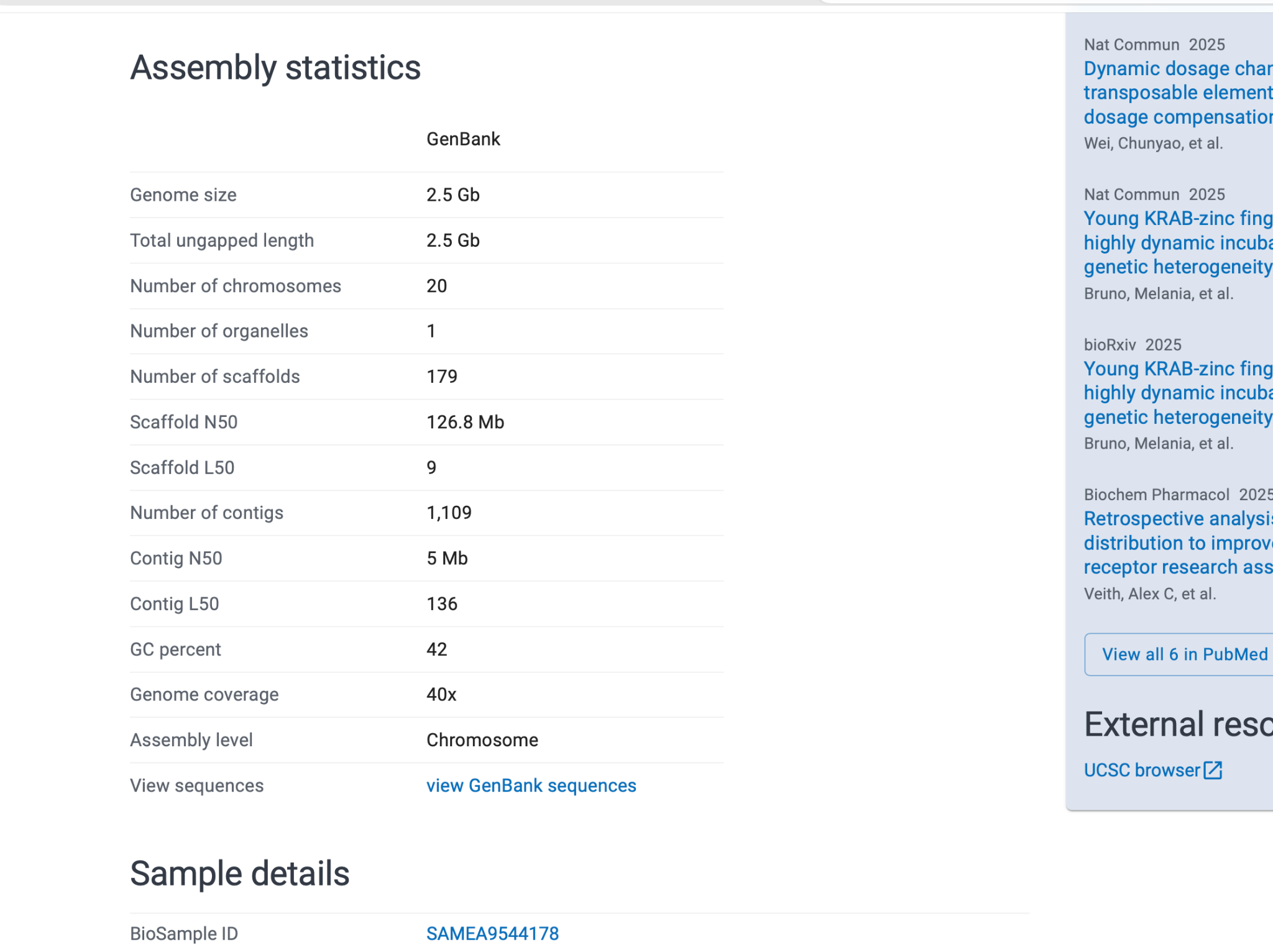

### image(20261110-135549).png

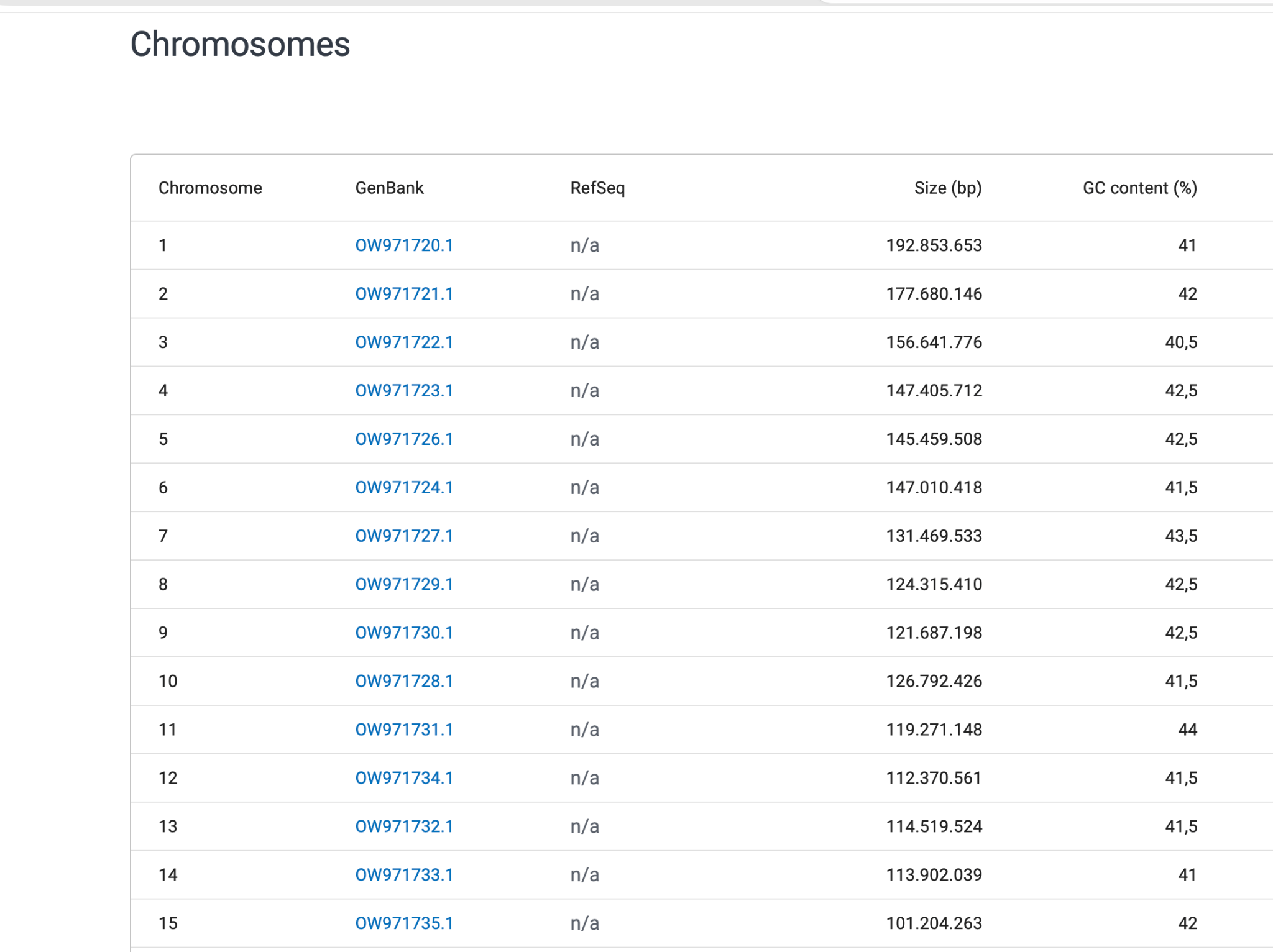

### image(20261110-141011).png

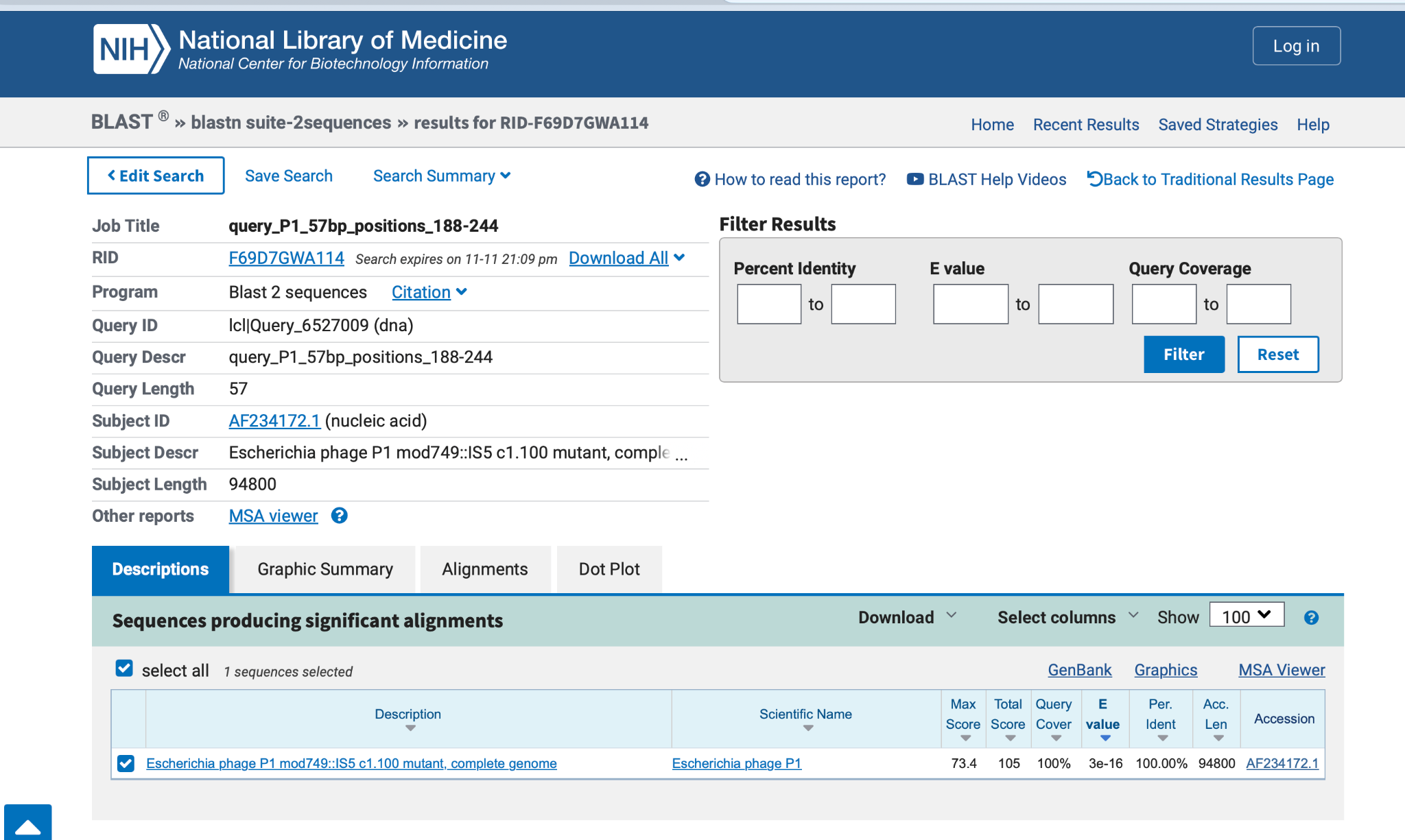

### image(20261110-141040).png

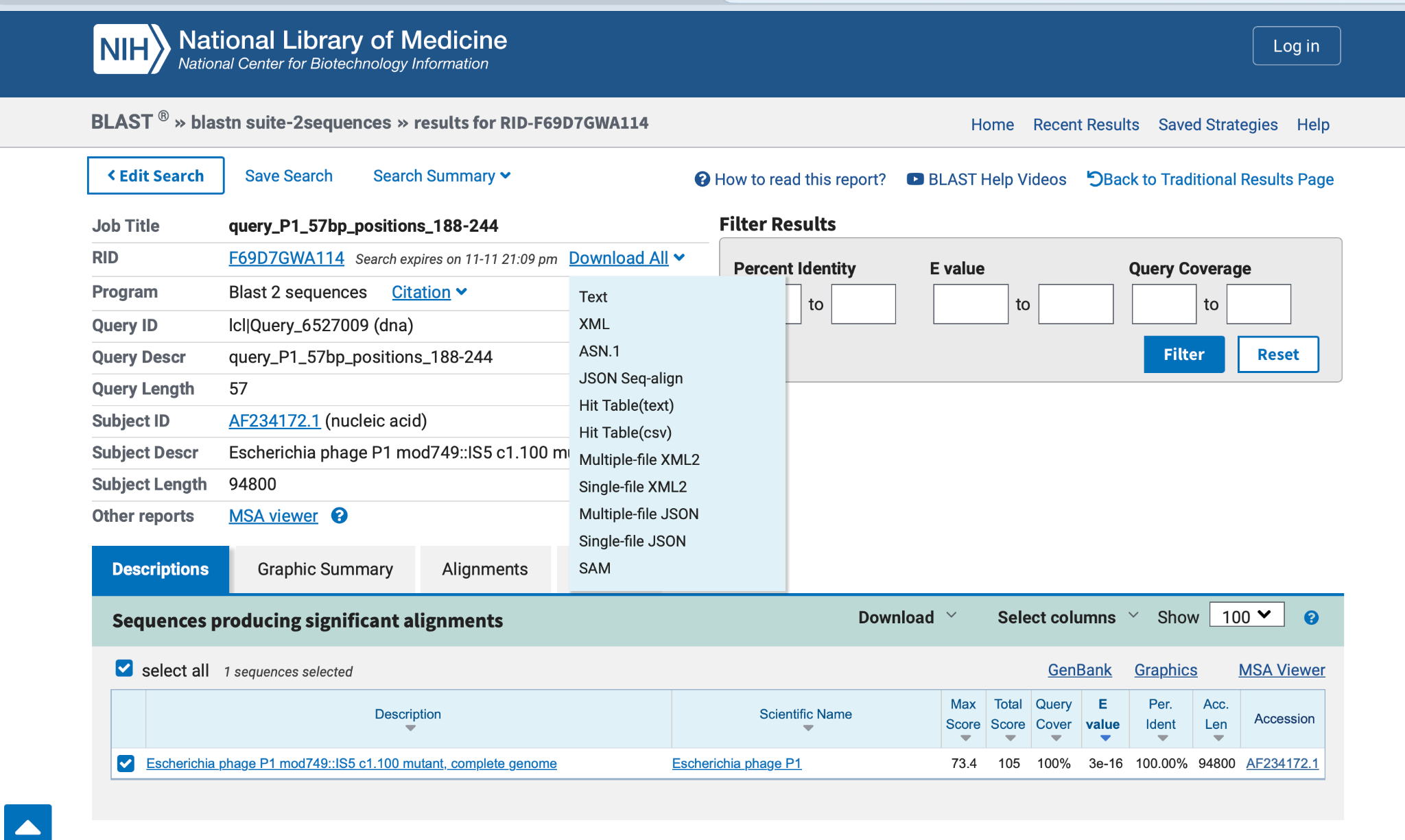

### image(20261110-143944).png

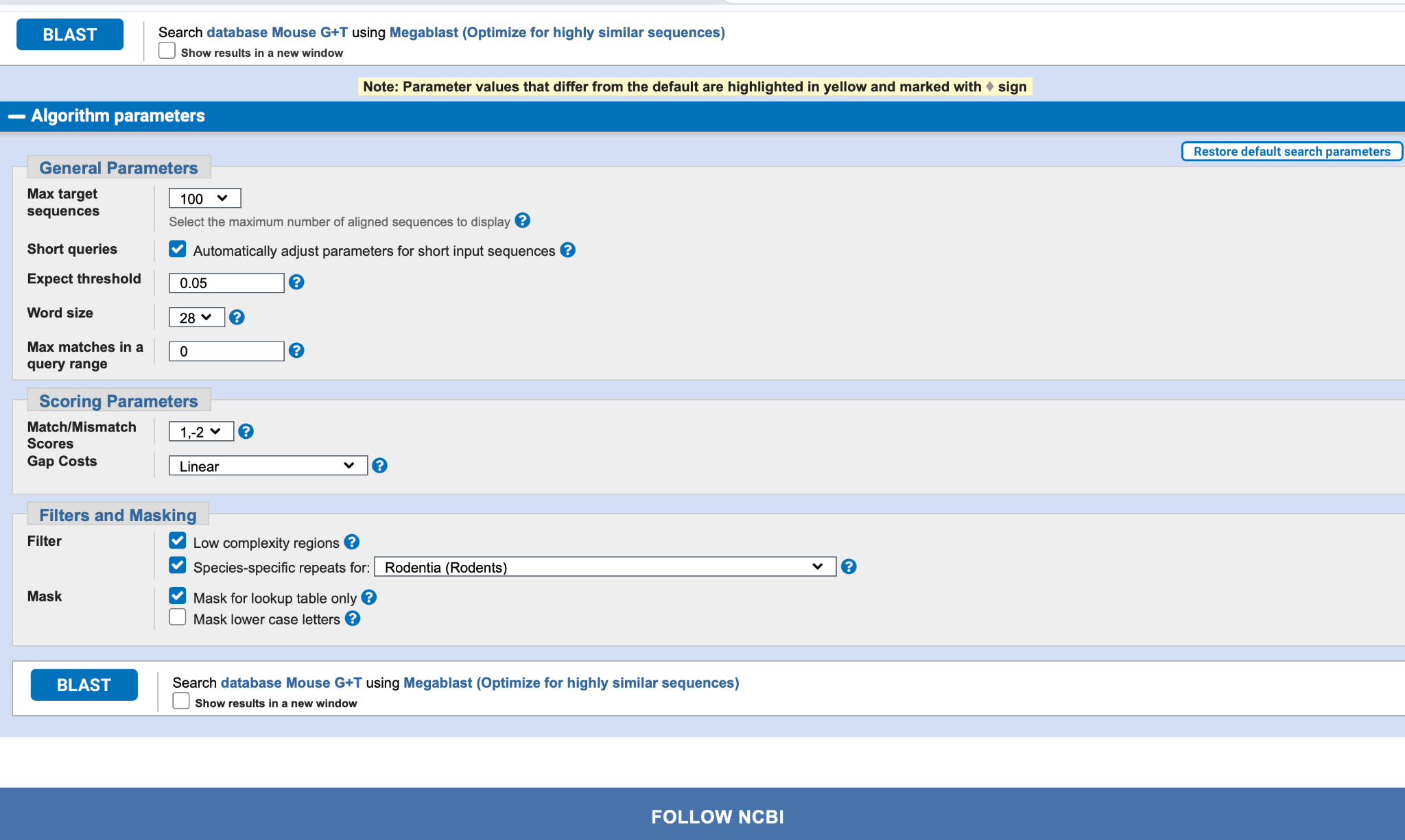
